# *Foxg1* Organizes Cephalic Ectoderm to Repress Mandibular Fate, Regulate Apoptosis, Generate Choanae, Elaborate the Auxiliary Eye and Pattern the Upper Jaw

**DOI:** 10.1101/2020.02.05.935189

**Authors:** Claudia Compagnucci, Michael J. Depew

## Abstract

Gnathostome jaw patterning involves focal instructive signals from the embryonic surface cephalic ectoderm (SCE) to a fungible population of cranial neural crest. The spatial refinement of these signals, particularly for those patterning the upper jaws, is not fully understood. We demonstrate that *Foxg1*, broadly expressed in the SCE overlying the upper jaw primordia, is required for both neurocranial and viscerocranial development, including the sensory capsules, neurocranial base, middle ear, and upper jaws. *Foxg1* controls upper jaw molecular identity and morphologic development by actively inhibiting the inappropriate acquisition of lower jaw molecular identity within the upper jaw primordia, and is necessary for the appropriate elaboration of the λ-junction, choanae, palate, vibrissae, rhinarium, upper lip and auxiliary eye. It regulates intra-epithelial cellular organization, gene expression, and the topography of apoptosis within the SCE. *Foxg1* integrates forebrain and skull development and genetically interacts with *Dlx5* to establish a single, rostral cranial midline.

## Introduction

The evolutionary success of vertebrates is intimately tied to developmental innovations affecting the head, including those leading to the elaboration of the brain, the emergence of the cranial neural crest (CNC) and ectodermal placodes, and the manifestation of jaws (1–3). As the vast majority of vertebrates are gnathostomes, and thus possess jaws, the true enormity of the observed radiation and diversification of vertebrates must be regarded in the particular light of the acquisition and modification of jaws (4–25). Jaws are prehensile oral apparatuses that can be further defined by their possession of two appositional units – the upper and lower arcades – that are articulated (hinged) and which largely arise during development from the first branchial arch (BA1) (5–12, 23–31).

Both the jaws and their BA1 primordia are characterized by their polarity: Within BA1, polarity is bidirectional and is roughly centered midway along the arch, extending from the so called ‘hinge’ proximally through the maxillary BA1 (mxBA1) and distally through the mandibular BA1 (mdBA1) (Figure 1 and Suppl. Figure 1; 5–9, 30–32). This directionality is subsequently mirrored by the associated structural polarity, centered at the functional jaw articulation, of the realized upper and lower jaw arcades. The fundamentality of this particular positional directionality for jaw development was initially molecularly revealed by the induced morphologic homeotic transformation of lower jaws into hinge-centric, mirror-imaged upper jaws in transgenic mice lacking a linked pair of *Distal-less* genes (*Dlx5/6*) whose expression is restricted to the lower jaw primordia (i.e., mdBA1) (6, 32, 33). Importantly, jaws exhibit additional levels of polarity: for instance, the upper jaws are overall more intimate with the neurocranium (the portion of the skull that houses the brain and primary sensory capsules) and its development, while the lower jaws are in closer developmental association with the emerging heart (5–11, 27). A functional manifestation of this additional polarity is the evolutionary inclusion of a premaxillary component derived from the olfactory placode-associated frontonasal prominences (FNP) in the upper jaw arcades of gnathostomes (except chondrychthyans; 4–31).

**Figure 1:**
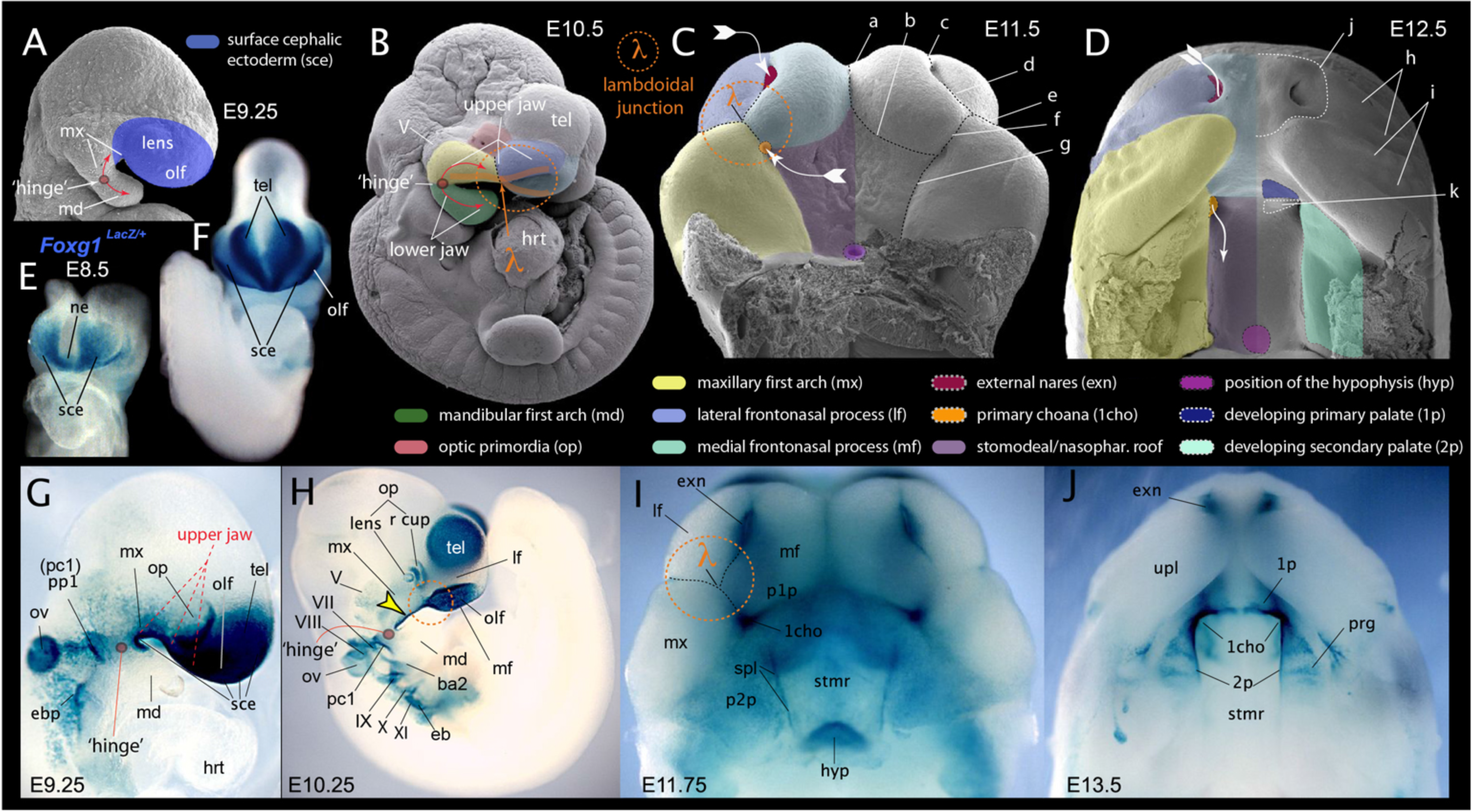
*Foxg1* is expressed in the SCE associated with epithelial morphogenesis and upper jaw development, including the λ-junction. <u>(**A-D**)</u> Scanning electron micrographs (SEM) of phenotypically wild type *Foxg1^+/−^* mouse embryos. <u>(**A**)</u> SEM of wild type embryo at E9.25 highlighting the area of the SCE connected with upper jaw development and the topography of polarity of the first branchial arch. ***md***, mandibular branch of the first branchial arch; ***mx***, maxillary branch of the first branchial arch. <u>(**B**)</u> SEM of an E10.5 mouse embryo indicating the λ−junction and its association with developing jaw polarity. tel, telencephalon. <u>(**C**)</u> SEM of E11.5 mouse embryo (with the lower jaw primordia removed) showing the embryonic primordia of the λ−junction and correlated anatomical boundaries. The converging white arrows indicate the invaginating external nares and internal (primary) choanae whose infoldings must unite and perforate to form a nasal cavity (fossa) that communicates directly with the lower respiratory system via a nasopharynx. ***a***, internasal line (frontal groove); ***b***, naso-stomodeal line; ***c***, dorsal intranasal line; ***d***, ventral intranasal line (nasal fin); ***e***, nasolacrimal line (groove); ***f***, oronasal groove (medial-choanal line); ***g***, stomodeo-palatal line. <u>(**D**)</u> SEM of E12.5 mouse embryo λ−junction correlated structures viewed (minus the lower jaw) from *norma basilaris externa*, including the primary and secondary palates, the internal choanae, and the external nares. ***h***, frontonasal mystachial pad; ***i***, maxillary mystachial pad; ***j***, regio rhinarica; ***k***, origin of caudal septum nasi. (**E-J**) β-galactosidase enzymatic activity in the SCE of *Foxg1^LacZ/+^* embryos. <u>(**E**)</u> *Norma frontalis*, E8.5. <u>(**F**)</u> *Norma frontalis*, E9.25. <u>(**G**)</u> *Norma lateralis*, E9.25. Compare with ‘A’. <u>(**H**)</u> *Norma lateralis*, E10.25. Compare with ‘B’. <u>(**I**)</u> *Norma basilaris externa*, E11.75. Compare with ‘C’. <u>(**J**)</u> *Norma basilaris externa*, E13.5. Compare with ‘D’. **<u>Abbreviations</u>**: ***ba2***, second branchial arch; ***eb***, epibranchial cells; ***exn***, external nares; ***hrt***, heart; ***hyp***, hypophysis; ***lf***, lateral frontonasal process; ***md***, mandibular branch of the first branchial arch; ***mf***, medial frontonasal process; ***mx***, maxillary branch of the first branchial arch; ***ne***, neuroectoderm; ***olf***, olfactory placode or pit; ***op***, optic primordia; ***ov***, otic vesicle; ***pc1***; first pharyngeal cleft; ***p1p***, primordia of primary palate; ***p2p***, primordia of secondary palate; ***ppl***, first pharyngeal plate; ***prg***, ruggae primordia; ***r cup***, rostral (nasal) optic cup; ***sce***, surface cephalic ectoderm; ***spl***, stomodeo-palatal line; ***stmr***, stomodeal roof; ***tel***, telencephalon; ***V***, Nervus trigeminus; ***VII***, Nervus facialis; ***VIII***, Nervus vestibuloaccousticus; ***IX***, Nervus glossopharyngeus; ***X***, Nervus vagus; ***XI***, Nervus accessorius; ***1cho***, primary choanal pit; ***2p***, secondary palatal shelf; **λ**, center of the lambdoidal junction.

The developmental system that establishes these polarities and patterns the developing jaws involves an intricate spatiotemporal series of reciprocal inductive and responsive interactions between the cephalic epithelia (both ectodermal and endodermal) and the subjacent CNC mesenchyme (6–10, 25, 30–50; **reviewed in 9**). Notably, augmentations of the developmental mechanisms and dynamics specifically regulating the cephalic epithelia and mesenchyme of the nascent upper jaw primordia (i.e., mxBA1 and FNP) and their associated structures have been connected with a number of significant gnathostome evolutionary events (8–22, 51–63), including: the teleostean radiation (impelled in part by making the maxillary–premaxillary– neurocranial connectivity less rigid and more mobile), the colonization of land by tetrapods (being in part enabled by acquisition of internal choanae thereby enabling respiration with a closed mouth), both the mammalian ability to masticate while breathing (enabled by the presence of a secondary palate) and the possession in mammals of highly expanded and coordinated olfaction and respiration (propelled by the evolution and elaboration of often labyrinthine turbinal bones in the nasal cavity and paranasal sinuses in the associated cranial skeleton), the cetacean radiation, as well as the primary split of primates into strepsirhini or haplorhini (following the truncation of the nasal cavity, the loss of the rhinarium [the specialized glabrous skin surrounding the external nares] and the acquisition of mobile upper lips in haplorhines).

Each of these significant evolutionary transitions involved modulation of embryonic developmental events associated with the lambdoidal junction (λ-junction), a transient embryonic structure so named for the shape formed at the confluence of the mxBA1 and the medial (mFNP) and lateral (lFNP) FNP (Figure 1; 8,9,64). The organization and morphogenesis of the λ-junction and its derivatives is complex and embodies the future positions of the nose (including the nares, nasal (olfactory) capsules (NC), and choanae), the upper lips, the premaxillae and their incisors, the maxillae and their molars, the primary and secondary palates, as well as the optic capsules (OPC) and the orbito-nasal (e.g., nasolacrimal) ductal system (Figure 1). Highlighting this complexity in humans, two of the most common and disfiguring congenital malformations – cleft lip and cleft palate – are associated with disrupted development and elaboration of the λ-junction; so too are less common, though no less significant, malformations such as choanal atresia and arhinia (65–76).

While substantial progress has been made in understanding how the epithelium of BA1 signals to the underlying CNC mesenchyme, in particular within mdBA1, much less is clear regarding how the epithelium of the surface cephalic ectoderm (SCE) associated with the λ-junction is organized and genetically regulated ontogenetically to participate in the development and evolution of the upper jaws and associated structures. In an effort to further clarify the relationship between the elaboration of the λ-junction and the development and evolution of the upper jaws we have been investigating the ontogeny of developmental influences on the λ-junction. We have been particularly interested in functional investigations of genes that are ectodermally expressed early in the gnathostome embryo both in the SCE that eventually overlies the λ-junction (and which gives rise to the olfactory placode) and in the early neuroectoderm (NE): Such genes, being expressed on either side of the patterning influence of the anterior neural ridge (ANR), are ideally expressed to play roles in the integration of the upper jaw with the neurocranium as well as the functional integration of the neurocranium and the brain. Among these genes are those encoding the master regulatory transcription factors *Dlx5*, *Pax6*, and *Foxg1* (originally *BF-1* for *Brain Factor-1)*. *Foxg1* is an evolutionarily conserved member of the Fox/Forkhead family of winged-helix transcription factors, and, while both *Dlx5* and *Pax6* have been extensively examined for their roles in both neural and craniofacial development in mice (6, 77–86), *Foxg1* has principally been investigated for its role in forebrain development and corticogenesis (87–130).

Herein we utilize previously generated loss-of-function transgenic mice (130) to demonstrate that *Foxg1*, which is expressed in the SCE and NE but not in the CNC is required for proper upper jaw development as well as the development of each of the primary sensory capsules (otic, optic and olfactory) of the neurocranium, and the calvarium, and we add to and extend previous observations regarding the necessity for *Foxg1* in eye and ear development. We present evidence that *Foxg1^−/−^* mutants do not develop internal choanae (leading to choanal atresia), that their palatal shelves do not descend and fully elongate vertically and are partially cleft, and that a nasopharyngeal duct is absent and thus a functional nasal respiratory channel fails to form. Moreover, we show that *Foxg1* regulates intra-epithelial organization and the cellular dynamics of the SCE, including regulating patterns of gene expression and the topography of apoptotic cell death within the SCE and proximal BA1 epithelia. Notably, we further show that *Foxg1* acts to suppress the induction of lower jaw identity in the upper jaw primordia as, without *Foxg1*, mxBA1 partially acquires a deep mdBA1 molecular identity. To initiate a larger picture of the hierarchy of genetic interactions governing the development of the upper jaws and rostral neurocranium, we additionally demonstrate that *Foxg1* and *Dlx5* genetically interact in the elaboration of the SCE as, for instance, compound *Foxg1^−/−^*; *Dlx5^−/−^* embryos fail to generate a λ-junction and three (rather than one) nasal ‘midlines’ develop. Finally, our demonstration of the range, nature, and significance of the non-neural craniofacial deficits exhibited in *Foxg1^−/−^* mutants mandates that its roles outside of neurogenic control must be appreciated when evaluating the function of *Foxg1* during development and evolution. As the functional and integrative interactions between the SCE, the NE, and the intervening CNC are increasing being elucidated and adduced, and as the significant roles we have uncovered for *Foxg1* in cephalic development outside of neurogenic control are recorded, the work presented herein further presents a clear caveat to both past and future investigations ascribing biological functions when utilizing the endogenous *Foxg1* as a ‘*Cre*-deleter’ in genetic studies of cranial structures, including the central nervous system and the craniofacial complex.

## Results

### *Foxg1* is expressed in both the NE and SCE, including the epithelium of the *λ*-junction and its derivatives

To confirm that the ontogeny of *Foxg1* expression includes the SCE (and thereafter the λ-junction), we took advantage of the *Foxg1* mutant allele’s in-frame *LacZ* cassette (93, 100, 106, 130). We detected β-galactosidase enzymatic activity in the SCE of *Foxg1^LacZ/+^* (hereafter simply *Foxg1^+/−^*) embryos beginning at embryonic day 8 (E8) (Figure 1; Supplementary Figures 1, 2). As has been previously reported, we found positive cells throughout the nascent SCE epithelium as well as in the NE on the opposite side of the ANR. Activity remained intense throughout the SCE through E9.25, including in the epithelial cells of the developing otic, optic (lens), olfactory, and epibranchial placodes. The telencephalic NE likewise exhibited highly positive staining.

Notably, in addition to regions of *LacZ* expression previously characterized, we found that the oral ectoderm of mxBA1 was heavily stained in *Foxg1^+/−^* embryos, as was the proximal first pharyngeal plate (where the SCE meets the endodermal epithelia of the first pharyngeal groove); however, the ectoderm distally covering mdBA1 showed little sign of β-galactosidase activity. Moreover, the stomodeal epithelium underlying the developing forebrain also stained highly positive, as did the hypophyseal placode and the rostral foregut endoderm. We found that this pattern of expression was replicated by whole mount *in situ* hybridization for *Foxg1* at comparable stages (Supplementary Figure 1).

From E9.5 onward the widespread SCE *Foxg1^LacZ^* expression characteristic of earlier embryos began to focalize such that by E10.25 the highest levels of β-galactosidase activity detected were within the epithelia of the invaginating olfactory pits, the rostral (nasal) lens, Rathke’s pouch, the already invaginated otic vesicle, the nasal side of the optic cup and forming stalk, and the developing telencephalic vesicles, (Figure 1). While *Foxg1^LacZ^* expression was maintained in the trigeminal, geniculate and epibranchial ectoderm, as well as the stomodeal roof and oral mxBA1, expression in the lateral SCE of mxBA1 and the FNP began to diminish at this time. β-galactosidase activity was also detected in the in-folding nasolacrimal groove. With progressive development from E11.5 to E13.5, transcription from the endogenous *Foxg1* locus continued to characterize the epithelia of the nasal invagination, the otic epithelia including the endolymphatic duct and semi-circular canals, the rostral optic cup and lens, and cells associated with the forming nasolacrimal duct and caudal vibrissae placodes. Notably, the developing primary (internal) choanae, primary palate and rostral secondary palate were all covered by positively-stained epithelia; however, at no time did we detect activity in the distal mdBA1.

We also visualized fully penetrant β-galactosidase activity in *Foxg1^+/−^* embryos in further locations previously un-reported or uncharacterized, including in cells representing the rostral midline cells of the roof plate of the mesencephalon and diencephalon beginning at E10.5 (Supplementary Figure 3). While these cells initially ran rostro-caudally along the midline, by E13.5 the line of *Foxg1* positive cells started to deviate from the midline, rostrally diverting laterally either to the left or right in a random fashion. Additionally, from E10.5 to E13.5 tracks of positive cells in the mesencephalon were detected running mediolaterally just parasagittal to the positive roof plate cells noted above. Moreover, at E13.5 positively-stained cells were seen lining the rim of the outgrowing auditory pinnae.

### Evidence of Perturbed Development and Pattern of External Cephalic Ectodermal Structures in Foxg1^−/−^ *Mutants*

Previous studies have implicated *Foxg1* in the process of neurogenesis of epibranchial, otic, optic and olfactory placodes (94, 103, 104, 107–109, 118, 131, 132); however, because of the extent and location of *Foxg1* expression during SCE ontogeny we posited that *Foxg1* also regulates the development of non-neural ectodermal derivatives of the SCE. We therefore examined the morphological consequences of the loss of *Foxg1* for the ontogeny of the external cranial anatomy. Beginning at E10.5 and continuing until birth the most obvious cranial defect of *Foxg1^−/−^* mutants was the hypoplasia of the telencephalon which left neonates with flat, rather than domed, crania (Figure 2; Supplementary Figures 3, 4). However, numerous other defects of external anatomy were patent by E15.5 in *Foxg1^−/−^* mutants, including dysmorphic, misplaced and misoriented auditory pinnae and auditory meatal antra, as well as disrupted topographic patterns and development of the mystachial vibrissae and upper lip (Figure 2; Supplementary Figure 3) and a smaller rhinarium with occluded nares. Also notable in *Foxg1^−/−^* mutants was the appearance of (typically) three rostrally placed ectopic vibrissal sinus tubercles in a pattern mimicking the arced pattern of the endogenous supra-orbital, post-orbital and post-oral tubercles (Figures 1A-D; Supplementary Figure 3A, B). The ectopic analogs of the supra-orbital and post-orbital tubercles are particularly notable for their topography as the former were often seen developing on the presumptive corneal surface of the mutant eyes and the latter on the maxillary rims of the inferior orbital furrows from which the lower eyelids normally develop. Moreover, unlike in wild type embryos, the orbital furrows in E15.5 *Foxg1^−/−^* mutants were typically not of uniform depth around the developing eyes, and a ventral expansion of the furrow was evident where the nasolacrimal grooves should have met the eyes. A general absence of the periocular mesenchyme (POM) that supports the development of the lower eyelids was also evident between the ectopic tubercles and the nasolacrimal depressions of *Foxg1^−/−^* mutants at E15.5; consequently, *Foxg1^−/−^* neonates often exhibited an ‘eyes open at birth’ (EOB) phenotype (Supplementary Figure 4B). These external defects of eye development coincide with the clear disruption of lens, conjunctival, corneal, and nictitating membrane development evident in *Foxg1^−/−^* mutants (Supplementary Figures 3, 4).

**Figure 2:**
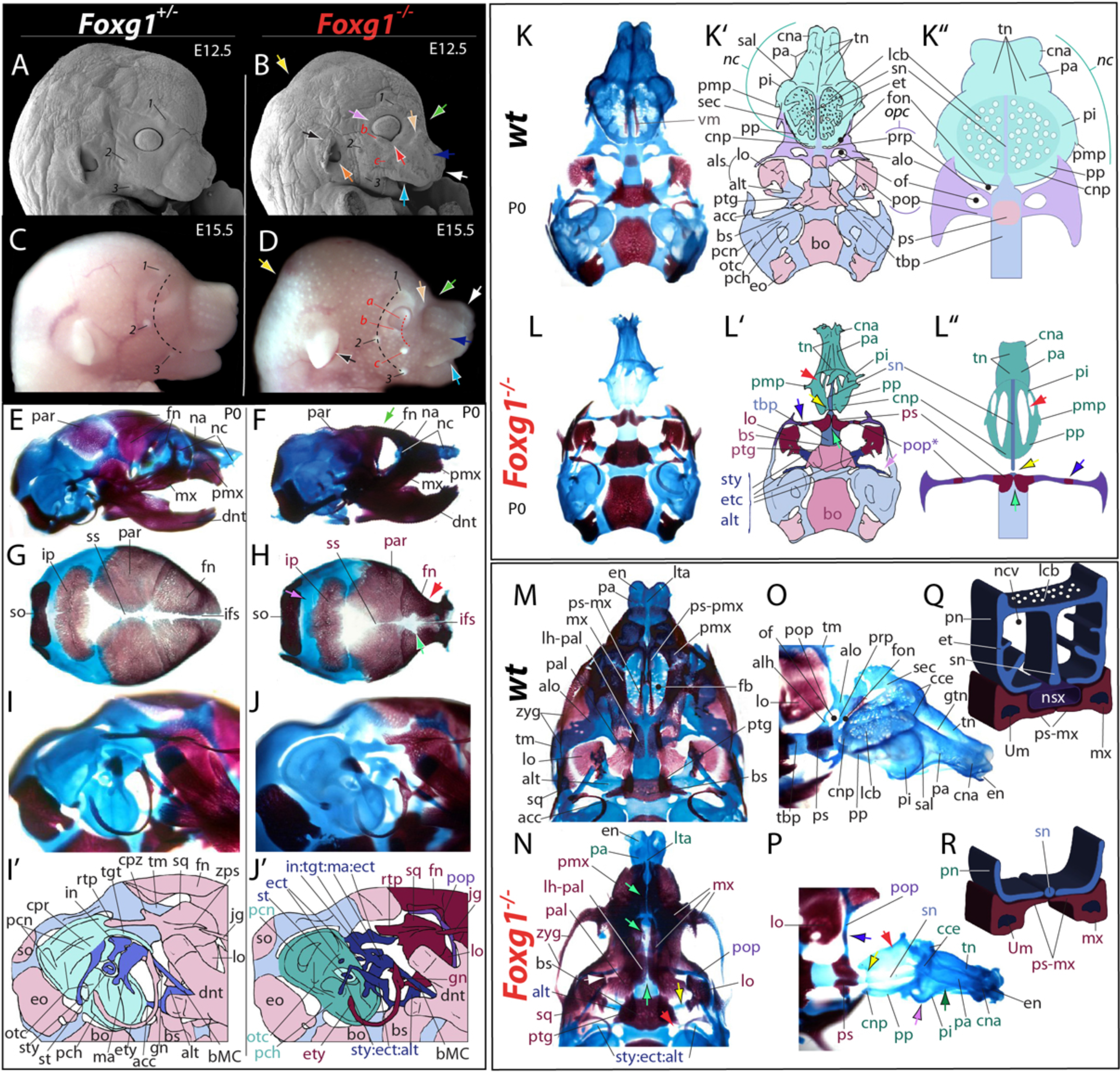
Significant non-neural craniofacial skeletal and ectodermal defects in *Foxg1*^−/−^ mutants. **(A-D)** Scanning electron micrographs <u>(**A,B**)</u> and gross external anatomy <u>(**C, D**)</u> of *Foxg1^+/−^* and *Foxg1^−/−^* littermates demonstrating dysmorphology mis-patterning of upper jaw and primary sensory organ system associated ectodermal structures. **<u>Arrows</u>**: ***green*** flattened crania highlighting the hypoplastic forebrain; **yellow** deficits of the posterior crania; ***black*** dysmorphic, misplaced, and mis-oriented auditory pinnae; ***orange*** external auditory metal antra; ***tan*** truncated post-vibrissal pad groove; ***dark blue*** hypoplastic, mis-patterned vibrissae and pads; ***light blue*** dysmorphic upper lip; ***white*** smaller rhinarium and occluded nares; ***red*** perdurance of the orbital (optical) terminus of the nasolacrimal groove; ***lavender*** under-developed, shallowed eyelid-corneal (conjunctival) furrow. ***1*** supra-orbital sinus follicle; ***2*** post-orbital sinus follicle; ***3*** post-oral sinus follicle; ***a, b, c*** ectopic sinus follicles. **<u>Dashed lines:</u> *black*** arc of primary sinus follicles; ***red*** arc of ectopic sinus follicles. <u>(**E-R**)</u> Cranial skeletal defects. Abbreviations in color indicate skeletal elements with altered morphology in the mutants while those connected by a colon indicate aberrant synchondrosis. <u>(**E-F**)</u> Differentially stained *Foxg1^+/−^* and *Foxg1^−/−^* neonates viewed in *norma lateralis*. ***green arrow*** highlights the flattened crania and loss of optic capsular and associated skeletal structures. <u>(**G, H**)</u> Calvarial alterations, viewed in *norma verticalis* externa. **<u>Arrows</u>**: ***red*** medio-lateral compression of fronto-orbital skeleton; ***light green*** characteristic pyramidal expansion of the posterior interfrontal (metopic) suture; ***lavender*** expanded lambdoidal suture. <u>(**I-**</u> **<u>J’</u>**) Altered structure and pattering of the otic capsule (i.e., inner ear) and branchial arch-derived primary jaw joint elements (i.e., mammalian middle ear) and secondary, junctional upper jaw skeleton viewed in norma lateralis. ***mauve*** normal ossified structures; ***steel blue***, normal non-otic chondroranium; ***blue*** branchial first and second arch-derived chondrocanium; ***green*** otic capsule. <u>(**K-L”**)</u> Loss of structure and integration of the nasal and optic capsules and the trabecular basal plate. Most of the anterior dermatocranial bones and the calotte have been removed to increase clarity. Viewed in *norma verticalis interna*. Colors as above. **<u>Arrows</u>**: ***red*** paris nasi; ***yellow and black*** discontinuity of nasal septum and the more caudal trabecular basal plate; ***light green*** lack of full media-lateral integration of the (here precocious) ossification centers of presphenoid; ***blue*** precocious ossification center in dysmorphic post-optic pillar of the optic capsule; ***lavender and white***, ectopic structure comprised of realigned alicochlear commissure (from the Ala temporalis) and styloid components. <u>(**M-R**)</u> Aberrant upper jaw development, including palatal dysmophology and loss (topographical collapse) of the nasal cavity and pharynx. Most of the anterior dermatocranial bones in **O** and **P** have been removed to increase clarity. **<u>Arrows</u>**: ***red*** paries nasi posterior; ***yellow and black*** point of nasal septal and trabecular basal plate discontinuity; ***green*** -paries nasi intermediale; ***lavender*** paries nasi posterior; ***blue*** post-optic pillar of the hypomorphic optic capsule; ***green*** palatal midline; ***white*** change in the relative position of the lamina pterygopalatina of the palatine and the lamina obturans and post-optic pillar. **<u>Abbreviations</u>**: ***acc*** alicochlear commissue; ***alh***, ala hypochiasmatica; ***alo***, ala orbitalis; ***als***, alisphenoid; ***alt***, ala temporalis; ***bMC***, body of Meckel’s cartilage; ***bo***, basioccipital; ***bs***, basisphenoid; ***cce***, crista cribroethmoidalis; ***cna***, cupula nasi anterior; ***cnp***, cupula nasi posterior; ***cpr***, crista parotica; ***cpz***, caudal process, zygomatic; ***dnt***, dentary; ***ect***, ectopic structure; ***en***, external nares; ***eo***, exoccipital; ***et***, ethmoturbinal; ***ety***, ectotympanic; ***fb***, fenestra basalis; ***fn***, frontal; ***fon***, fissura orbitonasalis; ***gn***, gonilae; ***gtn***, groove, tectum nasi; ***ifs***, interfrontal suture; ***in***, incus; ***ip***, interparietal; **jg**, jugal; ***lcb***, lamina cribrosa; ***lh-pal***, lamina horizontalis, palatine; ***lo***, lamia obturans; ***lta***, lamina transversalis anterior; ***ma***, malleus; ***mx***, maxillae; ***na***, nasal; ***nc***, nasal capsule; ***ncv***, nasal cavity; ***nsx***, nasopharynx; ***of***, optic foramen; ***opc***, optic capsule; ***otc***, otic capsule; ***pa***, paries nasi anterior; ***pal***, palatine; ***par***, parietal; ***pch***, pars chochlearis, otc; ***pcn***, pars canalicularis, *otc*; ***pi***, paries nasi intermediale; ***pmp***, processus maxillae posterior; ***pmx***, premaxillae; ***pn***, paries nasi; ***P0***, post natal day 0; ***pop***, post-optic pillar; ***pp***, paries nasi posterior; ***prp***, pre-optic pillar, opc; ***ps***, presphenoid; ***ps-mx***, palatal shelf, mx; ***ps-pmx***, palatal shelf, pmx; ***ptg***, pterygoid; ***rtp***, retrotympanic process, sq; ***sal***, sulcus anteriorlateralis; ***sec***, sphenethmoid commissure; ***sn***, septum nasi; ***so***, supraoccipital; ***sq***, squamosal; ***ss***, sagittal suture; ***st***, stapes; ***sty***, styloid process; ***tbp****, t*rabecular basal plate; ***tgt***, tegmen tympani; ***tm***, taenia marginalis; ***tn***, tectum nasi; ***Um***, upper molar; ***vm***, vomer; ***zps***, zygomatic process, squamosal; ***zyg*** zygomatic process.

### *Foxg1^−/−^* Mutants Have Significant Neurocranial and Viscerocranial Defects, Including of the Primary Sensory Capsules (Otic, Optic and Olfactory), the Upper Jaws, Middle Ears and Calvaria

Previous developmental studies of *Foxg1* have primarily addressed its potential roles in the control of neurodevelopment. Because of its pattern of expression on both sides of the ANR, we hypothesized that *Foxg1* also plays a significant role in cranial skeletogenesis and integration, in particular in the development of the upper jaws. To address this hypothesis directly we specifically analyzed the development of the skulls of *Foxg1^−/−^* mutants and their littermates from E15.5 to P0 both through histological sectioning and by differential skeletal staining using alizarin red for bone and alcian blue for cartilage (133, 134). This analysis demonstrated that *Foxg1^−/−^* mutants exhibit severe perturbations of cranial skeletal development in diverse craniofacial regions (Figure 2; Supplementary Figure 4).

#### Neurocranial Defects of Foxg1^−/−^ mutants

The neurocranium is that portion of the skull which houses the brain and primary sensory capsules and is composed of both chondrocranial (cartilage-derived) and dermatocranial (membrane bone-derived) components (5, 7, 11, 135, 136). Defects were evident in both these components of the *Foxg1^−/−^* neurocranium (Figure 2 and Supplementary Figure 5). The absence of an expanded telencephalon and the presence of an abnormal optic stock (see below) left the dermatocranial frontal bones clearly highly dysmorphic, reduced medio-laterally, and lacking orbital laminae. *Foxg1^−/−^* neonatal calvaria also exhibited open anterior fontanelles between the frontal bones, displaying distinctly pyramidal shaped extended patency of the posterior interfrontal (metopic) sutures (Figure 2H). The parietals and interparietals, moreover, were often smaller which led to wider lambdoidal sutures.

Within the neurocranium, the otic capsule (OTC) envelopes the vestibular and cochlear apparatuses in the pars canalicularis and pars cochlearis, respectively; both of these were dysmorphic in *Foxg1^−/−^* neonates (Figures 2I-J’). Reflective of defects of the semi-circular canals, within the pars canalicularis both the prominentia semicircularis anterior and posterior were abbreviated. Moreover, the oval and round windows appeared anomalous in size and orientation in the capsules. Within the pars cochlearis, the cochlea failed to make the usual one and a half turns. Additionally, two extra-capsular structures that are normally synchondrotic with the otic capsule, the tegmen tympani (TT; a mammalian neomorphic structure) and the styloid process, were both found to be dysmorphic and disconnected from the OTC and were otherwise found anomalously associated with the altered middle ear skeleton (see below). A third extra-capsular structure that normally connects the rostromedial margin of the pars cochlearis to the ala temporalis of the alisphenoid, the alicochlear commissure, was also dysmorphic and misrouted from joining the OTC to joining with the remnants of the styloid process (see below).

While relatively small, the murine OPC is comprised of pre-optic and post-optic pillars that, together with the laterally placed ala orbitalis and medial presphenoid, frame the optic foramen. Normally, the anteriolateral margin of the ala orbitalis rostrally extends a sphenethmoidal (orbitonasal) commissure that meets the dorsolateral NC, thus forming the lateral boundary of the orbitonasal fissure that separates the nasal and optic capsules. Likewise, the ala orbitalis normally extends a caudal posterior commissure to meet the taenia marginalis of the neurocranial sidewall. The OPC of *Foxg1^−/−^* neonates, however, lacked a pre-optic pillar and most of the ala orbitalis (Figures 2K-P), and thus also any connections to the NC. Instead, a single, aberrantly straight rod-like post-optic pillar with a precocious ossification center, a sharp thin posterior commissure directed towards a diminished taenia marginalis, and a precociously ossified ala hypochiasmatica was found extending laterad from the caudal boundaries of the presphenoid. The preshenoid was found to be dysmorphic and evincing extended, precocious lateral ossification but a distinct lack of ossified unity at (and across) the midline.

Development and morphogenesis of the NC was also significantly perturbed in *Foxg1^−/−^* mutants (Figures 2F, K-R; Supplementary Figure 4). Because the normally developing murine nasal cavity is structurally complex and developmentally intimately tied to the λ-junction and the forming upper jaws a broader description of its normal skeletal development is expedient. The perinatal skeleton of the murine nasal cavity (cavum nasi) is composed of two bilateral NC separated medially by an extension, the nasal septum (septum nasi), of the midline trabecular basal plate (TBP) that underlies the brain rostral to the hypophysis. The TBP is formed from paired midline extensions (trabeculae cranii) running rostrad from the center of the basi-sphenoid (at the hypophysis), through the presphenoid and extending rostrad to form the septum nasi between the NC. These paired structures typically condense and chondrify in such a manner that a single midline cartilaginous structure and presphenoidal ossification center is normally seen in skeletal preparations. To either side of the rostroventral part of the nasal septum lie paraseptal cartilages that house the vomeronasal organs (VNO). The NC themselves are paired cartilaginous sacs formed from frontonasal CNC condensing around the invaginating olfactory pits, each generating the roof, sidewall and floor of the cavum nasi.

Anteriorly, the dorsal NC normally arcs medially to fuse to the nasal septum thereby forming the midline grooved anterior roof, the tectum nasi. The anterior extremity of the tectum extends beyond the overlying dermatocranial nasal bones and ends with the cupula nasi anterior and the outer nasal cartilages that frame the external nares and support the rhinarium. The caudal border of the tectum, the crista cribroethmoidalis, forms a sharp crest that meets the obliquely oriented lamina cribrosa that forms the caudal NC roof, underlies the olfactory bulbs, and is perforated by the olfactory nerves and ethmoidal vessels.

Each NC sidewall, or paries nasi, is grossly composed of three subdivisions: the pars anterior, pars intermediale, and pars posterior. It is the paries nasi that essentially establishes the functional organization of the interior nasal spaces. Rostrally, the elongate, narrow pars anterior helps to establish the anterior vestibule (the space just internal to the external nares) associated with the cupula nasi anterior and the initial management of nasal respiration. The murine pars anterior does not form a complete sidewall and it is ventrally fenestrated for the passage of vessels and ducts. The middle subdivision, the pars intermediale, contains a laterally expanded prominentia Iateralis that houses an internal meatus, the recessus lateralis. It is separated from the pars anterior by the largely dorsoventrally oriented sulcus anteriolateralis. The external surface of the caudolateral prominentia extends a small lateral spur, the processus maxillae posterior, thought to be of extra-capsular (mxBA1) origin. The pars posterior forms the caudal NC sidewall, the interior of which is the recessus ethmoturbinalis. The pars intermediale and pars posterior are both roofed by the lamina cribrosa.

The NC floor, or solum nasi, is necessarily incomplete as the internal and external choanae must join to form a continuous respiratory passage: this is achieved in part by an anteroposterior vacuity of the solum, the fenestra basalis, that lies between the medial nasal septum and paraseptal cartilages and the lateral paries nasi. The anterior solum nasi is formed by the paired lamina transversalis anterior that meet the nasal septum at the ventral midline. At the caudal end, the floor takes contribution from the paired lamina transversalis posterior, extensions from the planum antorbitale of the pars posterior. The caudomedial end forms the lamina orbitonasalis that meets the TBP (here, forming the interorbitaI septum) medially, the lamina cribrosa superiorly and the lamina transversalis posterior inferiorly thus forming the cupuIa nasi posterior.

The internal landscape of each NC is ontogenetically dominated by the progressive development and ramification of turbinals (turbinates, or nasal conchae), capsular projections into the nasal fossa from the tectum, paries nasi and lamina transversalis that greatly increase the surface area covered by the various nasal epithelia. The anterior-most murine turbinal, the atrioturbinal, develops from the lamina transversalis anterior in each nasal fossa; caudally, it is followed by a nasoturbinal descending from the tectum and a maxilloturbinal that projects from the pars intermediale. Both of these latter turbinals normally extend rostrally through much of the pars anterior. A cartilaginous turbinal crest, the crista semicircularis, projects inward from the prominentia lateralis thereby segregating the anterior and middle nasal chambers. Within the prominentia lateralis, a horizontal lamina functionally offsets an upper recessus frontoturbinalis and a lower recessus maxillaris. From the pars posterior and lamina cribrosa arise multiple frontoturbinals (superiorly) and several ventral ethmoturbinals.

Most of the above described complex NC structures were found to be deficient or lacking in the absence of *Foxg1*. Overall, the *Foxg1^−/−^* perinatal snout was compressed mediolaterally, dorsoventrally and rostrocaudally, and this is reflected in changes to the NC (Figure 2 and Supplementary Figure 5). A nascent rhinarium and external capsular narial openings were patent in mutant perinates (see Fig. 9C3); however, the cupula nasi anterior was smaller and the outer nasal cartilages were anomalous in size and orientation. The initial intranarial epithelium was thicker in *Foxg1^−/−^* mutants, however, and led to a smaller vestibular entrance. The tectum nasi with shortened nasoturbinals, the nasal septum, and the lamina transversalis anterior with atrioturbinals were each present in mutant skulls though they were all generally less refined and thicker. Pars anterior with diminished maxilloturbinals also characterized the NC in *Foxg1^−/−^* perinates. However, caudal to the laminae transversalis anterior and atrioturbinals, the floor of the mutant NC was disorganized and asymmetric; and, while present, paraseptal cartilages were dysmorphic, misaligned and lacked VNO.

Although affected by the loss of *Foxg1*, the rostral NC was of more or less patent; the middle and caudal NC, however, were devastated by either the truncation or the complete loss of structure. The pars intermediale lacked both an expansive prominentia lateralis and its associated crista semicircularis; thus, there were no recessus laterales nor sulci anteriolaterales. Presumptive processus maxillae posterior did, however surprisingly, project from the diminished pars intermediale of mutant skulls. In this mid region of the NC, the septum of *Foxg1^−/−^* mutants typically became discontinuous and asymmetric dorsoventrally. The tectum nasi of mutant perinates ended at an abrupt crista cribroethmoidalis. Significantly, however, caudal to the crista there were no laminae cribrosae, and the caudal NC was represented only by ventrolateral, unadorned cartilaginous plates. Thus, there were no cupulae posterior, frontoturbinals or ethmoturbinals nor any of their support structures. The caudal nasal septum, moreover, presented no dorsal extension and was represented only by a cylindrical interorbitaI septum which was itself typically abnormally disconnected from the remainder of the TBP just rostral to the presphenoid. Notably, the loss of the posterior NC meant that the nasal cavities of *Foxg1^−/−^* mutants did not empty ventrally into a nasopharyngeal duct, and they would have but for a thin caudal epithelial barrier emptied directly into the cavum cranii itself.

#### Upper Jaw Dermatocranial Defects of Foxg1^−/−^ mutants

A number of dermatocranial elements, including jaw elements, normally develop in association with the NC, OPC, and TBP (5, 7, 11, 135, 136). For instance, paired nasal bones overlap the tectum nasi while frontal bones dorsally cover the olfactory bulbs, overlap the crista cribroethmoidalis, and are associated with the OPC. Laterally and ventrally, the maxillae cover the pars intermediale and pars posterior, while lacrimal bones are associated with the maxillae and orbital face of the NC. The lateral and ventral pars anterior are covered by the premaxillae, the palatine (palatal) processes of which underlie the paraseptal cartilages. Caudal to this, the paired vomers frame the ventral nasal septum while the maxillary and palatine bones overlap the caudal NC and TBP (including the presphenoid), respectively.

The defects associated with the tectum nasi and pars anterior of the NC were accompanied by alterations of the nasal and premaxillary bones (Figure 2). The nasal bones of *Foxg1^−/−^* mutants appeared mediolaterally compressed, often asymmetric, and rostrally under ossified. Premaxillae were associated with the lateral and ventral aspects of the anterior NC in *Foxg1^−/−^* mutants; however, the inter-premaxillary foramen was pushed rostrad and extended inter-premaxillary sutures were evinced. The bodies of the premaxillae contained incisors, though they were slightly anomalous in shape. Oddly, the rostral tips of each incisor was typically associated with an extremely small, yet very clear, independent mineralized nodule (Supplementary Figure 5). Moreover, two of 15 premaxillae specifically examined housed ectopic incisors. The maxillary processes were shortened and there were no discrete palatine processes *per se* projecting caudally at the midline and overlying the dysmorphic paraseptal cartilages.

The maxillae are viscerocranial bones (i.e., BA-derived; see below) and are the largest elements of the perinatal upper jaw. They mature to take a significant part in forming the mature nasal, orbital and oral cavities and are thus discussed here in this context. In *Foxg1^−/−^* neonates the maxillae were dysmorphic, in part because there were no palatine (palatal) processes as such projecting rostrad at the midline and framing the fenestra basalis. Rather, in the mutant maxillae, un-elevated (see below) and asymmetric osseous plates of the maxillary body were seen projecting both toward the midline (though they were never close to abutting and thus left an unossified cleft) and rostrad toward the premaxillae (though never reaching them). Moreover, between these plates, small isolated ectopic ossifications were encountered. Loss of the prominentia lateralis of the NC was associated with the loss of the frontal process of the maxillae and, while projecting caudally, the maxillary zygomatic processes showed less of an arc and were thinner. The bodies of the maxillary bones of *Foxg1^−/−^* mutants housed developing molars. The lacrimal bones, usually associated with the maxillary frontal processes and the prominentia lateralis, were smaller and misshaped. Paired medial vomer bones, which usually underlie the maxillary palatine processes and are associated with the nasal septum, were present but did not elevate and were often asymmetric (following asymmetries associated with the septum nasi).

Overall, the junction of the primary and secondary palates (such as they were) was dysmorphic, and the fenestra basalis (the incisive foramen precursor and a very prominent attribute of palatal development in mice) is mostly obscured in *Foxg1^−/−^* mutant perinates by the aberrant medial ossification patterns. Moreover, patency of the maxillary-premaxillary sutures associated with the elemental bodies of these bones was often disrupted by aberrant synostosis; synostosis was also found across the palatine-maxillary sutures. The palatal shelves of the palatine bones did project toward the midline. Notably, however, none of the ‘palatal’ shelves of the premaxillae, maxillae or palatine (here, the lamina horizontalis) bones were properly elevated from the ventral neurocranial base of either the NC or the TBP, and thus mutants lacked any suprapalatal respiratory passage.

Importantly, the upper jaw arcade also includes the squamosal, which is a key integrating dermatocranial element contributing to the posterior orbit, the cranial (temporal) sidewall, and the functional jaw articulation. The normal squamosal is characterized by a small central body that houses the mandibular (glenoid) cavity and from which a number of lamina and processes extend. A rostrodorsal squamosal lamina, overlying the parietal plate of the chondrocranial sidewall, extends rostrad to the frontal bone and the lamina obturans of the alisphenoid, while a zygomatic process arcs out to the jugal in the zygomatic (orbital) arch, and a ventrocaudal sphenotic lamina runs toward the cupula cochlea and the sphenoid. Moreover, there are additionally two posterior processes on the murine squamosal: a retrotympanic process that runs caudad to overlie the TT and middle ear elements and a caudal process that extends dorsocaudally between the retrotympanic and the squamosal lamina. Most of these components of the squamosal bones of the *Foxg1^−/−^* skulls exhibited deficiencies. While an articulation with the condylar process of the dentary was present, each lamina and process of the squamosal bones of *Foxg1^−/−^* mutants was either truncated (e.g., squamosal, sphenotic, caudal and zygomatic) or absent (e.g., retrotympanic).

#### Viscerocranial Elements, including those in the Middle Ear, are Mis-patterned and Deficient in Foxg1^−/−^ mutants

Viscerocranial elements originate from the BA, consist of both splanchnocranial (chondrocranial) and dermatocranial elements, and include the jaws and middle ear ossicles (5, 7, 11, 135, 136). For instance, from each mxBA1 are derived the incus (splanchocranial), the maxillae, palatine, pterygoid, squamosal, jugal (dermatocranial) and the alisphenoid (both). From mdBA1 arises the core of the lower jaw viscerocranium — the chondrocranial Meckel’s cartilage (MC; the proximal end of which yields the malleus) — and the dermatocranial dentary, ectotympanic, and gonial bones that are associated with the MC, the malleus and the auditory bulla. The second, or hyoid, arch (BA2) viscerocranium lacks a dermatocranial contribution in mice; its splanchnocranial contribution, however, consists of the various derivative components of Reichert’s cartilage. These include the stapes of the middle ear, the styloid process (itself formed of a proximal tympanohyal attached synchondroticly to the otic capsule at the crista parotica and a curved elongate stylohyal covering the round window), as well as the lesser horns and part of the body of the hyoid bone.

In *Foxg1^−/−^* mutants, the development of the largest viscerocranial elements of the lower jaws, the BA1-derived dentary and body plus rostral process of MC, were largely spared; the remainder of the BA1 and proximal BA2 splanchnocranial elements were, however, not (Figure 2). The malleus and incus of *Foxg1^−/−^* mutants failed to form a synovial joint; instead, they were synchondrotic between the neck of the malleus and the body of the incus. The crus brevis of the incus was itself abnormally synchondrotic with an enlarged dorsal ectopic cartilage that, based on its position overlying the incus and rostrocaudal to the pars canalicularis, we have taken as the remnant of TT which has been detached from the otic capsule (labelled as ‘in:tgt:ma:ect’ in Figure 2J’). The mxBA1-derived alisphenoid, including the dermatocranial lamina obturans and the ala temporalis that forms the cartilaginous base of the alisphenoid and underlies the trigeminal ganglion, was found to be significantly altered in *Foxg1^−/−^* mutants. For instance, the horizontal lamina was thinner and both the prominent pterygoid process and the anterolateral process were lacking. Moreover, the mutant lamina typically was not met by an alicochlear commissure coming from the pars cochlearis; instead, a mediolaterally oriented ectopic cartilaginous strut was present in the region (labelled as ‘sty:ect:alt’ in Figures 2J’, 2N). In the mutant skulls, this strut was seen to first run laterally toward the malleus before bending caudally and becoming synchondrotic with the stylohyal of the styloid process. Rather than a tympanohyal attaching to the crista parotica, *Foxg1^−/−^* mutants topographically presented a small cartilage devoid any skeletal connection to the otic capsule. Furthermore, the stapes was only loosely associated with the window and lacked both a stapedial foramen and any functional connection with the crus longus of the incus.

The dermatocranial elements of the middle ear and nasopharynx were likewise found to be affected by the loss of *Foxg1*. In addition to above mentioned maxillary, palatine and squamosal bones, *Foxg1^−/−^* mutant mxBA1-associated pterygoid bones were dysmorphic. The pterygoids, normally developing in close association with the lateral margin of the basisphenoid and occupying the lateral margins of the nasopharynx, were malformed, dorsoventrally flattened and synostotic with the basisphenoid. Two proximal (i.e., hinge regional) mdBA1 associated dermal bones, the gonial and ectotympanic, were also slightly aberrant in *Foxg1^−/−^* skulls. The gonial, which normally contributes to the middle ear by its direct osseous investment of the cartilaginous malleus, was shortened, and the breadth of the curvature of the mutant ectotympanic bone, which normally rims the tympanic and contributes to the auditory bulla, was found to be condensed in the mutant skulls, consistent with changes in the meatus.

Phenotypic analysis of *Foxg1^−/−^* crania from E15.5 to P0 demonstrates that *Foxg1* regulates the development of the skeleton associated with the λ-junction, including the upper jaws, NC, and OPC, and their associated support skeletons. Furthermore, the loss of *Foxg1* has additional significant consequences for the pattern and morphogenesis of much of the cranial skeleton, including each primary sensory capsule, the middle ear, and their associated structures. The skeletal defects of the perinatal mutant skull were also accompanied by disruptions of many of the associated soft tissues (Supplementary Figures 3, 4). For instance, in addition to the absence of the VNO itself, development of the ductal systems of both the orbital (e.g., the harderian glands) and intra-NC glands were vitiated. Moreover, the presence, topography, and transitions of the extra-capsular, respiratory, transitional and olfactory epithelia were anomalous in *Foxg1^−/−^* mutants. Though present, both the hypophysis and trigeminal ganglion additionally presented anomalies of development, as notably did the eyes, the eyelids and the aforementioned auditory pinnae.

### Alterations of both *Foxg1* transcriptional dynamics and the structural organization of the mature λ-junction in *Foxg1*^−/−^ embryos

Both the perturbation of the development and pattern of external SCE derivatives and the significant re-patterning and dysmorphogenesis of the upper jaws, OPC, and NC evinced in *Foxg1^−/−^* mutants suggested that the developmental elaboration of the λ-junction was affected by the loss of *Foxg1*. To initiate an investigation of possible disruptions in the ontogeny of the λ-junction, we examined the λ-junction of *Foxg1^−/−^* embryos and their littermates just after ‘maturation’ (as seen at E11). This developmental stage occurs just after the establishment of significant relative topographical relationships of substantial and essential patterning information that is manifest around E10.5 (7, 9).

We first investigated the patterns of transcriptional activity of the endogenous *Foxg1* locus in mutant embryos and compared them to those of their *Foxg1^−/−^* littermates. At E11, the pattern of activity detected in *Foxg1^+/−^* embryos followed that described above; *Foxg1^−/−^* mutant embryos, however, exhibited a distinct pattern of activity within the SCE associated with the λ-junction and the development of the upper jaw (Figure 3A-F). Unlike their E11 heterozygous littermates, *Foxg1^−/−^* embryos showed more intense, extensive staining in the SCE of the lateral and oral aspects of mxBA1, the trigeminal ganglion, and the optic primordia, and throughout the SCE of the stomodeal roof. A similar aberrant increase was detected in the SCE associated with both the lFNP and mFNP of E11 *Foxg1^−/−^* embryos. Additionally, ectopic X-gal staining was detected at this time point in the epithelium of mutant embryos between the otic vesicle and the epibranchial placodes.

**Figure 3:**
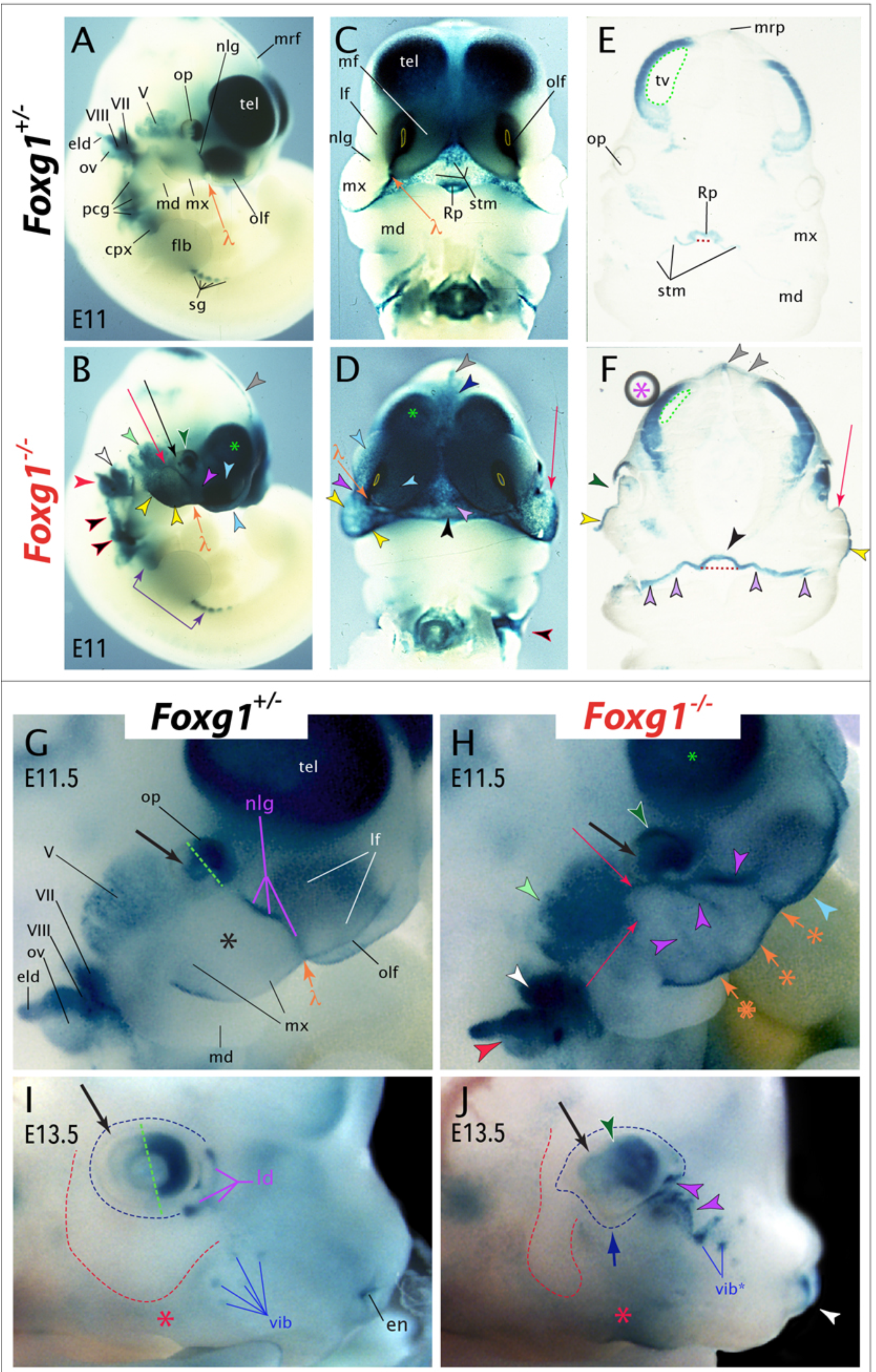
Loss of *Foxg1* alters the dynamics of its own transcription, the development of the SCE and λ−junction, and morphogenesis of the upper jaw craniofacial primordia. **<u>(A-F)</u>** Transcription of the *Foxg1* locus in *Foxg1^+/−^* (*i.e*, *Foxg1^LacZ/+^*) and *Foxg1^−/−^* littermates as detected in E11 embryos by X-gal staining for β-galactosidase enzymatic activity. Yellow dashed lines, outlining the openings of the olfactory pits, highlight the change in the orientation of the pits and frontonasal processes. Green asterisk indicates the under-developed telenchehalon. <u>(**E, F**)</u> Coronal sections of the same embryos in **A** and **B**. Dashed red lines indicate the width of invaginating Rathke’s pouch while green dashed lines outline the lining of the telencephalon vesicles. The purple asterick indicates a preparation artifact. <u>Arrows</u>: *orange* lambdoidal junction; *red* ectopic mxBA1 bulge; *black* increased X-gal staining in the temporal optic primordia; *purple* relative staining of the *cpx* and *sg*. **<u>Arrowheads</u>**: *light blue* frontonasal processes; *green and white* demarcation line of nasal (rostral) optic cup staining; *light green and black* V; *white and black* VII; *red and black* otic vesicle, including endolymphatic duct, and VIII; *yellow and black* margins of maxillary process; *black and red* aberrant staining within ganglia for IX, X, XI; *purple and black* presumptive nasolacrimal groove; *lavender and black* altered expression within the stomodeal ectoderm; *black* Rathke’s pouch; *grey and black* the forebrain midline; *dark blue and black* expression lateral to the roofplate. <u>(**G-H**)</u> Transcription at E11.5 highlighting dysmorphogenesis associated with the λ−junction, including the endogenous nasolacrimal groove and two ectopic grooves within mxBA1 . The black asterick in the heterozygote overlies a mxBA1 with little ectodermal X-gal staining. Orange arrowheads with astericks indicate discontinuities at the upper lip margin associated with the ectopic grooves (*purple and black* arrowheads). Other arrows and arrowheads as above. <u>(**I, J**)</u> Transcription at E13.5 makes clear the disruption of the lacrimal and nasolacrimal ductal system, vibrissae, and eyelid-corneal furrow and its association with *Foxg1* (β-galactosidase) expressing cells. <u>Dashed lines</u>: *green* delineates the highly stained nasal (rostral) optic cup from the lightly stained temporal (caudal) primordia; *red* demarcate the boundaries of stained (*red asterick*) and unstained post- and sub-orbital, maxillary epithelia; *blue* outline the boundaries of the eyelid-corneal furrow. <u>Arrows</u>*: blue* diverticulum of the lower eyelid furrow; *black* enhanced temporo-optic ectodermal signal. <u>Arrowheads</u>: p*urple and black* β-galactosidase activity associated with the disrupted nasolacrimal epithelium; g*reen and white* highlights the disorganization of the optic cup, lens and corneum characteristic of the *Foxg1^-/^*mutants; *white* extended string associated with the rhinarium and external nares, **<u>Abbreviations</u>**: *cpx*, cervical plexus; *en*, external nares; *flb*, forelimb bud; *lf*, lateral frontonasal process; *md*, mandibular BA1; *mf*, medial frontonasal process; *mrp*, midline, roof plate; *mx*, maxillary BA1; *nld, lacrimal and* nasolacrimal ducts; *nlg*, nasolacrimal groove; *olf*, olfactory pit; *op*, optic primordial; *ov*, otic vesicle; *pcg*, posterior cranial ganglia; *Rp*, Rathke’s pouch; *sg*, presumptive sympathetic ganglia; *stm*, stomodeum; *tel*, telencephalon; *tv*, telencephalic vesicle; *vib*, vibrissae primordia; *V*, trigeminal placodal gangli; *VII*, facial placodal gangli; λ, center of the lambdoidal junction.

This striking difference in the *Foxg1* transcriptional profile between heterozygous and mutant embryos led us to examine the morphology of E11 embryos in greater detail. This assessment highlighted five further areas of morphologic distinction visible by light microscopy between the E11 *Foxg1^−/−^* embryos and their heterozygous littermates (Figure 3). First, in *Foxg1^−/−^* embryos the FNP and mxBA1 were positionally reoriented and rotated ventrolaterally, an alteration accompanied by an increase in the breadth of Rathke’s pouch and the entire stomodeum. Second, this increased stomodeal breadth was associated with a decrease in epithelial thickness of Rathke’s pouch but a general increase in the thickness of the epithelial layer of the remainder of the SCE of the stomodeal roof. Third, the olfactory pits of mutant embryos were smaller and less elongate. Fourth, the architecture of the developing lens and optic cup was already clearly anomalous and disorganized with the loss of *Foxg1*. And fifth, a distinctly anomalous, dorsolateral bulge of tissue protruded from the mxBA1 ventro-caudal to the eye and dorsal of the mxBA1-mdBA1 boundary of mutant embryos.

These trends were still patent at E11.5, though this time point coincided with additional morphological changes of mxBA1 and the lFNP. For instance, rather than a single nasolacrimal groove running from the developing orbit to the center of the λ-junction, E11.5 *Foxg1^−/−^* embryos displayed two additional (ectopic) shallow grooves within mxBA1 and a dorsal reorientation of the groove between mxBA1 and the lFNP. Moreover, the invaginating olfactory pits of *Foxg1^−/−^* mutants continued to be both less deep and less elongate.

By E13.5, *Foxg1* heterozygous embryos exhibited, as expected, β-galactosidase activity along the rim of the invaginated lens vesicle and in the rostral optic cup as well as in the forming nasolacrimal ductal system exiting the developing orbit (Figure 3). High levels of activity were further detected in the epithelia associated with the external nares and in discrete vibrissae primordia. Low levels of activity were also detected along the upper lip and vibrissal pad; however, the maxillary region below the eye was distinctly absent of even low-level activity. By comparison, E13.5 *Foxg1^−/−^* embryos displayed β-galactosidase activity throughout most of the malformed eye, as well in a large area of surface epithelium associated with the abortive nasolacrimal ductal system leaving the orbit. Additionally, a large swath of lightly positive staining was detected in the maxillary epithelium from the eye and nasolacrimal region to the margin of the upper lip. Thus, the loss of *Foxg1* clearly alters both the dynamics of its own transcription and the ontogenetic elaboration of the upper jaw-associated facial primordia morphogenesis.

### Structural reorganization at the λ-junction in *Foxg1^−/−^* mutants includes the loss of the coordinated directionality and orientation of the oral epithelium, the loss of internal choanae, and impaired elevation of the palatal shelves

The observed gross changes in regional morphology, including in the apparent increased thickness of the oral epithelium, coupled with regional changes in the transcriptional activity of the endogenous *Foxg1* locus, led us to further examine the embryonic cranial epithelium. Because scanning electron microscopy (SEM) is known to provide crisp, detailed images of surface characteristics of embryos (137, 138), we utilized SEM to examine the cranial epithelia of *Foxg1^−/−^* embryos and their heterozygous littermates.

Removal of the caudal, post-maxillary embryo permits visualization of the oral aspect of the λ-junction as well as the stomodeal roof and internasal groove. While low magnification SEM micrographs of E11.5 *Foxg1^−/−^* embryos clearly showed the wild type demarcation of landmarks associated with the λ-junction as well as the stomodeal roof and internasal groove (detailed in Figure 1), micrographs of mutant embryos both confirm defects already noted by light microscopy and reveal less obvious alterations in the orientation and/or topography of these same landmarks (Figures 4A, B). For example, the internasal line and nasolacrimal groove were seen to be shallower and shorter in mutants embryos, while the ventral intranasal line, stomodeal-palatal line, and oronasal groove were each found to be shallower and longer. Additionally, SEM micrographs of mutant embryos confirmed that mutant embryos were characterized by the re-orientation of the FNP as well as a medio-lateral expansion of the stomodaeum and Rathke’s pouch.

**Figure 4:**
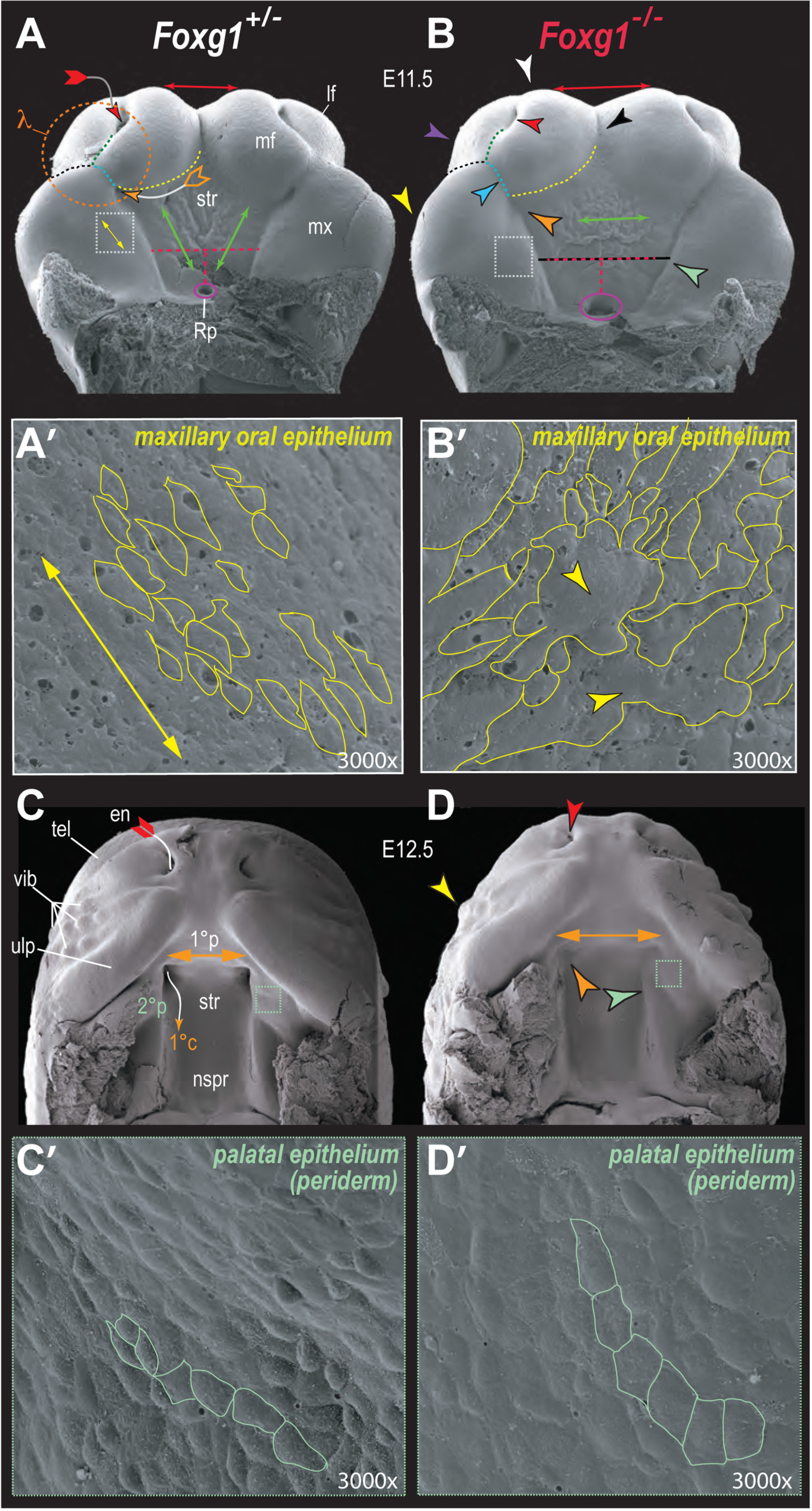
Choanal atresia, truncation of palatal elevation, and mis-coordination of cellular directionality, size, and orientation of the upper jaw associated epithelia. <u>(**A, B**)</u> Scanning electron micrographs of the stomodeum and λ−junction of E11.5 *Foxg1^+/−^* and *Foxg1^+/−^* littermates. The lower jaw primordia have been removed for clarity. *Red and black tipped arrow* indicates the normal invaginating olfactory pit, while the *orange and black arrow* highlights the normally invaginating primary (internal) choanae. *Purple circles* outline the invaginating Rathke’s pouches. **<u>Dashed line</u>s**: *red* relative width of the stomodeal roof; *green* ventral intranasal line; *black* nasolacrimal groove; *light blue* oronasal groove; *white* fields of superficial maxillary oral epithelium magnified in **A** and **B; yellow** nasostomodeal line; *3* outer boundaries of the λ-junction . Yellow lines outline cell boundaries in the developing eye. **<u>Arrowheads</u>**: *red and black* shallow olfactory pit; *black* shallow inter-nasal groove; *white* hypoplastic *mf*; *lavender* hypoplastic *lf*; *orange and black* lack of choanal invagination; *light blue and black* lengthened oronasal groove; *light green and black* extended, shallow stomodeo-palatal line. **<u>Double-headed arrows</u>**: *red* inter medial frontonasal process width; *green* axis of orientation of main stomodeal ridges; *yellow* axis of orientation of ectodermal cell bodies. <u>(**A’, B’**)</u> Magnified fields outlined with white dashed-lined squares in **A** and **B**. Yellow lines outline cell boundaries in the superficial maxillary oral epithelium. Yellow double-headed arrow as in **A**. <u>(**C, D**)</u> Scanning electron micrographs of the developing palate and primary choanae of E12.5 *Foxg1^+/−^* and *Foxg1^−/−^* littermates. The lower jaws and tongues have been removed for clarity. *Doubleheaded orange arrow* relative breadth of the primary palate. *Red, white and orange arrow* indicates the naso-respiratory passageway that runs from the external nares, through the anterior nasal cavity, to exit at the primary choanae. **<u>Arrowheads</u>**: *red and black* underdeveloped, shallow external nasal aperture; *yellow* under- developed vibrissae field; *orange and black* lack of choanal invagination (leading to choanal atresia); *light green and black* under-elevated palatal shelf. <u>(**C’, D’**)</u> Magnified fields outlined with light green dashed-lined squares in **C** and **D**. *Light green lines* outline cell boundaries in the palatal periderm. **<u>Abbreviations</u>**: *en*, external nares; *lf*, lateral frontonasal process; *mf*, medial frontonasal process; *mx*, maxillary BA1; *nspr*, nasopharyngeal roof; *Rp*, Rathke’s pouch; *str*, stomodeal roof; *tel*, telencephalic dome; *ulp*, upper lip; *vib*, vibrissae; *1°c*, primary choanae; *1°p*, forming primary palate; *2°p*, forming secondary palate; λ, center of the lambdoidal junction.

SEM micrographs, moreover, revealed further morphological changes in E11.5 *Foxg1^−/−^* embryos undetected by light microscopy (Figures 4 A’, B’). Normally, the surface of the stomodeal roof of E11.5 embryos exhibits a fan-like sets of ridges radiating rostro-laterally from Rathke’s pouch to the mFNP and to where the internal choanae are beginning to invaginate (137, 138). However, E11.5 *Foxg1^−/−^* embryos showed no initiation of choanal invagination and exhibited fewer such ridges; additionally, those ridges that were present were un-associated with Rathke’s pouch, and abnormally ran medio-laterally. Moreover, because light microscopy suggested that the oral epithelium was thicker in *Foxg1^−/−^* embryos, we examined it at higher magnification. We found that the usual coordinated cellular directionality, size, and orientation characteristic of the epithelial surface of *Foxg1^+/−^* (and wild type) embryos was lacking in *Foxg1^−/−^* embryos: For instance, along the oral surface of mxBA1 the surface cells typically are of relatively uniform size and the long axis of the cells is nearly universally oriented to extend along the long axis of the arch (roughly parallel to the ridges noted above). This was not found to be the case in E11.5 mutant embryos as the cells of the oral surface of mutant mxBA1 varied widely in size and shape, and many cells of the mutant embryos were otherwise mis-oriented along the short axis of the arch.

Furthermore, in normal murine embryos from E11.5 to E12.5 the primary choanae typically invaginates where the developing primary and secondary palates laterally converge to generate antra on each side of the caudal end of the future nasal septum; eventually, the internal, deep epithelia of each antrum will make contact with the epithelia of the corresponding ventrocaudal invagination of the nasal cavity thus forming the oronasal (bucconasal) membrane. Rupture of this membrane thus permits complete passage from the external nares to the posterior respiratory system. Choanal atresia, effectively the developmental obstruction of the choanae, was clearly presaged in SEM micrographs of *Foxg1^−/−^* embryos at E12.5 (Figures 4C, D). Moreover, while both the mutant primary and secondary palatal primordia were present and positioned appropriately they were also both less clearly robust and dysmorphic. The trend toward a medio-laterally expanded stomodeal roof seen in younger mutant embryos was still patent with the broader primary palate and the laterally displaced upper lip primordia evinced in older embryos. Concordant with the abnormalities observed in the upper jaw skeleton, the leading edges of the secondary palate were seen to barely descend from the stomodeal roof in *Foxg1^−/−^* embryos.

Additionally, by E12.5 the murine oral epithelium has normally generated a specialized transient superficial cellular population, the periderm, thought to act as a protective barrier that inhibits potentially pathological adhesions between otherwise immature, adhesion-competent epithelia (139–143). SEM micrographs of the periderm of the rostral secondary palatal shelf of *Foxg1^−/−^* embryos at higher magnification showed an epithelial layer wherein the cellular size, directionality and orientation appears coordinated and uniform while SEM micrographs of comparable periderm of mutant embryos evinced cells that were distinctly flatter, less elongate, and less uniform in directionality (compare Figures 4C’, D’).

### Ontogenetic Elaboration of Expression Patterns of the λ-junction Associated Epithelial Signaling Factors *Bmp4*, *Fgf8, Raldh3* and *Shh* Is Altered in the SCE of *Foxg1^−/−^* Mutant Embryos

Development of the cranial skeleton is known to require an exquisitely timed and positioned molecular cross-talk between (and within) the embryonic cephalic epithelia and mesenchyme (6–10, 25, 30–33). This communication is manifest in the induction, maintenance, and elaboration of intricate patterns of gene expression, the spatial and temporal details of which underlie the precise translation of patterning processes and information into discrete, appropriate skeletal morphologies. Included in this craniofacial patterning dialogue are secreted signaling molecules (and their numerous cellular effectors) and master regulatory transcription factors. To address whether the defects noted in the maturation of the λ-junction and regional epithelia of *Foxg1^−/−^* embryos correlated with ontogenetic changes in the epithelial expression patterns of genes encoding signaling molecules (or their effectors) associated with cranial skeletal development, we first examined through whole mount *in situ* hybridization the expression patterns of *Bmp4*, *Fgf8*, *Raldh3* and *Shh* in *Foxg1^−/−^* and littermate embryos at E9-9.25 (time points when patterning within the SCE exhibits clear partitions, or compartments, as cephalic sensory placodes have been induced and discriminated) and E10.25-10.5 (the onset of λ-junction maturation when numerous significant essential patterning relations are topographically set).

We found that the loss of *Foxg1* fundamentally altered the topography of the expression patterns of these genes at these stages (Figure 5). For instance, at E9.25 *Bmp4* (a member of the large *Tgf-β* superfamily of signaling molecules) is dynamically expressed in the SCE of *Foxg1^+/−^* embryos, including at the presumptive center of the future λ-junction as well as in the distal mdBA1, the first pharyngeal plate, and epibranchial placodes, and the dorso-caudal optic primordia (Figures 5A-D) (48–50, 144–151). It is also expressed in cells in the region of the commissural plate (CP) that derives from the ANR. Within the forming λ-junction, *Bmp4* mRNA expression is typically detected in cells overlying the portion of mxBA1 growing toward the olfactory placode and in a dorsally projecting stripe delimiting optic SCE from olfactory SCE. In *Foxg1^−/−^* embryos, however, *Bmp4* was expressed more broadly in cells of the optic primordia, the forming λ-junction and the epibranchial placodes. Moreover, in *Foxg1^−/−^* mutants *Bmp4* expression in the dorsally projecting stripe associated with the λ-junction was stronger and extended further both dorsally and caudally toward the optic primordia. Additionally, precocious *Bmp4* expression was seen running through the SCE of the olfactory region and abnormally extensive expression was detected associated with the CP.

**Figures 5.**
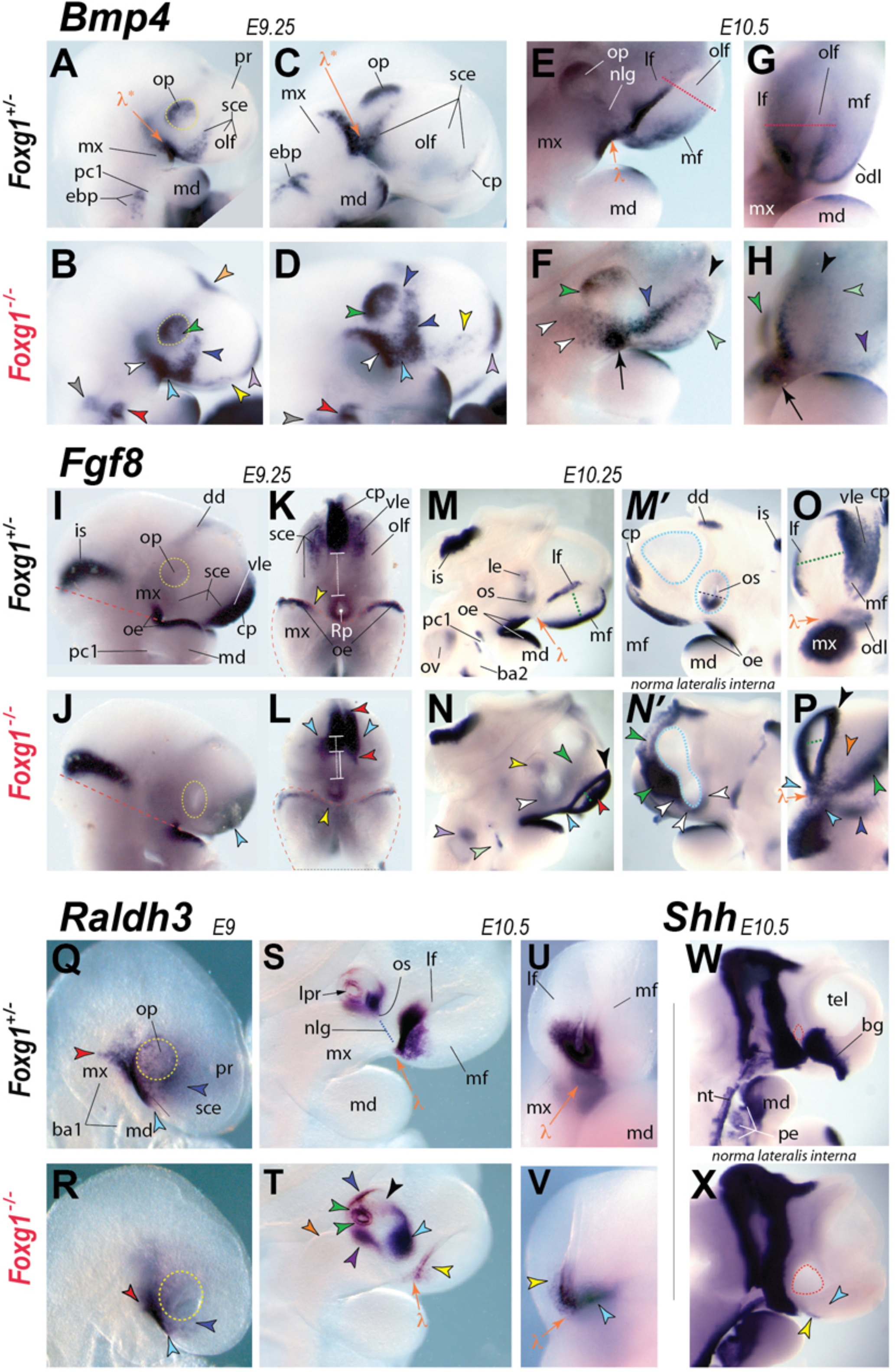
In situ hybridization reveals topographic alteration of regional signaling networks in the early SCE. <u>(**A-H**)</u> In situ hybridization of *Bmp4* in *Foxg1^+/−^* and *Foxg1^−/−^* littermates at E9.25 (A-D) and E10.5 (E-H). **<u>Dashed lines</u>*: yellow*** outline of optic primordia; ***red*** demarcation line between the normally high ventral frontonasal expression and low dorsal expression riming the olfactory pit. **<u>Arrowheads</u>**: ***salmon*** increased diencephalic signal; ***green*** increased signal in the optic ectoderm; ***white and black*** expanded post-optic maxillary signal associated with the future ectopic bulge; ***dark blue*** expanded pre-optic frontonasal signal; ***light blue*** intensified signal at center of future λ-lambdoidal junction; ***yellow*** signal in the olfactory sce; ***lavender*** commissural plate; ***red*** increased epibranchial expression; ***black*** increased, diffuse expression riming the dorsal olfactory pit; ***grey*** signal detected between the otic and epibranchial placodes; ***light green*** diffuse medial frontonasal signal; ***purple*** abnormally diffuse signa along the odontogenic line. <u>(**I-P**)</u> *In situ* hybridization of *Fgf8* in *Foxg1^+/−^* and *Foxg1^−/−^* littermates. (**I,J**) *Fgf8* expression at E9.25 viewed in norma lateralis (I,J) and norma basilaris externa showing the developing stomodeum after removal of post maxillary tissue (K, L). **<u>Lines</u>*: dashed yellow*** external outline of optic primordia; ***dashed red*** plane of dissection for K and L; ***dashed white*** distance from Rathke’s pouch expression of *Fgf8* to commissural expression in the heterozygote; ***solid white*** distance from Rathke’s pouch expression of *Fgf8* to commissural expression in the mutant. **<u>Arrowheads</u>**: ***light blue*** loss of detectable *Fgf8* transcripts in the sce; ***yellow*** relative medial extent of expression in the oral epithelium; ***red*** relative compaction of the commissural plate toward Rathke’s pouch. <u>(**M-P**)</u> *Fgf8* expression at E10.25 viewed in norma lateralis externa (M, N), norma lateralis interna (M’, N’) and norma frontalis (O, P). **<u>Dashed lines</u>: *green*** relative expansion of expression into the olfactory pit; ***light blue*** outline of prosencephalic vesicles. **<u>Arrowheads</u>**: ***light blue*** ectopic expression at the center of the λ-junction connecting frontonasal and mxBA1 expression; ***white*** ventral optic stalk; ***red*** expanded expression in the medial olfactory pit; ***yellow*** diffuse, expanded expression encompassing the lens; ***orange*** loss of expression in the vle; ***green*** anterior forebrain expression expanded both ventrally and dorsally; ***lavender*** topographic alteration associated with the ov; ***light green*** loss of expression associated with the ba2 pharyngeal cleft; ***dark blue*** change of the presumptive odl. <u>(**Q-V**)</u> *In situ* hybridization of *Raldh3* in *Foxg1^+/−^* and *Foxg1^−/−^* littermates at E9 (Q, R) and E10.5 (S-V). (Q, R) **<u>Dashed lines</u>: *yellow*** outline of optic primordia. **<u>Arrowheads</u>**: ***light blue*** distal extent of expression in mdBA1; ***red*** proximocaudal extent of expression in mxBA1; dark blue pre-optic sce. (S-V) **<u>Dashed lines</u>: *blue*** axis of the nasolacrimal groove. **<u>Arrowheads</u>**: ***orange*** ectopic bulge in mxBA1; ***yellow*** significant loss of expression in the ventral olfactory pit at the λ-junction; ***light blue*** expanded expression at the ventral nasal optic cup; ***black*** swath of positive cells across the optic ectoderm to the lens pit; ***green*** increased expression associated with the rim of the lens pit; ***blue*** increased expression dorso-templar optic primordia; ***purple*** increased expression in the ventral temporal optic cup. <u>(**W, X**)</u> *In situ* hybridization of *Shh* in hemisected *Foxg1^+/^*and *Foxg1^-/^*littermates at E10.5 viewed in norma lateralis interna. **<u>Dashed lines</u>: *red*** position of the optic stalk. **<u>Arrowheads</u>**: ***light blue*** absence of expression corresponding to loss of basal ganglial structure; ***yellow*** ectopic expression. **<u>Abbreviations</u>**: ***ba1***, first (mandibular) branchial arch; ***bg***, basal ganglia; ***cp***, commissural plate; ***dd***, diencephalon; ***ebp***, epibranchial placodes; ***exn***, external nares; ***is***, isthmus; ***lf***, lateral frontonasal process; ***lpr***, lens pit rim; ***md***, mandibular branch of the first branchial arch; ***mf***, medial frontonasal process; ***mx***, maxillary branch of the first branchial arch; ***ne***, neuroectoderm; ***odl***, odontogenic line; ***oe***, oral epithelium; ***olf***, olfactory placode or pit; ***op***, optic primordia; ***os***, optic stalk; ***pc1***; first pharyngeal cleft; ***pe***, pharyngeal epithelium; ***pr***, prosencephalon; ***Rp***, Rathke’s pouch; ***sce***, surface cephalic ectoderm; ***vle***, ventrolateral ectoderm; **λ**, center of the lambdoidal junction; **λ\***, future lambdoidal junction.

Aberrant *Bmp4* expression was also seen associated with the mature λ-junction of E10.5 *Foxg1^−/−^* embryos. Normally, strong but tightly restricted and focalized *Bmp4* expression is found in the epithelia of the distal tip of mxBA1 and along the ventral rims of the invaginating olfactory pits, in a small swath of cells on the lFNP directed toward the eye, and in the odontogenic line paralleling the naso-stomodeal line along the ventral mFNP. At this embryonic stage, lighter *Bmp4* mRNA expression is also typically detected in the dorsal rims of the olfactory pit, the cells along the future nasolacrimal groove, and the dorso-caudal optic primordia. In E10.5 mutant embryos, *Bmp4* expression was found to be strong at, but not tightly restricted within, the λ-junction. Evidence of greater levels of *Bmp4* mRNA in mutant embryos was seen along the dorsal olfactory pits and lower levels seen associated with the ventral mFNP. Noticeably, expression associated with mxBA1 differed greatly between heterozygous and homozygous mutant embryos. For instance, rather than being expressed in cells associated with the nasolacrimal line, ectopic *Bmp4* transcripts were detected in a broad swath of SCE cells running ventral to the eye and toward the ectopic, sub-optic maxillary bulge (described above).

*Fgf8*, a developmentally active and potent fibroblast growth factor, is essential for the normal development of the jaw (152–157) as well as the OPC (158) and NC (158–162), and ontogenetic disruption of its expression characterized E9.25 and E10.25 *Foxg1^−/−^* embryos (Figures 5I-P). In addition to the isthmic organizer, at E9.25 *Fgf8* mRNA is normally highly detectable in the oral ectoderm of BA1 and the SCE overlying the developing CP. Notably, *Fgf8* is also typically expressed in a more diffuse manner in the ventrolateral ectoderm (VLE), a domain of the SCE that lies between the CP and the forming olfactory placode. At E9.25, *Foxg1^−/−^* embryos exhibit significant loss of detectable *Fgf8* mRNA in the VLE just lateral to the CP. Expression in the CP itself was maintained in mutant embryos but did not extend as far dorso- ventrally within the CP as in *Foxg1^+/−^* embryos. Moreover, low levels of *Fgf8* transcripts were also detected extending further medially from the mxBA1 oral epithelium toward Rathke’s pouch in mutant embryos.

An embryonic day later, the *Fgf8* expression profile of the λ-junction in *Foxg1^−/−^* mutants continued to be anomalous (Figures 5I-P). In wild type embryos at E10.25 *Fgf8* mRNA is clearly detected in the isthmic organizer, dorsal diencephalon, and the ventral optic stalk. It is also found in the oral epithelium of mdBA1 and mxBA1 as well as in the outer-margins of the dorsal epithelium of both mFNP and lFNP that rims of the olfactory pit; at this stage, however, *Fgf8* is normally distinctly absent from the center of the λ-junction where mxBA1 and the FNP converge. Transcripts of *Fgf8* also continue to be expressed in a diffuse manner in the VLE of the mFNP. In mutant embryos, however significant *Fgf8* mRNA expression was found abnormally associated with the center of the λ-junction and cells rimming the entire olfactory pit. Within the mutant pits, *Fgf8* was aberrantly expressed in the medial wall where the primordial cells of the VNO are thought to arise. *Foxg1^−/−^* mutants also failed to express *Fgf8* in the VLE although they did ectopically express it rostrally and dorsally throughout the NE from the malformed ventral optic stalk to the dorsal diencephalon.

Retinoic acid (RA), another potent signaling and patterning molecule, is a lipid soluble morphogen generated as an active derivative of vitamin A and its developmental regulation must be tightly controlled (148–150, 163–168). *Raldh3* is a gene encoding a retinaldehyde dehydrogenase involved in the second oxidative step in RA synthesis, and it is focally expressed at the heart of the developing λ-junction in murine embryos from E9 to E10.5 (Figures 5Q-V). At E9.0, *Raldh3* mRNA is normally found within the SCE associated with the optic primordia, the distal mxBA1, and the ventral olfactory pit of both the mFNP and lFNP. By E10.5, *Raldh3* is conspicuously, though asymmetrically, expressed along the rim of the invaginated lens and optic cup as well as in the epithelium of the ventral-most invaginated olfactory pit. However, in E9.0 *Foxg1^−/−^* embryos *Raldh3* expression was not as extensive in the SCE associated with the optic primordia or mxBA1. By E10.5, a significant reduction in olfactory *Raldh3* expression was visible in mutant embryos. Moreover, while expression associated with the dysmorphic eye was more intense, including in the cells of the lens and optic cup, the asymmetric nature of *Raldh3* expression was maintained in mutant embryos.

*Shh* encodes another secreted protein, one with autoproteolytic activity that, in conjunction with covalent addition of cholesterol, produces both long and short-range morphogens (169–171). It is embryonically expressed in many ventral, midline tissues including the node, head process, notochord, and floor plate and basal ganglia of ventral forebrain, and its loss leads to dramatic holoprosencephaly (150, 172–177). It has been previously reported that, because *Foxg1* mutants lack a proper ventral forebrain, they also lack *Shh* in this region (105). We confirmed the absence of *Shh* in the ventral forebrains of E10.5 *Foxg1^−/−^* embryos (Figures 5W, X); we, noted, however that lower levels of *Shh* expression appeared in epithelium associated with the stomodeal roof at both E9 and E10.5 (**Supplementary Figure XX Shh**).

### Further Evidence of Impaired Molecular Maturation in and around the λ-junction of *Foxg1^−/−^* Mutant Embryos

In addition to epithelial signals, the mature λ-junction at E10.5 is a source of signaling molecules originating from the CNC ectomesenchyme. This includes production of Wnt5a, a protein that signals through non-canonical Wnt pathways (178–182). Typically, *Wnt5a* expression at E10.5 is strong within the mFNP, the lFNP and the portion of mxBA1 associated with the FNP at the center of the λ-junction, while weaker expression is encountered in the distal mdBA1 (Figure 6). In *Foxg1^−/−^* embryos, *Wnt5a* mRNA was detected in the λ-junction; however, *Wnt5a* expression was weaker than normal in mxBA1 but stronger than normal throughout the FNP and olfactory pit. Moreover, a noticeable gap in expression between the FNP and mxBA1 was encountered in *Foxg1^−/−^* mutants. Thus, we found the topographic elaboration of essential craniofacial patterning genes in both the embryonic cephalic ectoderm and mesenchyme to be significantly altered in *Foxg1^−/−^* embryos from E9 through E10.5.

**Figure 6:**
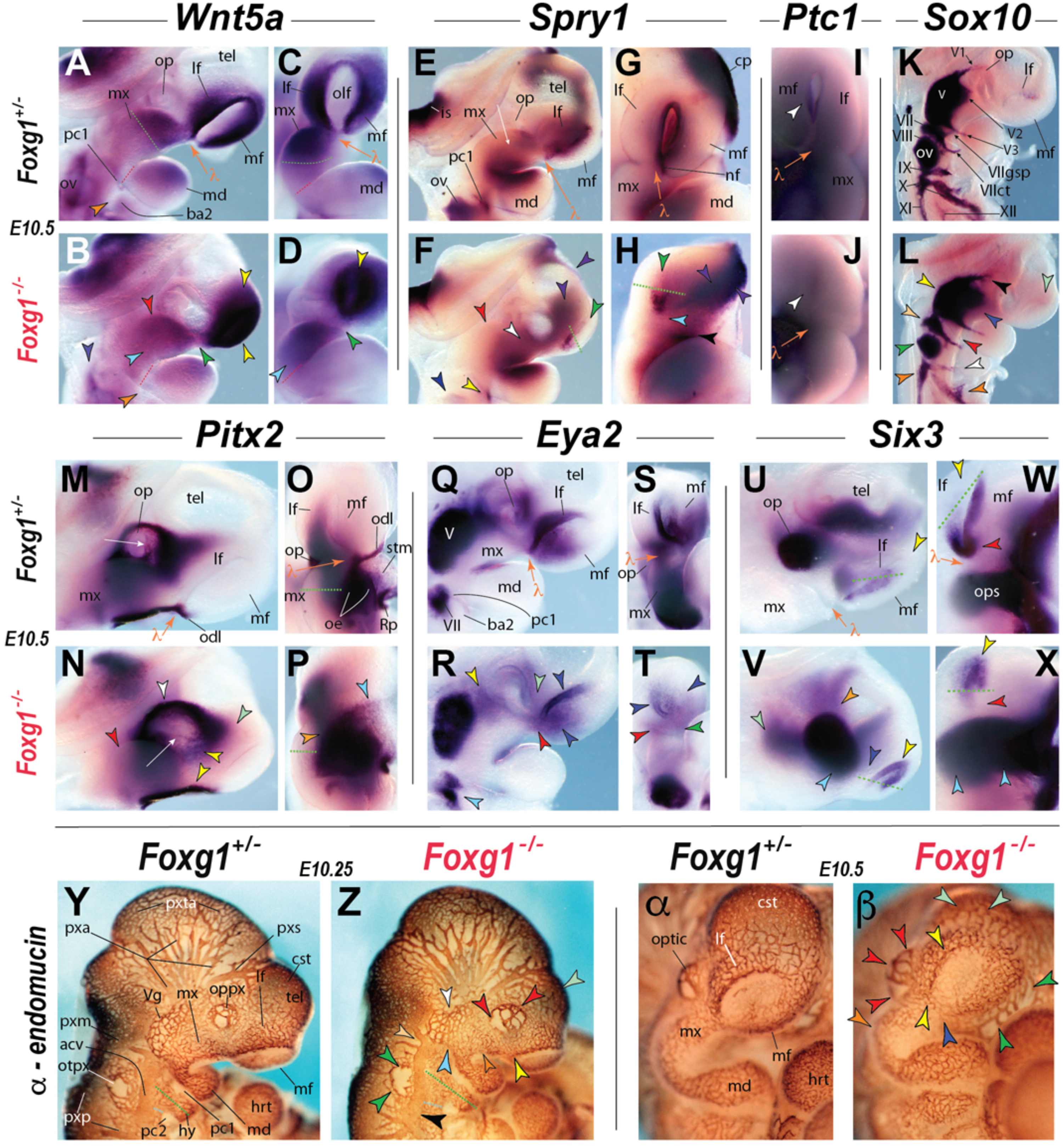
Impaired Cephalic Molecular Maturation and Vascular Patterning. (**A-X**) *In situ* hybridization of *Foxg1^+/−^* and *Foxg1^−/−^* littermates at E10.5. <u>(**A-D**</u>) *Wnt5a*. **<u>Dashed lines</u>: *green*** line demarcating position of normally higher level expression in mxBA1; ***red*** boundary of proximal (mx) and distal (md) BA1 coinciding with a hinge-centric expression free zone. **<u>Arrowheads</u>**: ***yellow*** increased expression associated with the olfactory pit and frontonasal processes; ***red*** position of the ectopic bulge; ***light blue*** expansion of higher expression within the mxBA1; ***dark blue*** change in transcript intensity associated with he facial ganglion and ov; ***green*** alteration of expression at the center of the λ-junction including uniting the mf and lf and separating the latter from mxBA1; ***orange*** hinge-centric expression free zone separating proximal and distal ba1. <u>(**E-H**)</u> *Spry1*. **<u>Dashed lines</u>: *green*** abnormal restriction of expression to the ventral olf. **<u>Arrowheads</u>**: ***light blue*** absence of expression along the presumptive nasal fin; ***red*** position of the ectopic bulge in mxBA1; ***green*** absence of detectable transcript in the dorsal olf; ***purple*** forebrain, including commissural plate and optic stalk; ***white*** changing expression between that in the oral epithelium and junction-associated mxBA1; ***black*** stomodeal expression; ***yellow*** first pharyngeal cleft; ***dark blue*** ov. <u>(**I, J**)</u> *Pct1*. ***White*** arrowhead indicates changes associated with the olf. <u>(**K, L**)</u> *Sox10*. **<u>Arrowheads</u>**: ***light green*** loss in the olf; ***black*** changing branching and extension in V1; ***dark blue*** aberrant shape and orientation of V2; ***yellow*** trigeminal ganglion; ***salmon*** dorsal facial nerve ganglion; ***red*** loss of the greater superior petrosal and truncation of the chorda tympani of VII; ***green*** change associated with the ov; ***white*** altered glossopharyngeal staining; ***orange*** altered vagal nerve staining. <u>(**M-P**)</u> *Pitx2*. Dorsal expansion and ventral loss of expression in peri-ocular mesenchyme and altered corneal wave of migration. ***White arrows*** indicate transcripts in the wave of corneal mesenchyme (CNC) between the lens and the SCE. ***Dashed green line*** width of transcript zone in the oral ectoderm of mxBA1. **<u>Arrowheads</u>**: ***light green*** loss in the pre-optic mesenchyme extending to the lFNP; ***white*** increased supra-orbital mesenchymal signal; ***yellow*** loss of signal associated with the mesenchyme and epithelia associated with the nasolacrimal grove; ***orange*** lateral expansion of signal in the maxillary oral ectoderm; ***light blue*** loss of definitive odontgenic line expression; ***red*** ectopic bulge in mxBA1. <u>(**Q-T**)</u> *Eya2*. **<u>Arrowheads</u>**: ***light green*** loss associated with the rostral optic primordia; ***dark blue*** decreased expression in he ventral olf; ***green*** diminished expression in the ventral mf; ***red*** loss of expression associated with he nasolacrimal groove and the center of the λ-junction; ***yellow*** decreased expression between the original ganglion and the optic primordia; ***light blue*** greater separation between the oral and aboral expression in BA2. <u>(**U-X**)</u> *Six3*. Significant alteration of expression in the forebrain, optic primordia, and olfactory pit. **<u>Dashed lines</u>: *green*** demarcation of ventral-dorsal of expression highlighting the reversed polarity of expression within the *olf*. **<u>Arrowheads</u>**: ***yellow*** dorsal rim of the olf; ***red*** ventral olf; ***dark blue*** loss of expression with the ventral interior of the olf; ***light blue*** increased expression in the optic associated forebrain; ***orange*** loss in the hypomorphic telencephalon; ***light green*** increased post optic cephalic expression. <u>(**Y-β**)</u> Anti-endomucin immunoreactivity revealing aberrant cephalic vascular development at E10.25. **<u>Dashed lines</u>: *green*** relative distance from the distal BA2 to the acv; ***light blue*** relative distance from the vessel free pc2 domain to the acv. **<u>Arrowheads</u>**: ***yellow*** vessel development associated with the λ-junction and lf; ***red*** developing optic plexus; ***light green*** capillary sheet of the telencephalon; ***white*** diminished vascularization at the trigeminal ganglion; ***light blue*** aberrant development associated with the hinge region; orange mxBA1; ***salmon*** sinus to the plexus medals; ***green*** under-development of the otic plexus; ***black*** decreased width of the acv. **<u>Abbreviations</u>**: ***acv***, anterior cardinal vein (vena wapitis prima); ***ba2***, second branchial arch; ***cst***, capillary sheet, telencephalon; ***hrt***, heart; ***is***, isthmus; ***lf***, lateral frontonasal process; ***md***, mandibular branch, first branchial arch; ***mf***, medial frontonasal process; ***mx***, maxillary branch, first branchial arch; ***nf*** nasal fin; ***odl***, odontogenic line; ***olf***, olfactory pit; ***op***, optic primordia; ***oppx***, embryonic optic plexus; ***otpx***, embryonic otic plexus; ***ov***, otic vesicle; ***pc1***, first pharyngeal cleft; ***pc2***, second pharyngeal cleft; ***pxa***, plexus anterior; ***pxm***, plexus medals; ***pxp***, plexus posterior; ***pxs***, plexus sagittalis; ***pxta***, plexus tentorii anterior; ***tel***, telencephalon; ***V1***, ophthalmic branch of trigeminal cranial nerve; ***V2***, maxillary branch of trigeminal cranial nerve; ***V3***, mandibular branch of cranial trigeminal nerve; ***VII***, facial cranial nerve; ***VIIct***, chorda tympani, facial cranial nerve; ***VIIgsp***, greater superior petrosal, facial cranial nerve; ***VIII***, vestibuloacoustic nerve; ***IX***, glossopharyngeal nerve; ***X***, vagal nerve; ***XI***, spinal accessory nerve; ***XII***, hypoglossal nerve; **λ**, center of the lambdoidal junction.

Because, in the normal course of epithelial-mesenchymal cross-talk, cell and tissue competence to signal is typically coordinated with cell and tissue competence to respond, we examined the cephalic gene expression patterns of early response and obligatory downstream effectors of signaling and compared the expression topographies of presumed molecular signal and response. Herein we highlight alterations detected in *Foxg1^−/−^* embryos of the expression in two such genes: *Spry1*, an early Fgf8-responsive gene and mediator of Fgf8 signaling (183), and *Ptc1*, an effector of Hedgehog signaling (184). As one might expect for an early responsive gene, *Spry1* expression in E10.5 murine embryos generally follows that of *Fgf8*: thus, in *Foxg1* heterozygotes it was expressed at the isthmic organizer, the dorsal diencephalon, the CP, the BA1 oral ectoderm, the pharyngeal plates and rimming the entirety of the olfactory pit and within the nasal fin (Figure 6E, G). Though we did not discriminate this in our whole mount *in situ* hybridization assays, low levels of *Spry1* have also been reported to be expressed in the lens and corneal epithelium where it is necessary for proper separation of the lens vesicle from the SCE and the formation of eyelids (185, 186). We found that in most tissues *Spry1* expression in E10.5 *Foxg1^−/−^* embryos followed that of *Fgf8* expression (Figures 6F, H). For instance, mutant embryos expressed *Spry1* in the oral epithelium and in an expanded manner associated with the expanded *Fgf8* expression in the dysmorphic rostral brain (including the optic sulcus). Notably, however, detection of *Spry1* mRNA in the mutant FNP was restricted to low-levels at the ventral end of the olfactory pit and did not mirror the strong ectopic expression of *Fgf8* at the center of the λ-junction and along the olfactory pit. Thus, unlike in other embryonic regions the topography of *Fgf8* expression associated with the λ-junction was not followed by similar topography in the expression of this Fgf8 responsive gene. Moreover, as with *Fgf8* and *Spry1*, *Ptc1* mRNA is typically expressed in the olfactory pits associated with mature λ-junctions (Figure 6). *Ptc1* acts, in addition to being as an effector of Hedgehog signaling through its role as a cell-surface receptor, as an early responsive gene to Shh signaling (184). We found its expression in the olfactory pit at E10.5 abrogated in embryos lacking functional *Foxg1*.

The evident ontogenetic perturbations of signaling gene expression exhibited in *Foxg1^−/−^* embryos prompted us to further investigate the molecular elaboration of the mature λ-junction in mutant embryos. We proceeded by characterizing the patterns of expression of genes and proteins implicated in disparate aspects of normal, regionally relevant cranial development, including *Sox10, Pitx2, Eya2, Six3*, and endomucin (an early marker of forming blood vessels).

*Sox10* is expressed in CNC as they delaminate and then migrate from the neural tube-ectodermal boundary (187–191). By E10.5, the CNC have already ceased their migration and are executing their responses to local developmental cues; at this point, *Sox10* mRNA is found associated with the cranial nerve ganglia and the presumptive olfactory ensheathing cells (OEC) within the mesenchyme of the olfactory pit (188, 192–194). The cranial ganglia are composite structures formed of both CNC and cells that have delaminated from the particular neurogenic placode; this process of gangliogenesis requires tightly controlled epithelial-CNC interactions and proper expression of *Sox10* during this process occurs in the context of a normally developed epithelial-mesenchymal interface. SOX10 has also been implicated in Kallman syndrome wherein a disruption occurs in the establishment of the routing of neuroendocrine gonadotropin-releasing hormone (GnRH) cells along the vomeronasal and terminal nerve fibers in the peripheral olfactory system to their final destination in the preoptic and hypothalamic region of the forebrain (195). It is also expressed in the otic vesicle (196, 197; Figure 6K). In E10.5 *Foxg1^−/−^* embryos we found that (where expressed) *Sox10* mRNA was strongly detectable; however, the pattern of *Sox10* expression was topographically altered (Figure 6L). For instance, we failed to detect *Sox10* mRNA in the FNP. Staining within the trigeminal ganglia, moreover, revealed both an abnormal shape to this structure and abnormality of ophthalmic branching and extension toward and above the optic primordia. Additionally, two projections of the geniculate (facial) ganglia normally detectable with *Sox10* expression, the chorda tympani and the greater superior petrosal, were either not observed (greater superior petrosal) or were abnormal in projection (chorda tympani) in mutant embryos. The development and course of both of these facial nerve branches is tied to structures deeply altered by the absence of *Foxg1*: the greater superior petrosal nerve with the regio pterygopalatina. Therafter with the lacrimal gland as well as the mucosal glands of the nose, palate, and pharynx; and, the chorda tympani with the inner and middle ear prior to its targeting the tongue. In addition to showing disrupted expression levels within the epibranchial ganglia, *Sox10* mRNA expression in mutant embryos highlight the abnormal shape in the otic vesicle seen in *Foxg1* mutant embryos at this time. *Sox10* expression, moreover, was undetected at E10.5 in mutant embryos in the mesenchyme of the olfactory pit.

Mice harboring mutations of *Pitx2* have served as a model for Axenfeld–Rieger syndrome, a congenital condition characterized by defects of optic and craniofacial facial development (198, 199). *Pitx2* is expressed at E10.5 in both the CNC and mesodermal components of the POM, the oral epithelium (including along the odontogenic line) and Rathke’s pouch (Figures 6M, O) (200–206). In *Foxg1^−/−^* embryos, *Pitx2* expression was expanded supra-optically but was constricted in the mxBA1 and anterior in the lFNP (Figures 6N, P). *Pitx2* expression in the nasolacrimal groove was abrogated in mutant embryos, while the normal anterior expression in the migratory wave of nascent corneal mesenchyme appeared to have been shifted ventrally toward the groove. Additionally, the discrete, intense band of *Pitx2* normally found along the odontogenic line of the mFNP of E10.5 wild type and heterozygous embryos was not evinced in embryos lacking *Foxg1*. Moreover, higher levels of *Pitx2* expression were seen extending medially from the λ-junction to cover more of the stomodeal epithelium in *Foxg1^−/−^* mutant embryos.

The SCE expresses numerous additional factors associated with cranial dysmorphology syndromes, including several *Eya* genes known to be essential for proper cranial development: The loss of Eya proteins has been linked with craniofacial defects and hearing loss characteristic of branchio-oto-renal (BOR) syndrome (207–211). *Eya2* is normally expressed in E10.5 murine embryos in the cranial ganglia, around the optic primordia, in the FNP and the ventral lining the olfactory pit associated with the λ-junction, as well as in the nasolacrimal groove and oral epithelium of mxBA1 (Figures 6Q, S). In *Foxg1^−/−^* embryos significantly less *Eya2* transcripts were detected in the olfactory pit, FNP, mxBA1, nasolacrimal groove and POM (Figures 6R, T).

Loss of *Six3* (a sine oculis-related homeobox transcription factor gene) in mice results in significant tissue loss in the forebrain as well as structures associated with the SCE and λ-junction (212–217). *Six3* heterozygosity has also been implicated in anosmia-based decreased male fertility due to disruption of GnRH cells migration stemming from defective olfactory axon targeting (218). *Six3* transcripts at E10.5 are typically detected within the optic primordia, including the optic stalk, the ventral telencephalon and the ventral olfactory pit epithelium close to the center of the λ-junction (Figures 6U, W). We found that the dysmorphic telencephalon and optic stalk of *Foxg1^−/−^* mutants also expressed *Six3* (Figures 6V, X). We also found that *Six3* continued to be expressed in the olfactory pits of mutant E10.5 embryos; notably, however, the normal dorso- ventral polarity of *Six3* expression was inverted in mutant embryos as *Six3* was absent ventrally but was ectopically present dorsally within the olfactory pits, a clear indication that the normal patterned polarity of the olfactory-related SCE had been corrupted by the loss of *Foxg1* (Figures 6W, X).

Because development of the nascent embryonic vascular plexus and its remodeling is believed to be sensitive to the local molecular environment (219–221), we assayed cephalic vascular development in E10.25 *Foxg1^+/−^* and *Foxg1^−/−^* embryos through whole mount anti-endomucin immunostaining. This revealed that while vasculogenesis and subsequent angiogenesis were not globally impaired in mutant embryos, there were regional changes in the cephalic vascular network (Figures 6Y-β). For instance, the mutant otic plexus was underdeveloped, with an expanded vascular-free zone, and was closely apposed to the dorsally-shifted anterior cardinal vein (Figure 6Z). The optic plexus was both expanded and disorganized, while the trigeminal plexus was smaller and constricted. Moreover, the vascularization of the FNP rimming the olfactory pit was more chaotic, the typically tight capillary sheet covering the telencephalon failed to manifest as such, and a larger region of the midline stomodeal roof remained largely vessel free in *Foxg1^−/−^* embryos.

### Loss of *Foxg1* Significantly Alters SCE Compartmentalization and The Topography of Apoptosis

Because our previous analysis had demonstrated (1) that the cellular organization of the SCE in *Foxg1^−/−^* embryos was altered by E11.5, (2) that mutant embryos failed to appropriately manifest a molecularly and structurally mature λ-junction by E10.5, and (3) that the expression patterns of *Bmp4*, *Fgf8*, *Raldh3* and *Shh* were patently aberrant in the SCE of mutant embryos by E9.25, we extended our molecular and cellular analysis of the early (E9.25) SCE (Figure 7). We were initially interested in addressing whether the topographic molecular and cellular compartmentalization in the early λ-junction associated SCE was likewise already compromised by the loss of *Foxg1* between E9 and E9.5; in particular, as a read-out of compartmentalization of the SCE, we were interested both in the expressional topography of transcription factors associated with olfactory placode and capsular development, and in whether the mis-coordination of cellular directionality, size, and orientation characteristic of E11.5 mutant ectoderm was already evident just as the CNC had just reached it post-migratory destinations within the developing head.

**Figure 7:**
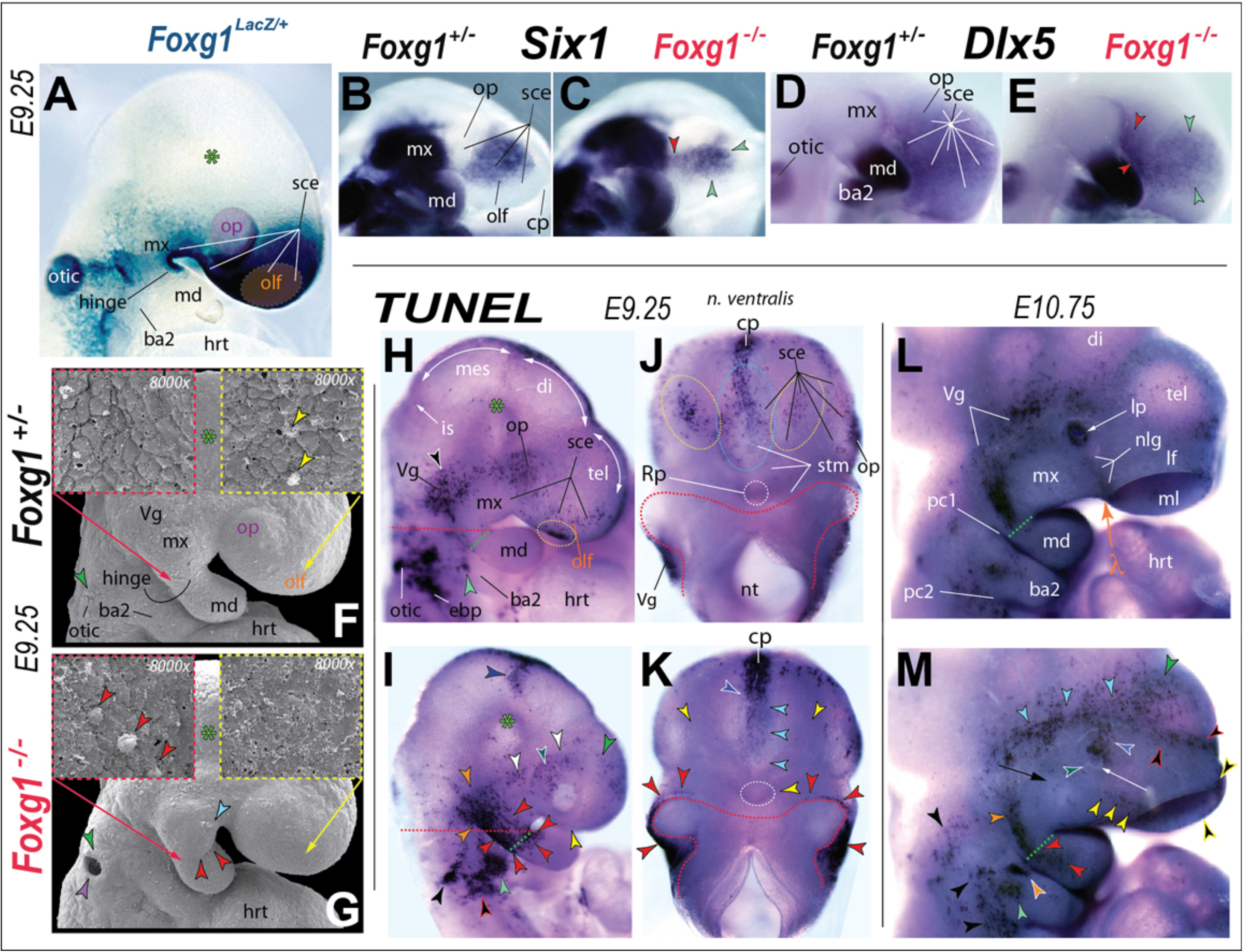
Loss of SCE compartmentalization and Drastically Altered Topography of Cephalic Apoptosis is Patent by E9.25 in *Foxg1*^−/−^ mutants. <u>(**A**)</u> Cephalic expression of *Foxg1* as revealed by β-galactosidase enzymatic activity in an E9.25 *Foxg1^LacZ/+^* embryo. (**<u>B-E</u>**) *In situ* hybridization of *Six1* (B, C) and *Dlx5* (D, E) in *Foxg1^+/−^* and *Foxg1^−/−^* littermates at E9.25. *Green arrowheads* highlight the rostro-medial and dorsal restriction of expression in the olfactory SCE. The *red arrowhead* highlights the de-compartmentalization of the *Six1* and *Dlx5* expression fields as evidenced by the ectopic, sub-optic expansion of olfactory ectodermal expression across the SCE and into the maxillary field. <u>(**F, G**)</u> Scanning electron micrographs of E9.25 *Foxg1^+/−^* (F) and *Foxg1^−/−^* (G) littermates demonstrating altered cephalic morphology and changes in the topographic profile of apoptotic epithelial cells associated with *Foxg1* expression domains in the SCE. *Green asterick* indicates the position of *Foxg1* negative ectoderm used as a control for cephalic epithelial appearance. <u>Arrowheads</u>: *yellow* cellular extrusions of the olfactory (yellow arrow) region SCE characteristic of apoptotic ectodermal cells; *red* cellular extrusions of aboral maxillary hinge region (red arrow); *light blue* change of morphology of the mxBA1; *green* distinct, tufted cellular cluster typical of the rostro-dorsal otic pit and early vesicle; *purple* delayed closure of the otic pit to form an enclosed vesicle. <u>(**H-K**)</u> Whole mount TUNEL assay of apoptosis in E9.25 *Foxg1^+/−^* (H, J) and *Foxg1^−/−^* (I, K) littermates confirming drastic alterations of the topography of cephalic ectodermal apoptotic cell death. **<u>Dashed lines</u>**: *red* line of resection to view the stomodeal ectoderm; *white* forming Rathke’s pouch. <u>Arrowheads</u>: *black* altered profile in topography and extent of otic region apoptosis; *blue and black* increased numbers of apoptotic cells overlying the mesencehalon-diencephalon boundary; *green* increased dorsal telencephalic cell death; *green and white* fewer apoptotic cells in the lens-associated SCE; *light blue* decreased apoptosis in the midline stomodeal ectoderm ventro-caudal to the intense cortical plate expression; *light green* increased, intense apoptotic cellular clustering in proximal BA2 in axial registration with the hinge region; *orange* change in the position and extent of cell death in the forming trigeminal ganglion; *red* intense clustering of apoptotic cells in the hinge-centered oral, lateral, and peri-cleftal (aboral,) mxBA1 SCE associated with the hinge with some extension into the presumptive proximal mdBA1 field; *red and black* aberrant topography and extent of ebibranchial ectodermal apoptosis; *white* peri-optical ectodermal apoptosis re-oriented toward the telencephalon; *white and blue* ventral extension of the intense apoptosis associated with the cortical plate; *yellow* loss of apoptotic cells in the olfactory-associated SCE. <u>(**L, M**)</u> Whole mount TUNEL assay of apoptosis in E10.75 *Foxg1^+/−^* and *Foxg1^−/−^* littermates. The *black arrow* indicates the relative position of the ectopic bulge while the *white arrow* points to the ventral lens pit. <u>Arrowheads</u>: *black* increased numbers of otic and ganglion-associated dying cells; *green* increased cell death overlying the telencephalic dome; *green and white* abnormal absence of apoptotic cells in the ventral lens pit; *light blue* highly apoptotic supra-orbital region connecting the trigeminal and telencephalon dome; *light green* increased cell death in the lateral patch in the proximal BA2 in typical axial registration with the hinge region; *orange* slight decrease in the hinge region cell death typically associated with the dorso-caudalcaudal, supra-cleftal mxBA1; *red* ectopic clustering of apoptotic cells in mdBA1 in line with the dorso-caudalcaudal, supra-cleftal mxBA1SCE apoptotic cells; *red and black* ectopic apoptotic cells lining the frontonasal telencephalon groove; *white and blue* ectopic, intensely stained cluster of apoptotic cells extending dorsally from the dorsal lens pit; *white and green* loss of apoptosis within the cells of the ventral lens pit; *white and orange* intensely stained ectopic cluster of apoptotic cells between the lateral patch and the first pharyngeal cleft; *yellow* re-orientation of directionality of the apoptotic cells associated with the nasolacrimal groove toward the ectopic mxBA1 bulge; *yellow and black* greater levels of apoptotic cells in the the dorsal and medial rims of the olfactory pits. **Abbreviations**: *ba2*, second branchial arch; *cp*, commissural plate; *di*, diencephalon; *ebp*, epibranchial placodes; *hrt*, heart; *is*, isthmus; *lf*, lateral frontonasal process; *lp*, lens pit; *md*, mandibular BA1; *mes*, mesencephalon; *mf*, medial frontonasal process; *mx*, maxillary BA1; *nlg*, nasolacrimal groove; *nt*, neural tube; *olf*, olfactory sce; *op* external optic primordia; *otic*, otic primordia; *pc1*, first pharyngeal cleft; *pc2*, second pharyngeal cleft; *Rp*, Rathke’s pouch; *sce*, surface cephalic ectoderm; *stm*, stomodeum; *tel*, telencephalon; *Vg*, trigeminal ganglion; λ, center of the lambdoidal junction.

Partition and refinement of the olfactory placode within the SCE requires the early restriction and compartmentalization of combinations of transcription factors within the SCE, including of *Six1, Six3* and *Dlx5* (222–226). We therefore first compared *Six1* expression in the SCE of *Foxg1^+/−^* and *Foxg1^−/−^* embryos at E9.25, (when restriction of *Six1* should be in effect). We found that, while mutant olfactory SCE expression of *Six1* in mutant embryos was abnormally restricted dorsally and rostro-medially, it was also ectopically extended in the SCE below the optic placode toward the ectoderm of the mxBA1 (Figures 7B, C). Additionally, we noted that expression of *Six3* was likewise attenuated in the presumptive olfactory SCE of E9.25 *Foxg1^−/−^* embryos (data not shown). *Dlx* genes have also been implicated in the elaboration of placodal SCE, and (like *Six1*) *Dlx5* is an essential factor for the proper development of NC structures (6, 7, 80, 227). *In situ* hybridization of *Dlx5* revealed that its SCE expression was maintained at E9.25 in the absence of *Foxg1*. Similar to that of *Six1*, *Dlx5* expression in *Foxg1^−/−^* embryos likewise also appeared stronger in the SCE of mutant embryos below the lens placode (Figures 7D, E) while that in the CNC of mdBA1 remained unchanged. Thus we concluded that the topography of compartmentalization within the E9.25 SCE was altered by the loss of *Foxg1*, and taken together with the abnormal early (E9-E9.25) expression patterns of *Bmp4*, *Fgf8*, and *Raldh3* these results demonstrate that the normally *Foxg1*-positive SCE of *Foxg1^−/−^* embryos was already molecularly compromised by E9.25.

To initiate an investigation of the temporal origins of the mis-coordination of cellular directionality, size and orientation in the SCE evinced in *Foxg1^−/−^* embryos at E11.5, we examined mutant and heterozygous E9.25 littermate embryos through scanning electron microscopy (Figures 7F, G). At low magnification, a number of distinctions between *Foxg1^+/−^* and *Foxg1^−/−^* embryos were evident: Among these were the delay in the closure of the otic pit and clear morphological changes at the mxBA1-mdBA1 boundary evident in *Foxg1^−/−^* embryos (Figures 7F, G). To assess the SCE cellular organization at higher magnification, we first compared the *Foxg1*-negative cephalic ectoderm lateral to the mesencephalon-diencephalon boundary at the bend of the cephalic flexure (positionaly denoted by the green asterisks, Figure 7), and found this ectoderm in *Foxg1^+/−^* and *Foxg1^−/−^* embryos to be similarly characterized by ciliated, flattened cells bound by numerous microvilli. Thus, we found no clear distinctions between *Foxg1^−/−^* and *Foxg1^+/−^* embryos in the overt cellular organization of non-*Foxg1* expressing cephalic ectoderm.

However, when we investigated the cellular organization and appearance in the normally *Foxg1*^+^ olfactory-associated SCE rostral to the eye at E9.25, we noted differences between *Foxg1^+/−^* and *Foxg1^−/−^* embryos. For instance, the *Foxg1^+/−^* SCE exhibited much rounder (less flattened) cells interspersed with numerous cellular extrusions typically indicative of apoptotic cell death (compare the yellow bordered insets in Figure 7F and G). Additionally, examination of scanning electron micrographs of the normally *Foxg1*-positve SCE associated with the hinge region between mxBA1 and mdBA1 showed that both the *Foxg1^+/−^* and the *Foxg1^−/−^* embryos exhibited rounded cells; notably, the SCE of the mutant embryos in this region also exhibited numerous apparently apoptotic cellular extrusions (compare the red bordered insets in Figure 7F and G).

Because scanning electron micrographs suggested distinct topographies of putatively apoptotic cellular extrusions within the *Foxg1*-positive SCE, we utilized whole-mount TUNEL assays of E9.25 *Foxg1* heterozygous and mutant embryos to confirm the topography of apoptotic cell death. Supportive of the scanning electron data, we found that the normally *Foxg1*-positive hinge-centered oral and caudal (i.e., aboral, peri-cleftal) mxBA1 SCE associated with the hinge presented an intense clustering of apoptotic cells in *Foxg1^−/−^* embryos whereas the *Foxg1^+/−^* embryos were, with the exception of a few cells associated with the first pharyngeal cleft, nearly apoptotic free in the same region (Figures 7H-K). The patent intense peri-cleftal mxBA1 apoptotic signal in mutant embryos, moreover, extended from the hinge region to the region of the presumptive trigeminal placode (which normally contains an elevated level of apoptotic cells). Despite its continuity with the mxBA1 apoptotic domain, however, *Foxg1^−/−^* embryos did not appear to display the same extent and rostral extension of apoptotic cells characteristic of the normal trigeminal. Dramatically increased and intensive apoptotic cellular clustering also characterized the mutant proximal second branchial arch (hyoid, or BA2). Additionally, the topography and extent of otic and epibranchial apoptotic cells differed between the E9.25 *Foxg1^+/−^* and the *Foxg1^−/−^* embryos.

TUNEL analysis further demonstrated a stark disparity between the presence of apoptotic cells in the olfactory and stomodeal SCE of E9.25 *Foxg1^+/−^* and the *Foxg1^−/−^* embryos, further confirming the scanning electron microscopic analysis (Figures 7H-K). Specifically, whereas the SCE of *Foxg1^+/−^* embryos was characterized by consistent, focalized concentrations of apoptotic cells in the olfactory-associated SCE (outlined in yellow, Fig 7J) and midline stomodeal ectoderm ventro-caudal to the cortical plate extending toward Rathke’s pouch (outlined in light blue, Fig 7J), these same regions in the *Foxg1^−/−^* embryos were notably absent of surface apoptotic cells. We noted furthermore that, while both *Foxg1* heterozygous and mutant embryos displayed a focal clustering of apoptotic cells overlying the cortical plate, the concentration of cells appeared slightly intensified in the mutant embryos and was, moreover, extended further ventro-caudally. Furthermore, while both *Foxg1^+/−^* and the *Foxg1^−/−^* embryos broadly exhibited apoptotic cells in the lens-associated SCE, there appeared to be slightly fewer in mutant embryos, and these were more topographically oriented with the SCE covering the middle of the telencephalon than with the olfactory SCE as in the heterozygous embryos.

The stark, unexpected alterations of the topographic profile of apoptotic cells in the SCE of E9.25 *Foxg1^−/−^* embryos led us to further profile cephalic apoptosis at E10.75 (Figure 7L, M). As with the earlier E9.25 embryos, we found a great disparity between the apoptotic profiles of *Foxg1* heterozygous and mutant embryos. As expected, cephalic cell death in *Foxg1* heterozygous embryos was characteristically wild-type in nature: For instance, the caudal mxBA1 proximal to the hinge exhibited a large swath of apoptotic cells running toward the developing trigeminal ganglion (itself characterized by distinct patches of apoptotic cells). A small discrete patch of dying cells likewise characterized the central region of the proximal BA2 but very few apoptotic cells were found associated with the first pharyngeal clefts. Moreover, a distinct supra-optic cluster of apoptotic cells overlay the lens pit, which was itself entirely rimmed by an intense, continuous clustering of apoptotic cells. Small numbers of apoptotic cells were also found to line the forming nasolacrimal groove as well as the dorsal rim of the olfactory pit of *Foxg1* heterozygous embryos.

In each of these in regions of *Foxg1^−/−^* embryos the apoptotic profile differed (Figure 7L, M). While the mxBA1 associated cluster of apoptotic cells typical of *Foxg1* heterozygous embryos was also evident in mutant embryos it ectopically expanded distally through the hinge region well into mdBA1. The proximal BA2 of mutant embryos presented both a cluster of apoptotic cells typical of heterozygous embryos but presented also a second, intensely stained cluster of apoptotic cells between the this first patch and the first pharyngeal cleft. Epibranchial and otic associated mutant ectoderm also displayed greater numbers of apoptotic cells. Notably, a swath of apoptotic cells in mutant embryos did not entirely rim the lens pit but were only found in the rostral and dorsal compartments, the latter of which formed the base of a continuous patch of apoptotic cells dorsally extending to a highly apoptotic supra-orbital region. Moreover, apoptotic cells ran caudally as a continuous sheet from the trigeminal region, over the lens, and through the groove between the telencephalic and frontonasal protrusions. The ectopic pattern of cell death encountered over the middle of the telencephalon at E9.25 was also clearly patent at E10.75. Rather than running toward the eye, cell death associated with nasolacrimal groove was re-oriented toward the ectopic mxBA1 bulge (which itself remained relatively free of cell death; yellow arrowheads, Figure 7M) in *Foxg1* mutants. Lastly, both the dorsal and medial rims of the olfactory pit exhibited greater levels of apoptotic cells in *Foxg1^−/−^* embryos. Thus, supporting the scanning electron microscopic analysis of the cellular organization and apoptotic profile appearance of the SCE, whole mount TUNEL analysis demonstrated patent topographic alterations of the patterns of SCE apoptotic cell death from E9.25 to E10.75, times essential to the establishment and functional execution of epithelial patterning centers for the craniofacial skeleton.

### Loss of *Foxg1* leads to the ectopic acquisition of deep mandibular (lower jaw) molecular identity in the maxillary (upper jaw) primordia

Extensive experimental evidence has demonstrated that two key features of gnathostome jaw development are, one, that the BA1 CNC giving rise to the skeletal structures of the jaws are (initially at least) fungible and, two, that the delineation of upper jaw from lower jaw identity requires the establishment of a specific nested pattern of *Dlx* genes within the BA1 CNC (6, 10, 228, 229). Specifically, expression solely of *Dlx1*/*2* within the BA1 CNC establishes maxillary, upper-jaw identity while focalized restriction, or nesting, of *Dlx5*/*6* and then *Dlx3*/*4* within mdBA1, in combination with *Dlx1*/*2*, establishes deep mandibular, lower-jaw identity. While the complete and precise mechanisms establishing the mxBA1-excluded, nested mdBA1- restricted expression of *Dlx3-Dlx6* in otherwise fungible CNC have remained unclear, they plausibly involve unfolding, discriminate epithelial signals emanating from the hinge and/or the λ-junction – both of which are formed of *Foxg1*^+^ SCE. This notion, together with 1) the extensive ontogenetic alteration in the topography of SCE apoptotic cell death, 2) the evident progressive changes of signaling (patterning) gene epithelial expression patterns patent in *Foxg1^−/−^* embryos, and 3) perturbation in the upper jaw and hinge-centric skeletal structures led us to hypothesize that, in the absence of *Foxg1*, BA1 molecular restriction (nesting) of *Dlx* genes would be compromised.

To address this hypothesis we assayed the nesting of expression at E10.5 of *Dlx2, Dlx5* and *Dlx3* in both *Foxg1^+/−^* and *Foxg1^−/−^* embryos. We found that expression of *Dlx2* was maintained in the CNC in both mxBA1 and mdBA1 in the absence of *Foxg1*, although it did differ in a number of other cephalic tissues. (Figures 8A-D). For instance, the normal, distinct epithelial expression of *Dlx2* associated with the λ-junction was completely lacking in *Foxg1^−/−^* embryos, as was expression associated with the basal telencephalon, while expression between the proximal BA2 and the otic vesicle was slightly expanded. Loss of expression associated with the basal telencephalon also characterized *Dlx5* expression in *Foxg1^−/−^* embryos. However, while usually strictly restricted to mdBA1 CNC, intense *Dlx5* expression surprisingly was also strongly detected in the mesenchyme of the ectopic bulge within mxBA1 (Figures 8E-L). Within mdBA1 and BA2, moreover, *Dlx5* expression remained intense (though it did not extend as much proximally). *Dlx5* epithelial expression associated with the λ-junction and olfactory pit was also disturbed by the loss of *Foxg1*, in particular by the lessening of expression intensity in the distal rim and medial pit associated with the VNO as well as by ectopic expansion through the λ-junction, under the optic primordia, and toward the ectopic mxBA1 bulge. Like *Dlx2* (though to a lesser extent), expression of *Dlx3* was aborgated within the λ-junction eithelium, while expression was also detected (like *Dlx5)* within the core of the ectopic mxBA1 bulge (Figures 8M, N).

**Figure 8:**
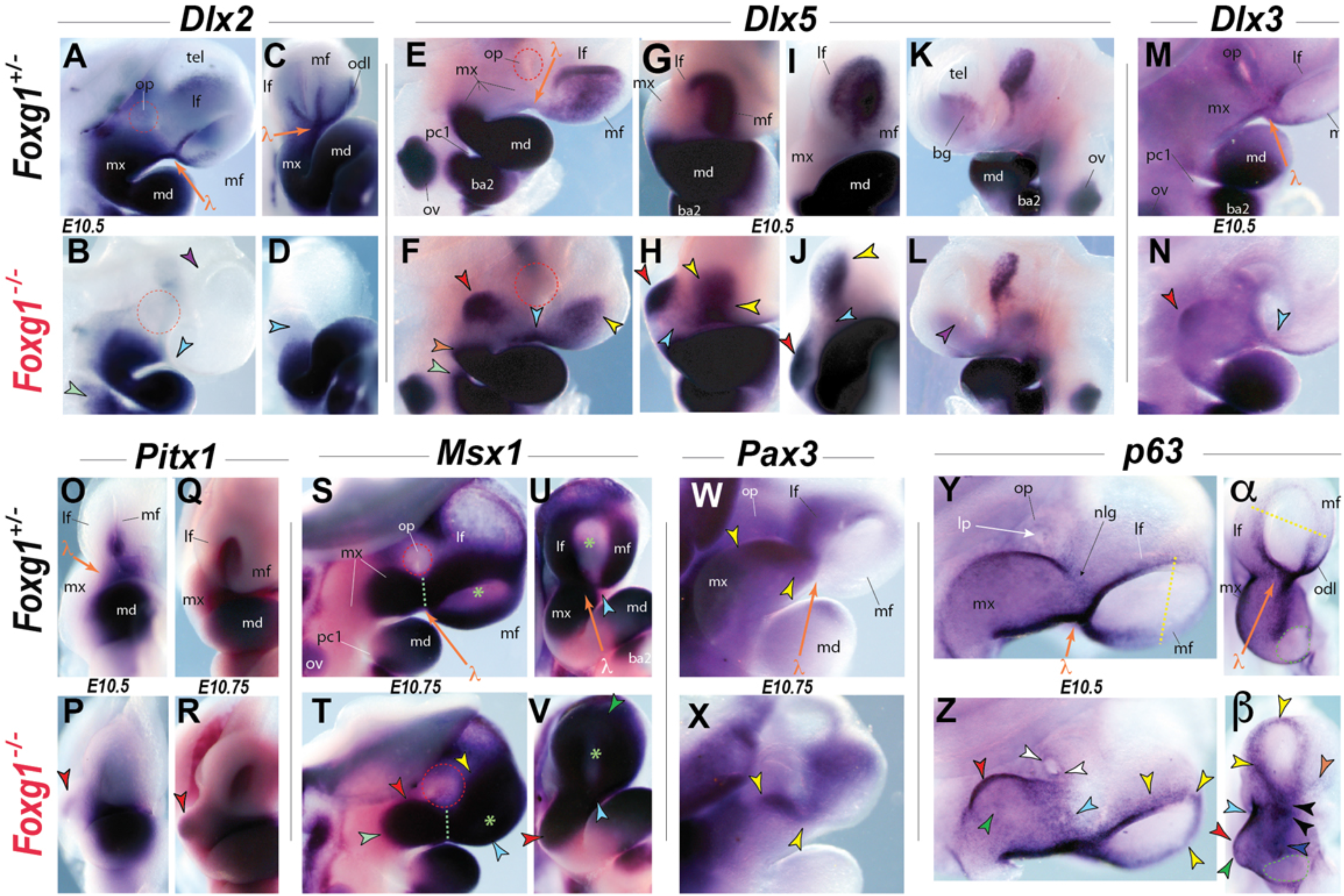
The acquisition of deep mandibular markers in the upper jaw primordia. (**A-β**) *In situ* hybridization of *Foxg1^+/−^* and *Foxg1^−/−^* littermates at E10.5 and E10.75. <u>(**A-D**</u>) Loss of epithelial *Dlx2* expression at the λ*-*junction. The ***dashed red line*** encircles the optic primordia. **<u>Arrowheads</u>**: ***green*** slight proximal expansion of expression in the BA2; ***light blue*** loss of expression in the epithelium of the **λ***-*junction; purple absence of expression in the basal telencephalon. <u>(**E-L**</u>) Expression of *Dlx5* in the ectopic bulge of mxBA1. **<u>Arrowheads</u>**: ***light blue*** ectopic streak of ectodermal expression running from the olfactory pit to the ectopic bulge; ***light green*** proximal extension of expression in BA1; ***orange*** proximal extension of expression in BA2; ***purple*** loss of expression in the basal telencephalon; ***red*** ectopic bulge of mxBA1; ***yellow*** altered expression in the olfactory ectoderm. The ***dashed red line*** encircles the optic primordia. <u>(**M, N**</u>) Expression of *Dlx3* in the ectopic bulge of mxBA1 (***red arrowhead***) and decreased expression at the **λ***-*junction (***light blue arrowhead***). <u>(**O-R**</u>) *Pitx1* expression in the ectopic bulge of mxBA1 at E10.5 and E10.75. <u>(**S-V**</u>) Alterations in *Msx1* at E10.75. **<u>Dashed lines</u>**: ***red*** encircles the optic primordia; ***green*** nasolacrimal line. ***Green asterick*** indicates the center of the olfactory pit. **<u>Arrowheads</u>**: ***green*** increased signal in the mesenchyme of the dorsal rim of the FNP; ***light blue*** increased expression in the ventral mFNP mesenchyme at the **λ***-*junction; ***light green*** expansion of the mesenchymal expression domain in mxBA1; ***red*** position of the ectopic bulge of mxBA1; ***yellow*** more intense staining in the mesenchyme overlying the telencephalic vesicle. <u>(**W, X**</u>) Relative restriction of *Pax3* in mxBA1 (***yellow arrowheads***). <u>(**Y-**</u>**β**) Altered topography of *p63* expression in the cephalic ectoderm. **<u>Dashed lines</u>**: ***green*** encircles the mesenchymal core of the resected hinge region; ***yellow*** normal demarcation line between higher ventral and lower dorsal expression rimming the olfactory pit. **<u>Arrowheads</u>**: ***black*** disorganized and patchy expression at the center of the λ-junction; ***dark blue*** discontinuous and disorganized expression in mxBA1 oral epithelium; ***green*** decrease in the level of *p63* signal associated with the ectoderm of the ectopic bulge in mxBA1; ***light blue*** broadened, caudally shifted patch of elevated *p63* expression associated with the nasolacrimal groove; ***orange*** abnormally defuse expression in the odontogenic line; ***red*** position of the ectopic bulge of mxBA1; ***white*** aberrant focalized expression along the lens pit; ***yellow*** increased *p63* expression abnormally broadly rimming both the dorsal and the ventral olfactory pit. **<u>Abbreviations</u>**: ***ba2***, second branchial arch; ***lf***, lateral frontonasal process; ***lp***, lens pit; ***md***, mandibular BA1; ***mf***, medial frontonasal process; ***mx***, maxillary BA1; ***nlg***, nasolacrimal groove; ***odl*** odontogenic line; ***op*** external optic primordia; ***ov***, otic vesicle; ***pc1***, first pharyngeal cleft; ***tel***, telencephalon; **λ**, center of the lambdoidal junction.

To confirm that the CNC of the ectopic mxBA1 bulge had acquired a mdBA1 molecular identity outside of de-regulated *Dlx* expression, we examined *Pitx1* expression at E10.5 and E10.75 in *Foxg1* heterozygous and null embryos. Normally, *Pitx1* is expressed within a distal subdomain of mdBA1 CNC and not in the mxBA1 CNC topographically akin to the ectopic bulge (Figures 8O, Q). However, in the absence of *Foxg1*, *Pitx1* expression was detectable in the mxBA1 ectopic bulge at both stages (Figures 8P, R). Moreover, *Pitx1* expression typically associated with the λ-junction and stomodeal ectoderm was absent in *Foxg1^−/−^* embryos.

To examine whether the *Dlx3/Dlx5/Pitx1*-positive ectopic bulge was delineated and entirely offset from the remainder of mxBA1, we examined mesenchymal *Msx1 expression*. Normally, *Msx1* is highly expressed in the both the upper and lower jaw ‘caps’ CNC; specifically with regard to mxBA1, it is highly expressed in the mesenchyme of the λ-junction, including the mxBA1 and FNP, wherein expression is weak dorsally and strong ventrally (except where the mFNP meets mxBA1), and not in most of the hinge region mesenchyme (Figures 8S, U). We found that, in *Foxg1^−/−^* embryos, *Msx1* expression in mxBA1 was both continuously extended from the λ-junction through to, and including, the ectopic bulge and was abnormally high throughout the lFNP and mFNP (Figures 8T, V). mxBA1 expression of *Pax3*, normally also restricted at the λ-junction to the mesenchyme of mxBA1 and the lFNP, however, did not follow the pattern of *Msx1* but rather was abnormally restricted closer to the λ-junction in *Foxg1^−/−^* mutants (Figures 8W, X).

The unexpectedly ‘mdBA1’ molecular nature of the ectopic mxBA1 bulge evinced in *Foxg1^−/−^* embryos, combined with the abnormal gene expression patterns and altered cellular dynamics patent in the λ-junction and the sub-ocular ectoderm between the λ-junction and the bulge, led us to further examine the molecular state of the cephalic ectoderm in this region. For this, we chose to examine the expression of the transcription factor p63 in part because *p63* mutations have been implicated in cephalic ectodermal dysplasias, and it is believed to control the relative state of proliferation and differentiation, as well as of apoptosis, in cranial ectoderm associated with the λ-junction (68, 230–235). Normally, discrete concentrations of *p63* positive cells are seen at the center of the λ-junction, including rimming both the lFNP and mFNP of the ventral olfactory pit, along the odontogenic line, along the center of the oral epithelium of mxBA1 and the nasolacrimal groove (Figures 8Y, α). Moreover, *p63*-positive cells are typically found spread out over the entire surface of the lateral mxBA1 ectoderm. Mutant embryos, however, displayed a different pattern: *p63* positive cells were detected abnormally rimming broadly both the dorsal and the ventral olfactory pit but only weakly in the odontogenic line (odl, Figure 8Z, β) while expression at the center of the λ-junction and the mxBA1 oral epithelium was discontinuous and disorganized. Furthermore, rather than discretely lining the nasolacrimal groove, mutant embryos showed a broad, caudally shifted patch of elevated *p63* expression; notably, this patch of elevated *p63* expression was discretely offset by a patent decrease in the level of *p63* associated with the ectoderm of the ectopic bulge in mxBA1. Moreover, we found an altered topographic profile in *p63* signal in the cells of the lens pit of *Foxg1^−/−^* embryos.

### *Foxg1* genetically interacts with *Dlx5* in cranial skeletal development and SCE organization

Together, our results have demonstrated that *Foxg1* regulates considerable aspects of non-NE craniofacial development. To approach a greater understanding of the broader context in which *Foxg1* regulates SCE development and craniofacial patterning, we began a systematic examination of potential genetic interactions of *Foxg1* and other regulators of λ-junction associated SCE and cranial skeletal development. Among these regulators is *Dlx5*, which, in addition to being expressed in the CNC of mdBA1, is extensively expressed in the early embryonic SCE before being restricted to the olfactory pit around E10.5. We present here direct evidence of a significant genetic interaction between *Foxg1* and *Dlx5* for SCE development and cranial skeletal patterning.

Elucidation of a genetic interaction between two genes involves a demonstration that the progressive loss of individual alleles of the two genes results in a distinct (not simply additive) phenotypic alteration, and examination of the phenotypic consequences of the progressive loss of *Dlx5* alleles in a *Foxg1*-null background clearly demonstrates that, with regard to cranial development, *Dlx5* and *Foxg1* genetically interact (Figure 9). The phenotypic alterations evinced in *Foxg1^−/−^*;*Dlx5^+/−^* and *Foxg1^−/−^*;*Dlx5^−/−^* embryos and perinates are extensive, and therefore our descriptions herein are not exhaustive and are limited to those aspects pertinent (1) to the demonstration of a genetic interaction and (2) to the elucidation of the nature of this interaction.

**Figure 9:**
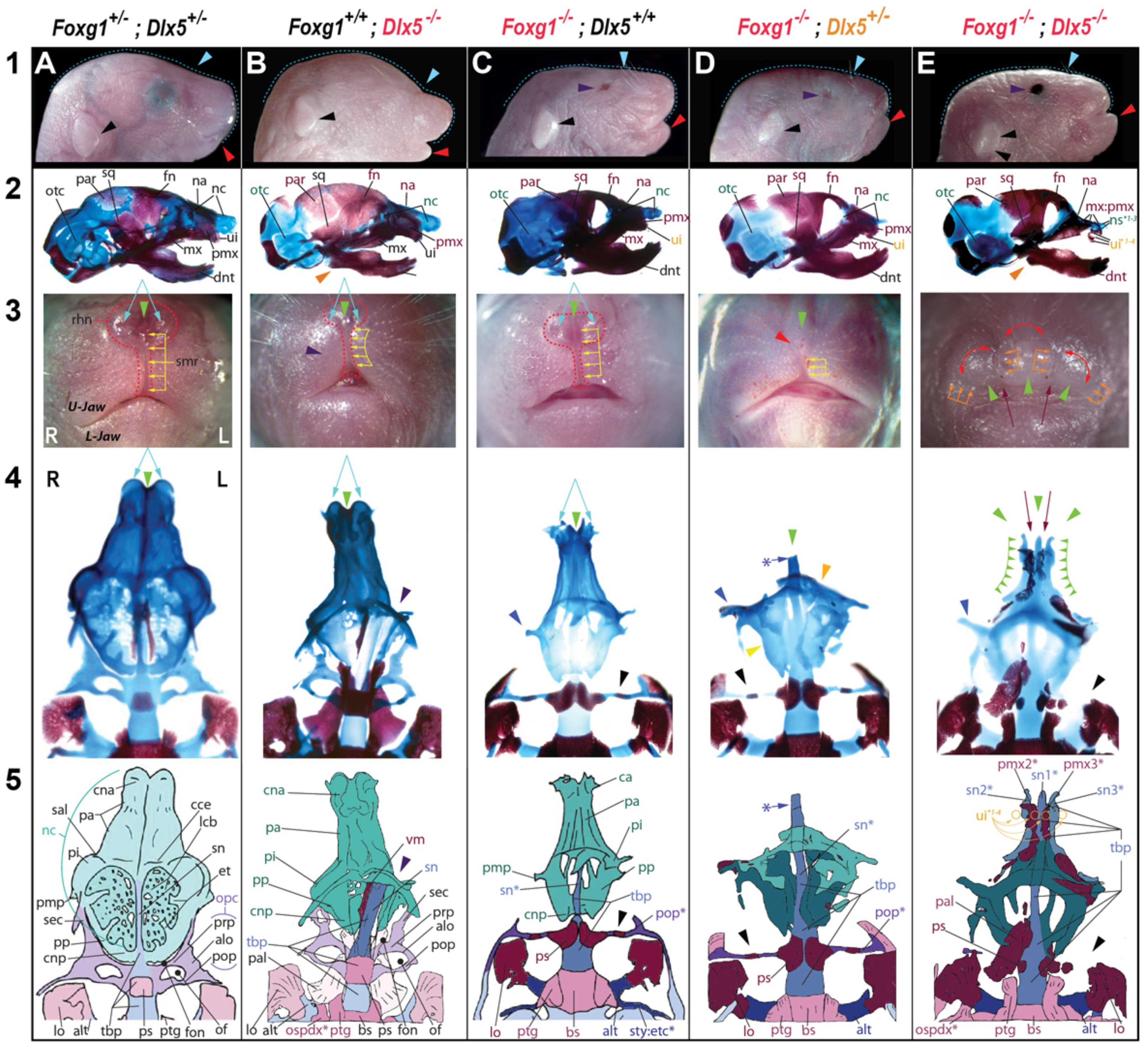
*Foxg1* genetically interacts with *Dlx5* and their mutual loss leads to anterior midline skeletal triplication, open eyelids, and complete absence of the rhinarium and external choanae. (**A1-E5**) Gross and cranial skeletal anatomy of compound *Foxg1; Dlx5* mutant neonates: *Foxg1^+/+^*;*Dlx5^+/+^* (**A**), *Foxg1^+/+^*;*Dlx5^−/−^* (**B**), *Foxg1^−/−^*;*Dlx5^+/+^* (**C**), *Foxg1^−/−^*;*Dlx5^+/−^* (**D**), and *Foxg1^−/−^*;*Dlx5^−/−^* (**E**). (**A1**-**E1**) Norma laterals view of the external anatomy. *Light blue dashed lines* represent the line of curvature (roughly from *inion* to *stomion superiu*s) of the head. *Arrowheads: black* apex of the auditory pinna; *red* relative position of the most distal point of the lower jaw (roughly *pogonion*); *light blue* caudal-most point of frontonasal vibrissae; *purple* open eyelids. (**A2**-**E2**) Norma laterals view of differentially stained skulls. *Abbreviations* in color indicate skeletal elements with altered morphology. *Orange arrowheads* indicate the unique proximal denture anatomy associated with the complete loss of *Dlx5*. (**A3**-**E3**) Norma frontalis view of the external anatomy. *Red dashed line* outlines the rhinarium. The *green arrowheads* indicate the nasal midlines. <u>Arrows:</u> *yellow* indicate the groove of the midline sulcus medianus of the rhinarium; *orange* indicate grooves that delineate the triplicate tissue masses associated with the ectopic midlines of the double null mutants; *maroon* interior grooves associated with the positions of the ectopic premaxillae and associated incisors; *double red* highlight the discrete nature of the triplicate midline masses; *double blue* positions of the external nares, which are absent in D3 and E3. (**A4**-**E4**) Differentially stained crania. The calvaria and most dermatocranial elements have been removed. For the sake of comparison, palatine bones have been preserved in the *Foxg1^+/+^*;*Dlx5^−/−^* B4) and *Foxg1^−/−^*;*Dlx5^−/−^* (**E4**) mutants. The attached green arrowheads highlight the course of the lateral, ectopic septum nasi. The *maroon arrows* indicate the ectopic dermatocranial ossifications, here taken as ectopic premaxillae, inserted between the ectopic septum nasi. The associated incisors (seen in **E2**) have been removed for clarity. The *blue asterisk and arrow* in **D4** indicates the position of where the anterior septum nasi of the sample has been removed. <u>Arrowheads:</u> *green* the anterior nasal midlines; *purple* asymmetric nasal capsular and trabecular basal plate development typical of the loss of *Dlx5*; *black* modification (**C4, D4**) or complete loss (**E4**) of the post-orbital pillar; *blue* modified posterior maxillary process; *orange* vertical ectopic cartilaginous plate; *yellow* horizontal ectopic cartilaginous plate. (**A5**-**E5**) Labelled schemae corresponding to **A4-E4**. **<u>Abbreviations</u>**: *alo*, ala orbitalis; *alt*, ala temporalis; *bs*, basisphenoid; *cce*, crista cribroethmoidalis; *cna*, cupula nasi anterior; *cnp*, cupula nasi posterior; *ect*, ectopic structure; *et*, ethmoturbinal; *ety*, ectotympanic; *fn*, frontal; *fon*, fissura orbitonasalis; *lcb*, lamina cribrosa; *L-jaw*, lower jaw; *lo*, lamina obturans; *mx*, maxillae; *na*, nasal; *nc*, nasal capsule; *of*, optic foramen; *opc*, optic capsule; *ospdx\**, ectopic os paradoxicum; *otc*, otic capsule; *pa*, paries nasi anterior; *pal*, palatine; *par*, parietal; *pi*, paries nasi intermediale; *pmp*, processus maxillae posterior; *pmx*, premaxillae; *pmx2*, ectopic* premaxillae 2; *pmx3*, ectopic* premaxillae 3; *pop*, post-optic pillar; *pp*, paries nasi posterior; *prp*, pre-optic pillar, opc; *ps*, presphenoid; *ptg*, pterygoid; *rhn*, rhinarium; *sal*, sulcus anteriorlateralis; *sec*, sphenethmoid commissure; *smr*, sulcus medianus, or phitrum, of the rhinarium; *sn*, septum nasi; *sq*, squamosal; *sty*, styloid process; *tbp, t*rabecular basal plate; *ui*, upper incisor; *ui1-4\**, mutant and ectopic upper incisors; *U-jaw*, upper jaw; *vm*, vomer.

We found that compound heterozygous *Foxg1^+/−^*;*Dlx5^+/−^* mice were viable and fertile, and when inter- breed they generated the expected ratios of wild type, single heterozygous, compound heterozygous, heterozygous plus homozygous, and both single and double homozygous genotypes, including *Foxg1^+/−^*;*Dlx5^−/−^*, *Foxg1^−/−^*;*Dlx5^+/−^*, and *Foxg1^−/−^*;*Dlx5^−/−^* mice (Figure 9). *Foxg1^−/−^*;*Dlx5^+/+^* perinates from *Foxg1^+/−^*;*Dlx5^+/−^* intercrosses exhibited the phenotypes described above (see also Fig 9C1-5), while *Dlx5^−/−^* perinates showed phenotypes as previously described (6, 79, 80, 227; see also Fig 9B1-5). For instance, the external anatomy of *Dlx5^−/−^* perinates exhibited hypoplastic nasal snouts, nares and rhinaria (with associated asymmetry of the sulcus medianus), as well as slight micrognathia, malformed auditory pinnae, and occasional exencephaly, while the cranial skeleton showed (as previously reported) otic capsular deficiencies, nasal capsular hypoplasia and (typically) asymmetry, deficiencies of the proximal dentary and its associated gonial and ectotympanic bones, and the presence of an ectopic ‘os paradoxicum’ involving Meckel’s cartilages and the pterygoid bones. While *Foxg1^+/−^*;*Dlx5^−/−^* perinates were distinct from both *Foxg1^+/−^*;*Dlx5^+/−^* and *Foxg1^+/+^*;*Dlx5^−/−^* perinates, for simplicity and brevity we focus below on the phenotypic consequences of the successive loss of *Dlx5* alleles in a *Foxg1* null background and their demonstration of a genetic interaction.

Compound *Foxg1^−/−^*;*Dlx5^+/−^* mutants were easily distinguishable from *Foxg1^−/−^* mutants (Figure 9); their snouts were noticeably shortened rostro-caudally and, while a reduced rhinal sulcus medianus was patent, the remainder of the rhinarium and external nares were not evident (Fig 9D3). While a rostroventral neurocranial midline was represented in compound mutants by a TBP it was more hypoplastic than that of the single *Foxg1^−/−^* mutants but was rostrally continuous with a cylindrical, rostro-caudally compressed rod-like nasal septum. Notably, nasal capsules and a true cavum nasi were absent (i.e., there were no pars anterior, pars intermediale, and pars posterior); rather, three dysmorphic cartilaginous structures were present associated with the rostral neurocranial midline. The first was a dorso-ventrally oriented, discontinuous plate (shaded lighter green in Figure 9D5) and the other two were bilateral laminar structures perpendicular to the first and likely representing the remnants of the paired lamina transversalis posterior (shaded darker green in Figure 9D5). These NC defects occurred concomitant with defects of the regional dermatocranial bones, including the premaxillae, maxillae, and palatines, and are associated with development at the λ-junction. Moreover, the TT of compound mutants was also found to be detached from the otic capsule but was synchondrotic with the re-oriented styloid process (stylohyal plus tympanohyal); together, this fused ‘stylo- tegmenal’ structure was widely separated from the incus (unlike in the *Foxg1^−/−^* mutants), which was itself further dysmorphic with its crus longus re-oriented toward the stylo-tegmen. The loss of a single *Dlx5* allele in a *Foxg1* null background, moreover, resulted in further hypoplasia of the pars canalicularis of the otic capsule. Importantly, each of these defects of the craniofacial skeleton and associated soft tissues are sufficient to demonstrate the genetic interaction of *Foxg1* and *Dlx5*.

### Loss of both *Foxg1* and *Dlx5* leads to fundamental re-organization of the anterior skull

In a number of fundamental respects the phenotype of *Foxg1^−/−^*;*Dlx5^−/−^* double mutants differed significantly from either the *Foxg1^−/−^* or the *Dlx5^−/−^* single mutants as well as the compound *Foxg1^−/−^*;*Dlx5^+/−^* (and *Foxg1^+/−^*;*Dlx5^−/−^*) mutants; the most significant of these is in the re-patterning of the anterior cranial skeleton, which in *Foxg1^−/−^*;*Dlx5^−/−^* double mutants is represented not by one midline extension but by what is effectively three midline extensions (Fig 9E1-5). Examination of the external anatomy of the frontonasal region of neonatal double mutants revealed in each a near complete absence of frontonasal vibrissae as well as the lack of a rhinarium, a midline sulcus medianus, and any external nares; in their place where three nearly identical raised tissue masses the middle one astride the anatomical midline (mid-sagittal plane) divided by sulci on each side (i.e., by four sulci in total) (Fig. 9E3). Remarkably, examination of the frontonasal skeleton reveled that each raised tissue mound was mirrored internally by one of three cartilaginous rods (Fig 9E4, 5). Each rod was bounded on either side by dermatocranial bone that was, in turn, associated with a dysmorphic, diminished incisor. These four rod-associated ossifications extended caudad to highly hypoplastic presumptive premaxillary bodies that were synostotic with likewise highly dysmorphic maxillae. The entire palatal space was found to be chaotic and cleft, not elevated, and characterized by numerous isolated palatine ossifications. Additionally, the elements of the middle ear and the otic capsule evinced further degradation of structures (data not shown). Notably absent, moreover, was the straight rod-like post-optic pillar seen in *Foxg1^−/−^* and the *Foxg1^−/−^*;*Dlx5^+/−^* mutants (Figure 9E5).

### Fundamental re-patterning of the SCE and LJ of Foxg1^−/−^;Dlx5^−/−^ embryo

Because the *Foxg1^−/−^*;*Dlx5^−/−^* double mutants showed such a drastic re-patterning of their anterior cranial skeleton, we posited that E10-E11 double mutant embryos would likewise evince fundamental architectural changes correlated with reorganized patterns of gene expression within the cephalic ectoderm. Indeed, when examined at E10.25, in addition to the hypoplastic telencephalon vesicles seen with the loss of *Foxg1*, the *Foxg1^−/−^*;*Dlx5^−/−^* double mutant embryos were found to be characterized by an absence of an olfactory pit (though there appeared to be some subjacent mesenchyme) and the truncation of the FNP- associated mxBA1 (Figure 10B, D). Notably, compound mutants did not exhibit the ectopic budge evident in the single *Foxg1^−/−^* mutants.

**Figure 10:**
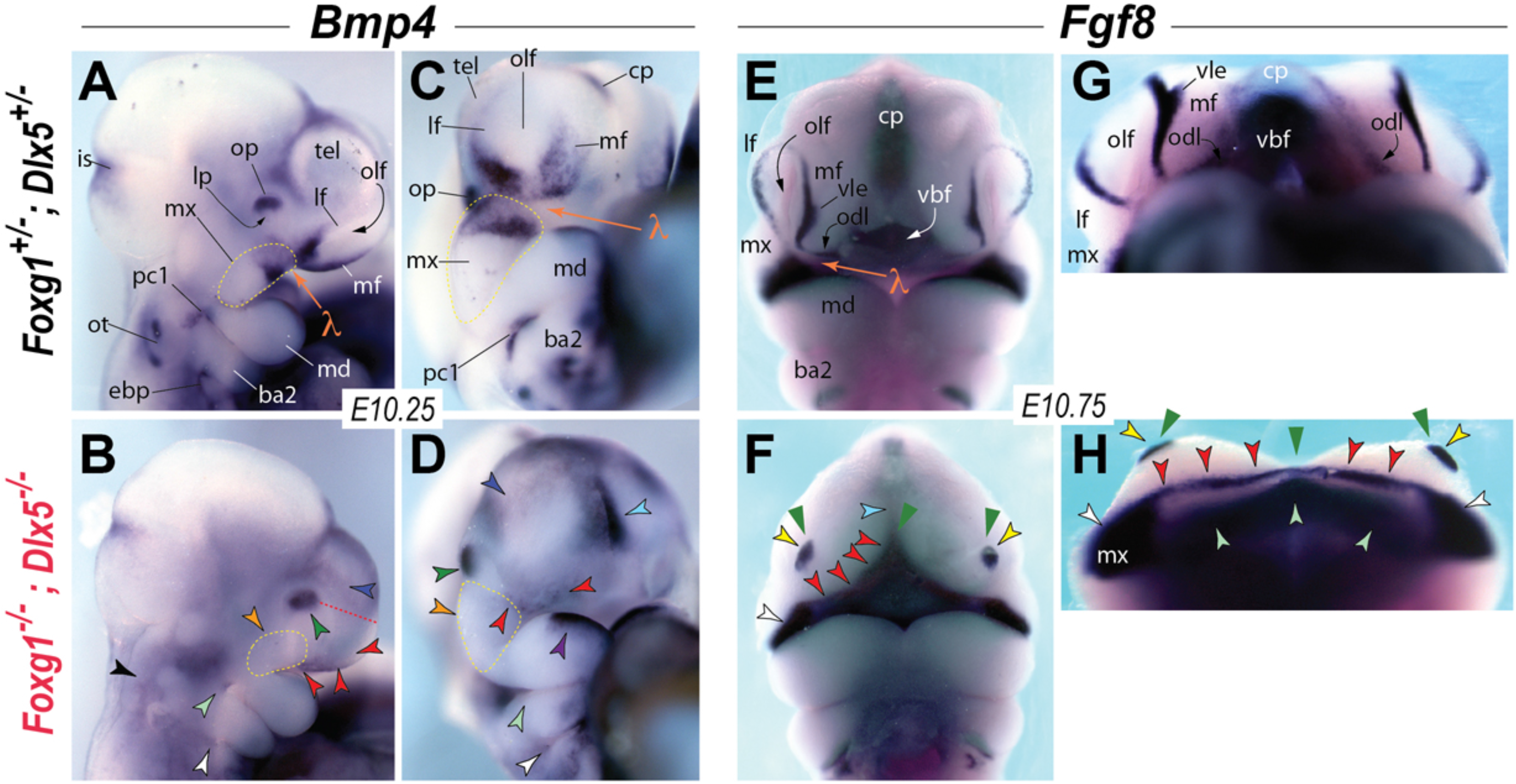
Fundamental re-patterning of the SCE and LJ of E10.5 Foxg1^−/−^;Dlx5^−/−^embryo presages the tripartite nasal midline structures. <u>(**A**-**D**</u>) Significant alteration of cephalic epithelial *Bmp4* expression at E10.25, including loss associated with the λ**-**junction. <u>Dashed lines</u>: *yellow*. outline of mxBA1; *red* distance from the lens placode (*op*) to the cephalic rostro-anterior extreme. **<u>Arrowheads</u>**: *black* loss of expression associated with the otic primordia; *dark blue* light supra-orbital and supra-olfactory expression; *green* absence of a lens pit and slight extension of expression; *light blue* increased signal at the *cp*; *light green* loss of expression at the *pc1*; *orange* absence of the ectopic bulge typically associated with the loss of *Foxg1*; *purple* maintenance of midline mdBA1 expression; *red* near complete loss of expression at the presumptive λ**-**junction, including in the *mx*, *lf*, *mf* and *odl*; *white* loss of expression associated with the *ebp*. <u>(**E**-**H**</u>) Alteration of cephalic epithelial *Fgf8* expression at E10.75 suggests re-patterning of positional information elaborated by the SCE to the subjacent CNC. <u>Arrowheads</u>: *light blue* decreased signal at the presumptive *cp*; *light green* expression associated with the *vbf*; *green* posited positional foci for the tripartite midlines; *red* strong, discrete band of presumptive odontogenic line (*odl)* ectodermal expression abnormally running medio-laterally to the meet the contralateral band; *yellow* small but focal patch of expression, topographically positioned where the olfactory pit (*olf)* should arise, centrally overlying a hillock of mesenchymal cells; *white* extensive signal in the oral ectoderm of mxBA1 (mx). **Abbreviations**: *ba2*, second branchial arch; *cp*, commissural plate; *ebp*, epibranchial placodes; *is*, isthmus; *lf*, lateral frontonasal process; *lp*, lens pit; *md*, mandibular BA1; *mf*, medial frontonasal process; *mx*, maxillary BA1; *odl*, odontogenic line; *olf*, olfactory pit; *op* external optic primordia; *ot*, otic primordia; *pc1*, first pharyngeal cleft; *stm*, stomodeum; *tel*, telencephalon; *vbf*, ventro-basal forebrain; *vle*, remnant of the ventrolateral ectodermal expression domain; λ, center of the lambdoidal junction.

Both of these aberrant phenotypes are plausibly correlated with molecular changes at the λ-junction. Therefore, as an initial read-out of the patterning road-map of the cephalic ectoderm and the λ-junction we examined the expression patterns of *Bmp4* at E10.25 and *Fgf8* at E10.75 (Figure 10). While we found, as expected, that normal distal mdBA1 *Bmp4* expression was robust in double mutant embryos, we also unexpectedly found a near complete loss of junctional *Bmp4* expression. Loss also characterized the first pharyngeal plate, the epibranchial placodes and the otic primordia of *Foxg1^−/−^*;*Dlx5^−/−^* double mutant embryos. The region of the antero-ventral CP, however, showed increased expression, as did (though to a lesser degree) the external optic primordia.

While the ectoderm of the double mutant was incapable of invaginating to form either a lens pit or an olfactory pit, the presence at E10.25 of frontonasal mesenchyme that subsequently yielded three structural midlines suggested that positional patterning was being imparted to the ectomesenchyme. As dynamic *Fgf8* expression is known to be essential for NC development and polarity as well as the establishment of the midline (158), we examined its expression at E10.75. We found that singular foci of epithelial *Fgf8* expression characterized the apex of conical frontonasal ectomesenchymal masses in the *Foxg1^−/−^*;*Dlx5^−/−^* double mutant embryos (Figure 10F, H). This expression was accompanied by a decreased level associated with the CP and a substantial increase in SCE expression running from the mxBA1 oral ectoderm, arcing dorsomedially to cross the midline, thereafter again bending ventrolaterally to the contralateral mxBA1 oral ectoderm. Thus both *Fgf8* and *Bmp4* expression profiles highlight the fundamental reorganization of the cephalic molecular pattering environment of the the *Foxg1^−/−^*;*Dlx5^−/−^* double mutant.

## Discussion

Identifying, recording, examining and subsequently outlining the etiology of organismal form, function and evolution remains a significant endeavor. Our own efforts in this regard have stemmed from the historic revelation that the development of the gnathostome skull is characterized by several universal traits (4–26, 135). Amongst these are 1) that regardless of the particulars of end-point phenotype, a basic seminal structural bauplan is initially ontogenetically followed and 2) that, in order to meet the distinct demands (e.g., trophic, reproductive, cognitive, locomotory, and so on) of a particular taxon, varied ontogenetic elaborations of this bauplan have manifested the great diversity in size, shape, and articulations of skeletodontal elements and overall forms observed. As fundamental, functional units within the skull, we have been specifically interested in the development and evolution of jaws — structures that notably integrate the organization of the splanchnocranium with that of the neurocranium. In discerning the comparative developmental paradigms of jaws we aim to examine and elucidate a fundamental example of the coordination of bauplan fidelity and design elaboration.

To address whether gnathostome jaws are all made in a like manner — and if not, then how not — a ‘Hinge and Caps’ model has previously been presented in which the articulation, and subsequently the polarity and modularity, of the upper and lower jaws is placed in the context of CNC competence to respond to positionally located epithelial signals (8–10, 30–32, 158). This model expands on an evolving understanding of the patent polarity within BA1 and explains a developmental patterning system that both keeps gnathostome jaws in functional registration (i.e., organizes the upper and lower jaw arcades to meet in appropriate functional alignment) and maintains developmental tractablity to potential changes in functional demands over evolutionary time. Further, it posits a system for the establishment of positional information where pattern and placement of the ‘Hinge’ is driven by factors common to the junction of mxBA1 and mdBA1 and of the ‘Caps’ by the signals emanating from the distal-most mdBA1 (i.e., forming the lower jaw Cap) and the λ-junction (the upper Cap). The functional registration of jaws is then achieved by the integration of Hinge and Caps signaling, with the Caps sharing at some critical level an earlier developmental history that potentiates their own coordination. Understanding of jaw development and evolution, then, follows from appreciation of the ontogeny of the positioning of these epithelial signaling centers and their influences on the subjacent CNC.

A salient sequent question becomes how the SCE associated with the λ-junction is ontogenetically organized and genetically regulated to participate in the development and evolution of the upper jaws and associated structures. In this regard, we have examined evidence of a developmental role played by a SCE- expressed master-regulatory transcription factor gene, *Foxg1.* We have hypothesized that, in addition to its known role in neurodevelopment, *Foxg1* participates in the ontogenetic elaboration of the λ-junction and subsequently in its regulation of jaw development. Herein, we have, for the first time, elucidated the significant breadth of influence that *Foxg1* exerts on non-neural cranial development. Specifically, our results, both anticipated and unexpected, demonstrate that *Foxg1* regulates: 1) cranial skeletogenesis, including the development of the neurocranium (e.g., the calvarium, each sensory capsule (OTC, OPC, and NC) and the TBP), the upper jaws, including the palate, and the middle ear; 2) the development of soft-tissue derivatives of the external cephalic ectoderm, including the auditory pinnae and meatal antra, as well as the upper lip, snout and sinus tubercle-associated vibrissae, the rhinarium and external nares, the auxiliary eye structures, and Rathke’s pouch, in addition to the NC olfactory and respiratory epithelia; 3) the formation of the internal choanae and elevation of the palatal shelves (both obligatory developmental events enabling mammals to simultaneously eat and breathe); 4) the elaboration of a properly organized and functioning embryonic λ-junction; 5) the organization and cellular dynamics of the SCE, including cell orientation, gene expression, and the ontogenetic topography of apoptosis; 6) the identity of upper-jaw elements by inhibiting, or restricting, the induction of deep mdBA1 molecular identity from within otherwise fungible mxBA1 CNC; and, with *Dlx5*, 7) the establishment of identity and patterning of the anterior cranial skeletal midline. Below, we further explore the significance of these results in the context of craniogenesis and evolution.

### Foxg1 is a significant craniogenic regulator of both the ‘contents’ and the ‘container’

Craniogenesis, the developmental process of elaborating a fully functional, integrated head, requires the coordination of developmental mechanisms regulating both the formation of the brain (i.e., the ‘contents’) and the skull (i.e., the ‘container’). Normal development of the contents and the container each require mechanisms that yield complex structures from precisely established developmental fields; and such developmental mechanisms for one must be, at some level, attuned to those of the other — a notion that has been clinically canonized in the aphorism that ‘the face predicts the brain’ (236). Indeed, coordinated cranial integration is reflected in the great numbers of both murine mutants and human patients suffering from both craniofacial malformations and coexisting brain malformations. Evidence presented herein, in combination with previous work on neurodevelopment, makes clear that *Foxg1* acts as a significant craniogenic regulator of both the contents and the container.

### *Foxg1* and the contents

Within the ‘container’, the mammalian ‘contents’ are spatially dominated by the telenecephalic cerebral hemispheres. As *Foxg1* regulation of their development has been extensively explored and discussed elsewhere (see **237 for a recent synthesis**), we only summarize here select observations beginning with some generalized views of telencephalon development.

Once the NE is induced from naive embryonic ectoderm, it becomes regionally partitioned rostrocaudally with the rostral-most domain fated to be the forebrain (prosencephalon) that will subsequently be subdivided into rostral telencephalic and caudal diencephalic domains (88, 110, 238–243). The telencephalic anlage is then further subdivided dorsally into the pallium, wherein most sensory afferent input is received from diencephalic (thalamic) nuclei, and ventrally into the subpallium, which includes the basal ganglia (BG), that largely process motor functions. Sequent patterning of the pallium generates a medial pallium forming the hippocampus and choroid plexus, a large dorsal pallium forming the neocortex (isocortex), and a lateral pallium (yielding, for instance, the olfactory cortex). These pallial subcomponents progressively acquire a complex layered organization in which precise neuronal circuitry underlies subsequent complex functional, perceptual, and cognitive behaviors. Neocortical layered organization and circuitry, forming the largest and most complex pallial unit in mammals, is developmentally elaborated during corticogenesis along two principle axes: 1) a tangential axis formed of migratory interneurons, streaming from the ganglionic eminences in the ventral telencephalon, that acts to partition neuronal populations into discrete functional domains that process aspects of sensation, movement and cognition; and 2) a radial axis of neurons progressively born *in situ* in the developing cortex through a controlled number of (at first) symmetric and then asymmetric cell divisions producing radial glial cells (RGC, or apical progenitor cells) and subsequently cortical neurons and/or intermediate progenitor cells (IPC) that radially migrate along RGC scaffolds to create, in an ‘inside-out’ manner, distinct cortical layers with unique patterns of connectivity.

*Foxg1* functions to regulate each of these neurodevelopmental steps, including forebrain induction, regional patterning and corticogenesis. Mice lacking functional *Foxg1* have significant cephalic hypotrophy with substantial morphogenic deficits of telencephalic structures, including dorsally of the neocortex and ventrally of the BG (Figure 2; Supplementary Figures 3, 4; 93, 113, 130). While notably already being expressed in the SCE anterior to the ANR at the three somite stage, subsequent expression of *Fgf8* in the ANR aids to induce *Foxg1* expression in the anterior, *Six3^+^* NE by the five somite stage. Once expressed in the rostral NE, *Foxg1* acts to establish a permissive telencephalic progenitor field (88, 237, 241).

After establishing this field, *Foxg1* then regulates the subsequent elaboration of the cerebral cortex in a number of ways. As embryonic growth proceeds, the neural tube is normally elaborated from the apposition of the left and right neural folds, which will have become molecularly regionalized antero-posteriorly, to establish several telencephalic organizing centers: 1) an anterior (rostral) organizer formed from the ANR and its ontogenetic derivatives (e.g. septal CP), which continues to express *Fgf8* as well as inhibitors of the Wnt/ß- catenin and Tgfß/Bmp signaling cascades that are being expressed by 2) a posterior (caudal) organizer formed in the caudomedial neural folds that will subsequently yield the roof plate and cortical hem (CH) in the dorsomedial ventricular walls of prosencephalon. Simultaneously, the ventral prosencephalic NE is axially organized by *Shh* expression from the notochord and prechordal plate (172–174, 177, 244). Integration of these centers of organization eventually yields a secondary organizing center, the ‘anti-hem’, at the pallial-subpallial boundary. *Foxg1* expression in the nascent telencephalic progenitor field has been shown to coordinate the cellular interpretation of these organizing centers (88, 89, 232). For instance, in the absence of functional *Foxg1* the ventral telencephalon of mice cannot properly interpret ventral signaling and the BG, derived from the ganglionic eminences and the source of tangentially migrating GABAergic cortical interneurons, fails to form (113, 237). Moreover, individual neural progenitor cells that lack *Foxg1* cannot contribute to the ventral telencephalic progenitors and GABAergic interneurons that arise from these progenitors (113).

Without *Foxg1*, moreover, Wnt- and Bmp-mediated signaling from the dorsomedial telencephalic anlage fails to be inhibited and subsequently pallial identity along the mediolateral axis is mis-specified with medial CH identity being extended laterad into tissue that would normally be forming the expansive, neocortex-yielding dorsal pallium; consequently, dorsal pallial neuroepithelial progenitor cells that should be filling out the neocortex as radial glial cells (RGC) instead act like cells of the CH and produce, en masse, Cajal-Retzius cells (CRC) (99–102, 115, 121, 123). CRC are regionally induced by the three dorsal, regional signaling centers (CP/pallial septum, the CH, and the anti-hem), are highly motile, and are thought to be the earliest post-mitotic cortical neurons to develop (246, 247). In mice, CRC normally exit the cell cycle beginning at E10.5 and migrate tangentially to evenly cover, through a combination of random walk migration and inter- CRC contact repulsion, the preplate/margial zone of the entire embryonic neocortex and, in a foundational manner, create an initial cortical layer 1. There, CRC interact at the neuroepithelial basement membrane with the nascent pial meninges and with the end feet of the pseudostratified neuroepithelia to regulate, principally through their expression of Reelin, the sequent layered, radial migration of pyramidal cells (248–257). This is a fundamental stage in corticogenesis but one that must be halted once completed. *Foxg1* acts to suppress CRC fate and thus expression is normally excluded from neuroepithelial cells in CRC producing regions (in a mechanism likely involving direct targeting by miR-9 (123)). *Foxg1* expression is strong, however, in the remainder of the necessarily mitotically active neuroepithelium where it acts to maintain the neuroprogenitor pool necessary to layer the cortex (110, 111). It is subsequently tied during cortigenesis to the regulation of the necessary temporal changes in competence to generate upper layer and deep layer neocortical neurons.

Notably, *Foxg1* expression is transiently downregulated as pyramidal neurons migrate through the neocortical intermediate zone and initially adopt a multipolar shape (115). Once the cells continue to their specific destinations in the cortical layers *Foxg1* is again up-regulated. Cells that fail to transiently inhibit *Foxg1* expression during this period stall their migration and subsequently fail to find their proper cortical layer. Pyramidal neuron precursors in their multipolar migratory phase and CRC share in common their predilection for decreased cell-cell connectivity, tangential migration, expression of Reelin, and repression of *Foxg1* expression. Moreover, telencephalic GABAergic interneuron precursors likewise selectively downregulate *Foxg1* during the tangential phase of their migration and reinitiate expression when they have invaded the cortical plate, suggesting that tangential migration in the forebrain universally requires transient repression of *Foxg1* (115). These observations also suggest that *Foxg1* regulates epithelial cell-cell interactions and cohesion as expression is lost when intra-epithelial cohesion is lost and subsequent migration ensues.

### *Foxg1* and the container

Gnathostome ‘containers’ are remarkable structures wherein a fidelity to an initial bauplan melds with a taxon specific elaboration of end-point design; a basic enterprise in studies of the skull is to understand and detail those patterning mechanisms at play in the manifestation of this initial structural bauplan as well as in the subsequent elaboration of skeletal form in disparate taxa, accounting for how taxonomic variability in morphology is achieved both ontogenetically and evolutionarily. Although some significant progress has been achieved in understanding the complex patterning mechanisms related to jaw development of BA-derived structures (10, 38, 44, 258), less has been achieved in understanding the intricacies of patterning the neurocranium or its integration with the FNP-associated upper jaw structures. We have been investigating the mechanisms by which the epithelium of the SCE associated with the λ-junction is organized and genetically regulated ontogenetically to participate in the development and evolution of the upper jaws and associated neurocranial structures. We have hypothesized that *Foxg1*, which is expressed in both the NE and the SCE associated with the λ-junction but not in the skeletogenic CNC (or cephalic mesoderm), acts to integrate neurocranial and viscerocranial development around the development of the upper jaws. Herein we have demonstrated that *Foxg1* is, in fact, a significant regulator of the craniogenesis of both the murine neurocranium and the viscerocranium, including of the upper jaws.

#### Foxg1 and the neurocranium

The rostral murine chondrocranial neurocranium is dominated by 1) the midline TBP that represents (at its caudal end) the skull base underlying the prosencephalic brain from the infundibulum and ventral hypothalamus to the tip of the olfactory bulbs and (at its rostral end) the intra-NC midline nasal septum, 2) the large NC, and 3) the OPC connecting them. Understanding the developmental mechanisms working to pattern and integrate these structures remains piecemeal despite their importance developmentally, structurally and evolutionarily (5, 8–15, 19, 20, 26, 54–63, 259–263). Though they unite dorsally, the NC are in essence paired, lateral cartilaginous cupules — themselves each composites of smaller, medially facing cupules — that unite with the TBP-derived midline structures to support the integration of the olfactory systems and the external nasal respiratory apparatuses. NC are inherently polarized along three axes and are patterned accordingly (5, 7, 11, 158). In addition to generating proper 3D polarity and proportionality, regulators of NC morphogenesis must accomplish numerous additional tasks, including: 1) positionally incorporate aperture access for both the external and internal choanae; 2) permit and establish access from the cavum nasi to the olfactory bulbs in the cavum cranii; 3) generate and sculpt the internal turbinate and paranasal sinus landscape to regulate directional airflow and to create a scaffold for the spatially ordered arrangement of olfactory and respiratory epithelia and associated glands; 4) integrate with midline TBP-derived structures; 5) connect with the OPC; 6) coordinate, along with the associated dematocranial elements (e.g., premaxillae, maxillae, nasals, lacrimals, vomers and frontals), skeletogenesis so that NC sub-structures properly undergo either specified (non-random) endochondral ossification and suturogenesis, remain cartilaginous, or are resorbed and disappear; and 7) elaborate ontogenically to meet the needs of the mature animal.

Detailed mechanisms for driving the attainment of these tasks mostly remain opaque though it is clear that the local SCE-derived epithelia are involved. A long history of experimental extirpation studies has demonstrated that the olfactory placode is essential to the formation and chondrogenesis of the NC (5, 168, 264–269). Moreover, lack of induction and specification of the olfactory placode within the SCE, such as is seen with the loss of *Pax6* (78, 82, 84), leads to the failure to form a NC (though the subjacent CNC do generate un-patterned cartilaginous plates). Once induced, olfactory placodal epithelia normally undergoes significant controlled morphogenic changes, in particular thickening followed by invagination, and medio- lateral, dorso-ventral and rostro-caudal partitioning that presages the subsequent polarity of the NC as well as the differential epithelial (e.g., respiratory or olfactory) domains within. Without such invagination there are no external chaoane and patterning of the cavum nasi and NC development is halted. Remarkably, loss of the olfactory placode does not necessitate the loss of an expansion of a frontonasal mass. For instance, *Pax6* mutant mice (by virtue of their lack of a placode) lack olfactory pits but do develop both a discrete frontonasal mass associated with a (albeit compromised) λ-junction in which CNC are patterned to form premaxillae as well as a (albeit truncated) TBP-derived nasal septum (78). This notion is likewise evident in the *Foxg1^−/−^*;*Dlx5^−/−^* mutants documented herein. Though compromised, rostral cranial skeletal patterning thus occurs in the absence of an olfactory placode and sources of regional patterning information must therefore exist outside of the placodal specific SCE and λ-junction. Experimental evidence, most significantly in avians, has identified a domain of the medial SCE, rostrodorsally formed of the *Fgf8^+^* epithelia overlying the CP and caudoventrally extending toward the *Shh^+^* epithelia medially overlying the stomodeum, as critical for the rostral outgrowth of the midline neurocranium and attendant parts of the upper jaws (158, 176, 270–274). This SCE sub-domain, referred to as the facial ectodermal zone (FEZ), appears to regulate the rostral extension of the TBP and its integration across the mid-line. Activity of the FEZ is ontogenetically controlled in part by an interplay between Bmp-, Shh- and Fgf8-signalling mediated by cells both in the developing subpallium and the intervening CNC. Though how, mechanistically, the patterning activities of the FEZ region and of the olfactory placode inter-relate and/or cross-regulate remains to be clarified, experimental evidence suggests that *Fgf8* in the SCE plays a clear role in both. Genetic attenuation of *Fgf8* signaling, for instance, results in re-patterning of NC polarity and dimensions and lack of integration across the midline of the NC-associated nasal septal TBP (158), and *Fgf8* has been implicated in repression of lens fate and induction of olfactory fate in the anterior SCE (222, 266). It is clear that *Foxg1*, expressed in the SCE yielding both the FEZ and the placode, partakes in regulation of the morphogenesis of the integration of the olfactory-associated neurocranium including both the midline TBP and the NC. Indeed, fulfillment of most of the NC-associated morphogenic tasks enumerated above is vitiated in the absence of *Foxg1*, including development of choanal access, elaboration of a skeletal cribriform plate delineating the cavum nasi from the cavum cranii, the formation of turbinates, and connection to the TBP and OPC.

With further regard to the TBP, careful embryonic cytoarchitectonic and correlative ontogenetic expressional topographic mapping and taxonomic comparisons of the adjacent sub-domains of the rostral prosencephalon have been conducted (e.g., 275–277) but it remains unclear exactly if or how prosencephalic sub-domains, as such, differentially partake in instructive cross-talk with the underlying CNC and SCE in influencing pattern of sub-domains along the rostro-caudal axis of the TBP. Moreover, as evinced by the loss of function of *Nkx2.1*, neurodevelopmental mis-specification within the basal prosencephalon does not automatically engender mis-patterning of the TBP (278; MJD, personal observations). It is notable, however, that loss of *Foxg1* results in a structural discontinuity between the presphenoidal portion of the TBP and the capsular portion as such discontinuities are typically not seen in other murine mutants where development of the TBP has been affected (e.g., *Pax6* or *Dlx5*). Together with the *Fgf8* (158) and FEZ (158, 176, 270–274) data, this discontinuity highlights the plausible notion that there is a plesiomorphic prosencephalic TBP (represented by that portion between the rostral end of the notochord at the basisphenoid and the rostral end of presphenoid and attachment of OPC) which is the pre-chordal, dura-adjacent cranial wall and the septum nasi extension of the TBP is an apomorphic evolutionary add-on. This suggests that there is profit in entertaining a pre-chordal TBP, primarily influenced by the developing sub-pallial/hypothalamic prosencephalon, integrated with a distinct capsular TBP, primarily influenced by the SCE, in particular the FEZ.

As with the TBP and NC, the OPC are greatly affected by the loss of *Foxg1.* Perhaps because of the lack of a discrete, front-and-center interorbital septum or of scleral cartilages and ossicles, or because of its smaller size relative to the NC and OTC, the OPC and its development are often over-looked in phenotypic analyses in mice. Investigations of the molecular mechanisms patterning the mammalian OPC are subsequently scarce (158, 279, 280). Herein we have shown that *Foxg1* is absolutely required for the normal development of the rostral OPC, including the pre-optic pillar, the ala orbitalis and the sphenethmoidal commissure: the *Foxg1^−/−^* OPC is represented only by an aberrantly straight, extended rod-like post-optic pillar with a precocious ossification center, an attenuated posterior commissure, and a precociously ossified ala hypochiasmatica synostotic with an abnormally split presphenoidal ossification center. Normally, the optic chiasm, wherein retinal ganglion cell axons from one side largely cross over to targets on the contralateral side, rests on the body of the presphenoid in the TBP. During *Foxg1^−/−^* craniogenesis, however, optic vesicles fail to resolve into normal optic cups or stalks. Rather, retinal tissue extends medially to the ventricle and the optic sulcus fails to close leaving an extensive ventral coloboma. The mutant retinal tissue does, however, generate an abnormal optic chiasm with a greatly increased number of ipsilateral projections (97, 105, 120). Thus, the disruption of midline patterning of the presphenoidal TBP appears mirrored by disruption in the overlying midline optic chiasm.

Moreover, the loss of OPC and related structures in *Foxg1^−/−^* skulls is particularly striking when viewed from norma lateralis (compare Figure 2E, F) where a complete absence of orbital skeletal structures permits visualization through the entire breadth of a cleared mutant neonatal skull, indicating that the cavum cranii here is utterly unprotected. This striking detail is not likely to be due simply to the loss of optic tissue. Indeed, orbital skeletal structures are present and patterned in the absence of retinal formation (e.g., *Rx^−/−^* skulls; 282); and, in fact, the absence of an entire eye, such as with *Pax6^Sey/Sey^* mutants, does not preclude the formation of OPC (or even an optic foramen; 78) or frontal bone orbital laminae. It is more likely that the lack of skeletal structure in *Foxg1^−/−^* mutants results from a knock-on effect due to the chaotic expansion of retinal tissue along with loss of appropriate skeletal patterning signals to the perioccular CNC.

The position and aberrant shape of the *Foxg1^−/−^* post-optic pilar is notable for a number of reasons. First, *Foxg1^−/−^* embryos exhibit an ectopic mxBA1 bulge at E10.5-E11 that is likewise post-optically positioned. Second, this bulge is filled with *Dlx5^+^* ectomesenchyme. And third, *Dlx5* is known to regulate both the straight, rod-like cylindrical shape and the cellular differentiation of MC in the lower jaw (6, 7, 79). Whether there is a direct correlation between the *Dlx5^+^* ectopic bulge and the aberrant *Foxg1^−/−^* post-optic pillar remains unclear, consideration must be given to the fate of these *Dlx5^+^* cells. Notably, *Foxg1^−/−^*;*Dlx5^−/−^* double mutant embryos lack — in addition to the regulatory effects of *Dlx5* — both an ectopic bulge and post-optic pillars of any kind.

Development in *Foxg1^−/−^* mutants of the vestibular and cochlear canals of the OTC has received some previous attention (104, 118). In addition to hypotrophy of both the pars canalicularis and pars cochlearis housing the canals (as previously described), we found that the oval and round windows of mutants appeared anomalous in size and orientation. The mammalian OTC is also normally perinatally synchondrotic with a number of extra-capsular structures, including the TT, the styloid process, and the alicochlear commissure; we found each of these structures to be dysmorphic and/or heterotopic in *Foxg1^−/−^* perinates. However, because each of these structures are also associated with the viscerocranium we discuss them in this context below.

Dermatocranial elements also make significant contributions to the calvarial neurocranium. These calvarial elements, including the frontal, parietal, interparietal, and squamosal bones, as well as the lamina obturans of the alisphenoid, are affected by the loss of *Foxg1*. While some of the evident calvarial deficits, such as those involving the parietal and interparietal bones, may reflect indirect structural changes due to hypotrophy of the *Foxg1^−/−^* brain, some are likely due to a direct loss of specific patterning information. For instance, *Foxg1^−/−^* perinates abnormally exhibited distinctly pyramidal shaped open interfrontal (metopic) sutures, much like those seen in the classic *short head* (*sho*) mutant (283). Sutural development here is highly dependent on cross-talk between the underlying meninges and the dorso-medially directed ossification fronts (284–290). For their part, the local cortical meninges also create a pial basement membrane that instructs both the local complement of CRC and the radial glia (247–256). It is tempting to posit that in *Foxg1^−/−^* mutants a local perturbation of CRC induction, numbers, and/or migration coupled with the loss of the normally underlying olfactory bulbs has had a counter affect on the local meninges and subsequently on patterning of the ossification front of the mutant frontal bones. Because of their association with the jaws, the squamosal and lamina obturans are further discussed below as are the FNP-derived premaxillae.

#### Foxg1 and the viscerocranium

Fundamental to gnathostome life is a set of functionally integrated jaws wherein the size, shape and articulations of each skeleto-odontal element are appropriately elaborated during ontogenetic development. Here we have demonstrated that *Foxg1* is an essential regulator of normal upper jaw development and morphogenesis. We have shown that, in the absence of *Foxg1*, the premaxillae, maxillae and palatine bones (the former two developing at the λ-junction and the latter two being core viscerocranial elements of the upper jaws) fail to properly develop palatal shelves that are elevated from the neurocranial base. Moreover, the pterygoids, which normally prop up the lateral boundaries of the internal choanae, are flattened and synostotic with the basisphenoids in the neurocranial base in mutant perinates. Therefore, mutants have no nasal respiratory passages, which results in choanal atresia. Thus, *Foxg1* regulates the development of structures deemed both indispensable to the colonization of land by tetrapods and to the mammalian ability to masticate while breathing, namely internal choanae and a secondary palate respectively.

In order to adapt to the new spatial demands of an expanding cerebral cortex, mammals evolved apomorphic modifications of their upper jaw elements at the expense of elements of the OPC neurocranial sidewall (5, 15, 19, 135, 291–293). In particular, the plesiomorphic antotic pillar (positioned posterior to the post-optic pillar but anterior to the OTC and connecting the OPC to the neurocranial base) was lost and was positionally replaced by a coopted segment of the splanchnocranial core of the upper jaw, the palatoquadrate cartilage (PQ). In the primitive condition, the PQ is typically integral to stabilizing the connection of the upper jaw to the neurocranium and is thus relatively intimate with it. Selective pressures resulted in the cooption and reorientation of a subset of mxBA1 CNC away from the rostral PQ core and toward the side wall (a process likely aided by the already intimate association of the palatal portion of the PQ with the neurocranial base). Through evolution the quadrate portion of the caudal PQ then became the mammalian incus. This process established the mammalian alisphenoid, comprised of the chondrocranial ala temporalis underlying the trigeminal ganglion and the dermatocranial lamina obturans which developed dorsal to the lateral tip of the ala temporals as a neomorphic dermatocranial structure that initiates endochondral ossification within the ala temporalis. Both of these components of the alisphenoid — the ala temporalis and the lamina obturans — are dysmorphic in *Foxg1^−/−^* skulls, suggesting the attractive hypothesis that *Foxg1* participated in mammalian craniogenic evolution by simultaneously aiding the lateral expansion of both the contents and the container.

While *Foxg1* regulation of inner ear development has been well investigated (104, 116) its role in middle (or outer) ear development has not. We have demonstrated here that the development of all three splanchnocranial middle ear ossicles is vitiated in the absence of *Foxg1*. In particular, the malleus and incus are aberrantly synchondrotic, and thus lack the mobile articulation necessary to properly amplify and transmit sound. Moreover, the mutant incus is further fused with an ectopic cartilage that, running rostrally towards the squamosal (itself drastically mis-patterned), overlies the incal-malleal articulation (‘in:tgt:ma:ect’, Figure 2). We take this ectopic structure to be the displaced (heterotopic) remnant of the TT. The TT, thought to be another mammalian neomorphic structure, typically rises from the OTC as a protective cover, or roof, in the tympanic cavity for the incal-malleal articulation. The TT is little studied developmentally, and understanding of the molecular mechanistic control of its development is limited, though disruption of *Dlx1* and *Dlx2* expression in the BA leads to its dissociation from the OTC and its association with ectopic PQ structures (80). The combination of 1) the location of its typical position, 2) its correlation with *Dlx* gene expression in the BA, and 3) its dysmorphic association with PQ-related structures in mutant mice (including that seen herein) altogether supports the hypothesis, counter to Allin and Hopson (1992), that the mammalian TT is actually a coopted mxBA1-PQ derivative rather than a neomorphic projection toward the quadrate originating from the otic skeleton (a discrepancy that focused, *Cre*-mediated CNC fate mapping in mice might resolve). Additionally, BA2 splanchnocranial derivatives are highly affected by the loss of *Foxg1*. For instance, the stapes lacks a foramen and the styloid process fails to synchondroticly attach to the OTC but instead shifts orientation and joins the aberrant alicochlear commissure. Notably, it is the proximal BA1 and BA2 derivative structures that are highly affected by the loss Foxg1.

When considered as a whole, we show that *Foxg1* plays a role in the development of structures involved in another remarkable mammalian innovation: the re-allocation of the functional jaw joint. In the pleisiomorphic gnathostome condition, the primary jaw joint is formed by the mobile articulation of two splanchnocranial elements, the quadrate in the upper jaw with the articular in the lower. In the apomorphic mammalian condition, this primary joint has been shifted to functioning in the middle ear wherein the quadrate has become the incus and the articular the malleus. A functional secondary jaw joint is then formed in mammals by the mobile articulation of two dermatocranial elements — the squamosal in the upper jaw and the dentary in the lower — that are ontogenetically and topographically closely associated with the quadrate/ incus and articular/malleus. As we have shown, the loss of *Foxg1* results in the mis-patterning of both the primary upper jaw joint element (the incus) and the secondary upper jaw joint element (the squamosal).

### *Foxg1* expression, the SCE, and epithelial morphogenesis

We initiated our investigation, in part, because *Foxg1* is expressed in what we posited were the right places and at the right times to play a key role in integrating development of both the content and the container during craniogenesis. Therefore, we examined embryonic *Foxg1* expression in the nascent and maturing SCE in more detail. Our β-galactosidase activity and confirmatory *in situ* hybridization analysis, taken together with that of previous investigations (e.g., 103, 130, 241), highlights several perhaps under- appreciated points regarding *Foxg1* expression and cranial development.

First, in addition to being expressed in the NE, *Foxg1* is initially expressed in the SCE that developmentally lies topographically anterior to the ANR. Specifically, it is expressed in the anterior cephalic ectoderm rostral and lateral to the neural plate beginning at the early 1-3 somite stage before subsequently being expressed in the ANR. It is not until later, at the 5-8 somite stage, that expression is then highly detectable in the NE of the anterior neural plate. Thus, expression in the anterior SCE precedes that in the neural plate. The fact that it is eventually expressed in the epithelium on both sides of the organizing influence of the ANR confirms that *Foxg1* is ideally expressed to act as an integrater of the patterning and functionality of both the contents and the container. While all of the plausible functional consequences of *Foxg1* being expressed first in the SCE are not fully understood, experimental evidence in chick and mouse has shown that midline ablation of this early *Foxg1^+^* SCE, without disruption of the nascent ANR, leads to the disruption of craniogenesis (294).

Second, *Foxg1* is more widely expressed than is perhaps commonly appreciated; indeed, we visualized fully penetrant β-galactosidase activity in *Foxg1^+/−^* embryos in cells of numerous locations previously unreported or otherwise uncharacterized, including along the apical ectodermal ridge of the limb buds and in cells representing the rostral midline of the roof plate of the mesencephalon and diencephalon beginning at E10.5. The positioning of these midline cells is notable in light of the calvarial defects encountered in *Foxg1* null perinates. Whether this expression, or any other newly noted above, represents directed *Foxg1* functionality remains unexplored.

Third, *Foxg1* is distinctly and dynamically expressed in SCE epithelia specifically implicated in the regulation of upper jaw pattern and development, and the ubiquity of early (i.e., through E9.5) *Foxg1* expression in the SCE indicates that most — if not all — epithelial cells associated with the epithelial component of the λ-junction have expressed *Foxg1* at some point. The developmental legacy for future craniogenic functionality of SCE cells having at one time expressed *Foxg1* remains to be fully clarified but is plausibly significant, particularly in light of the number of SCE ectodermal derivatives that are mis-patterned and/or malformed in the absence of *Foxg1*. Examination of the perioccular SCE, for instance, clearly reveals that the normally complex development of its derivatives has been disrupted by the loss *Foxg1*. Among other tasks, the periocular SCE normally must display positional competence to receive appropriate inductive signals, such as with induction of the lens, while also being unresponsive to inappropriate signals, and then participate in reciprocal cross-talk with both the developing optic cup and vesicle as well as the CNC in the POM. It must also be capable of repeated rounds of coordinated epithelial morphogenesis and migration to first generate a lens, invaginate to form a lens vesicle, separate as a vesicle from the SCE, form a distinct corneal layer, generate and properly position corneal (conjunctival) furrows, undergo focal glandular branching morphogenesis and tubulogenesis to form a nasolacrimal duct. It must also establish a migratory wave of epithelial periderm to generate eyelids that must first meet over the eye, fuse, and subsequently split open — all without becoming attached to the corneal surface (295–302). The periocular SCE in *Foxg1^−/−^* mutants fails in a number of these tasks. For example, the presumptive corneal SCE fails to suppress aberrant inductive signals resulting in ectopic sinus tubercles on its surface. Moreover, mutant lenses are truncated and misplaced (105) which stems, in part, from abnormal regulation of lens morphogenesis. Apoptosis is required for normal lens morphogenesis as it is required to establish the correct number of cells in the invaginating lens pit (303, 304), and apoptotic cells are clearly seen encircling the lens pit in E10.5 *Foxg1^+/−^* embryos; the forming lens pits in *Foxg1^−/−^* embryos, however, only have dorsally positioned apoptotic cells in the pits and lack ventral apoptotic cells.

The EOB phenotype in *Foxg1^−/−^* mutants provides a further instructive example. Mammalian eyelids are derived from the SCE and POM, and their formation involves a four step process: eyelid specification, migratory growth, closure, and re-opening (300, 305–312). As with the other SCE-derived auxiliary eye structures, such as the cornea and conjunctiva, the eyelids require — unlike the lens — neither an optic vesicle nor an optic sulcus to be specified and induced (282). Importantly, cells of the SCE must appropriately interact with the POM to ensure proper eyelid development. The molecular cross-talk between the SCE and subjacent POM is complex and involves numerous, diverse signaling cascades and transcription factors (300). Amongst the most crucial factors involved is RA engendered by *Raldh3* expression in the SCE and ventral optic cup, and *Pitx2* expression in the POM (200, 310, 311, 313, 314). Briefly, anterior ocular development depends on RA induction of *Pitx2* expression in the POM where it induces *Dkk2* to locally inhibit canonical Wnt signaling (311). Notably, there is evidence that *Pitx2* in the POM influences optic cup and stalk morphogenesis (200). Here we have shown that in the absence of *Foxg1*, *Raldh3* expression in the ventral ocular SCE is decreased, as is *Pitx2* in the ventral POM, plausibly explaining, in part, the significant anterior ocular defects, including EOB, phenotypes of *Foxg1* mutants.

Fourth, previously underreported *Foxg1* transcription is highly detectable at E10-E11.5 in a wide and intriguing pattern in the SCE covering the oral surface of mxBA1 and the stomodeal roof rostral to Rathke’s pouch (which is likewise *Foxg1^+^* and which underlies the forming TBP). The stomodeal midline cells here, conceivably contribute to the patterning activity purported to the FEZ. Stomodeal SCE expression ontogenetically persists even when other SCE expression is being restricted to only focally discrete epithelia such as the olfactory pits and invaginating choanae. This pattern of expression was accentuated when examining X-gal staining in *Foxg1^−/−^* embryos wherein the non-oral SCE stains abnormally high. Closer examination of this expression led to the realization that E10.5-11.5 *Foxg1^−/−^* embryos display a noticeably thicker stomodeal epithelium accompanied by an increase in the breadth of both Rathke’s pouch (the epithelium of which was actually thinner at this time) and the stomodeal roof itself — increases coincident with a positional reorientation of the FNP, mxBA1, and olfactory pits. SEM micrographs revealed further morphological changes in the *Foxg^+^* SCE of the stomodeal roof, mxBA1 and FNP of E11.5 mutant embryos, in particular the absence of invaginating choanae, the reorientation of stomodeal ridges, and a significant change in the coordinated and uniform directionality, size, and orientation of the surface epithelial cells (a trend continued through E12.5). Thus, with the loss of *Foxg1*, fundamental characteristics of the epithelial organization of the SCE are vitiated and the normal transcriptional distinction of oral verses non-oral ectodermal cells is lost.

And fifth, these changes in the characteristics of the oral ectoderm highlight the fact that *Foxg1* expression typically correlates with cranial epithelia that will, as a collective, undergo significant morphogenic changes. This is particularly true for the invagination of the adenohypophyseal and primary sensory placodes, the forming choanae, the epithelia lining the maturing NC, turbinates and the external nares, as well as for the sculpting of the epithelia of the primary and secondary palates, the upper lip, vibrissae, the auditory pinnae, the nasolacrimal duct, the rims of the conjunctival furrows from which the eyelids will develop, the pharyngeal plates, and (not least of all) the telencephalic neural epithelia (including the optic cup and sulcus). To ensure tissue integrity during epithelial morphogenesis, changes in cell motility and/or shape of the individual cells must be tightly coordinated with cell-cell adhesion, and *Foxg1* has been implicated in the coordination of neuroepithelial morphogenesis of the optic cup and patterning of the cup by regulating cell-cell cohesion within the retinal epithelia (119). Regulated cell-cell interactions are hallmarks of both the mechanics of epithelial morphogenesis and the planar polarization of cells in sheets (315–321). We posit that, where expressed, *Foxg1* universally acts to regulate cell-cell cohesion, integration, and orientation and thus to control epithelial morphogenesis and the dissemination of patterning information, a hypothesis that we are further investigating.

### *Foxg1*, the λ-junction, and upper jaw identity

Though there is great variability in size and functionality, all extant or extinct gnathostomes possess articulated jaw elements (4–27). From a mechanistic point of view, the principle developmental task in creating functionally integrated upper and lower jaws is to differentially and faciently instruct an otherwise fungible population of CNC to generate appropriately aligned skeleto-odontal structures. The ‘Hinge and Caps’ model was formulated, in part, to address this task. This model suggests that the first crucial step in achieving alignment is in establishing an epithelial ‘Hinge’ patterning source centered along BA1 that the CNC can differentially respond to based on their relative distance form the hinge source; thus fungible CNC within BA1 equidistant either proximally (i.e., within mxBA1) or distally (i.e., within mdBA1) from the source are predicted to respond similarly but in mirror image fashion. A testable readout of such a mechanism comes from the prediction that symmetrical patterns of responsive gene expression, centered around the hinge, should be evident and taxonomically conserved. We have previously tested this prediction and have shown the conservation of the predicted hinge-centered patterns of gene expression in shark embryos, as a representative basal gnathostomes, and amniotes (322).

The model also ensures developmental tractability in responding to evolutionary changes in functional demands on jaws by positing that the ‘hinge’ signaling is integrated with ‘caps’ patterning signals emanating from the distal mdBA1 ectoderm and the SCE at the proximal, mxBA1 end of BA1 (i.e., the λ-junction), two epithelia that share a very early developmental history (8, 9). Thus, the baseline of patterning jaws would have fungible CNC interpreting in mirror image fashion Hinge and Caps epithelial signals, and coordinated caps-related patterns of gene expression would be predicted by the model. Such patterns are indeed conserved in sharks, chicks and mice (322). Developmental modularity and integration are key mechanisms bridging development and evolution, and inherent within the patterning mechanism suggested by the Hinge and Caps model is the potential for developmental modularity within jaw units, a notion likewise supported by experimental evidence (30, 31). Ultimately, however, the model makes clear that understanding jaw development follows from further understanding the origins of, and influences on, the epithelial patterning centers. Notably, the model considers two additional factors. First, the upper jaws develop in more intimate association with the developing brain anlage, and thus upper jaw CNC has to contend with the interpretation of secreted signals otherwise being utilized to pattern the brain, while the lower jaw CNC has to contend with signals regulating the embryonic, pharyngeal vascular entry to the heart. Second, gnathostomes outside of chondrichthyans evolved a premaxillary component, derived from the FNP, in their upper jaw arcades and hence evolved a patterning organization to suit. Thus, the model predicts that there will be, on top of a common, mirror-image baseline of gene expression associated with BA1, factors independently influencing upper and lower jaw primordia. As with the ‘Hinge’-centered and ‘Caps’ coordinated patterns of gene expression noted above, the predicted presence of gene expression pattens unique only to upper jaw (e.g., *Raldh3*) or to lower jaw primordia (e.g., *dHAND*) is encountered and conserved in gnathostome embryos from sharks to amniotes (322). Importantly, the model posits that the λ-junction evolved as the coordinating ‘Caps’ patterning center in organisms incorporating premaxillary components in their upper jaws.

We and others have begun examining the developmental origins, influences, and functions of the λ-junctional epithelia (7–9, 68, 158). Here, we add *Foxg1* to the list of factors integral to upper jaw development and the establishment of a functional λ-junction. In the absence of *Foxg1*, the structural organization, orientation and size of λ-junction are altered, and the normal ontogenic topography of encoding signaling molecules (e.g., *Bmp4*, *Fgf8*) and transcription factors (e.g., *Dlx2/3/5, Sox10, Pitx1/2, Eya2, Six1/3, Msx1, Pax3, p63*) crucial to regional development is vitiated. An extensive body of evidence has demonstrated that the epithelia of the mature λ-junction (i.e., around E10.5 in mice) displays complex, refined patterns of morphogenic signals, including (among others) of Fgfs, Bmps/Tgfβs, Wnts, Shh, and RA synthesizing proteins, which regulate the morphogenesis of the craniofacial skeleton and its associated soft tissues (7–9, 68, 78, 107, 132, 158, 160–162, 295, 323–337). The specific question of the proximate instructive interactions leading to these refined, complex patterns within the mature λ-junction has unfortunately only infrequently been explicitly addressed outside of the context of 1) the possible role played by the nascent telencephalon and CNC or 2) the processes informing the individualization of cephalic placodes from the pre-placodal cephalic ectoderm.

With regard to this latter, from the presumptive cephalic and cervical ectoderm arises a pre-placodal region (PPR), unique to this surface ectoderm, that has been shown to represent a specialized competence to placodal induction (222, 225, 226, 266, 338–340). Experimental evidence supports the notion that once the PPR has been specified there follow sequent inductive steps to regionalize identity as posterior (otic and epibranchial) and anterior (olfactory, lens and adenohypophyseal), with the trigeminal being intermediate. It is within the anterior PPR that the SCE will give rise to the λ-junction. Notably, within this anterior domain all placodal progenitor cells are initially specified as lens regardless of subsequent fate, and induction of non-lens fate requires both the active induction of unique identity (through distinct sets of inductive cues) and the active repression of lens fate (which is mediated in part by the CNC) (222, 266). Induction of the olfactory placode appears to involve ANR-related Fgf expression to both help suppress lens identity and induce olfactory and trigeminal identity. Moreover, while short-term exposure to Bmp signaling promotes olfactory identity, long-term exposure promotes lens identity (225). We have shown that both *Bmp4* and *Fgf8* cephalic expression patterns are perturbed in *Foxg1^−/−^* embryos prior to maturation of the λ-junction. At E9.25, *Bmp4* is expanded in both the nascent lens placode and the presumptive λ-junction SCE that separates the lens from mxBA1 and the nascent olfactory placode, and, while expressed in the ventrally displaced ANR, *Fgf8* expression is also diminished in the SCE placed between the ANR and the forming olfactory placode. Disrupted *Bmp4* and *Fgf8* expressional topography subsequently also characterizes the maturate λ-junction of E10.5 mutant embryos.

Placodal progenitor cells within the PPR are molecularly characterized by their expression of the transcription factors *Six1/4* and *Eya1/2* (222, 225, 226). Examination of *Six1* null embryos reveals defects in placodal morphogenesis and neurogenesis — rather than in PPR formation — and a lack of both lacrimal glands and NC turbinates (334–33). Moreover, *Six1/4* compound mutant embryos have no invaginating olfactory placodes and extensive rostral neurocranial defects (341). Although *Six1/4* activate transcription in the PPR, their activity is topographically limited to the anterior PPR in chick, Xenopus and mouse (342). Investigation of the *Six1* enhancer region has found, moreover, that it is bound by Dlx5 protein and that it also has a *Fox* transcription factor binding domain which helps drive *Six1* expression in the SCE (343). Here we have shown that the topography of expression in the SCE of both *Six1* and *Dlx5* in E9.25 *Foxg1^−/−^* embryos is altered, in particular in the dorso-ventral condensing of the olfactory placode field and its aberrant extension under the lens and toward the mxBA1. Together with the vitiated *Bmp4* and *Fgf8* expression patterns, changes in *Six1* and *Dlx5* in the mutant SCE suggests that, while not required for placode induction, *Foxg1* is required for the normal topographic circumscription and compartmentalization of the placodal domains in the SCE. We posit that such circumscription of SCE domains is normally required for the appropriate elaboration of the λ-junction to effectively function as an essential integrating center of complex olfactory, optic, and upper jaw patterning.

The changes in the apoptotic profile of the SCE are particularly relevant in this light. As a normal, regulated cellular process, programmed cell death through apoptosis is well documented and clearly plays a role in organogenesis and tissue remodeling such as occurs during epithelial invagination (344–347). Apoptosis is also key to regulating cell number and the removal of transient tissues when and where no longer needed. Within the SCE apoptosis has been observed, for instance, separating the posterior placodes (epibranchial, otic, and trigeminal) (348, 349); it has furthermore been detected in the sculpting phases of olfactory pit and cavity elaboration (350) and in the postnatal pruning of olfactory receptor neuron numbers (351). Apoptosis is, moreover, a critical factor in the control of SCE-related ectodermal fusion at points of epithelial-epithelial contact, including closing of the neural tube, the lens and otic vesicles, and the secondary palate, as well as being required in the normal propagation mid facial epithelial seams between the mxBA1, lFNP and mFNP at the λ-junction, a process the dysfunction of which underlies clefting of the lip (68, 303, 342, 352).

We hypothesize that *Foxg1* controls upper jaw development and morphogenesis in part through its control of apoptosis in two locations within the SCE: the proximal hinge ectoderm and the ectoderm between the ventrolateral *Fgf8^+^* epithelia and the olfactory placode. With regard to the first, we posit that *Foxg1* is required to stabilize the patterning of the ectoderm of the Hinge region. Hinge-related epithelial signals act to positionally entrench the jaw articulation; they also instruct hinge-centered module identify in the mdBA1 and mxBA1 CNC. Instructing mdBA1 identity involves, among other things, the regulated secretion of Edn1 from the BA1 ectoderm at the distal margins of the hinge (353, 354). Edn1 executes pattering instructions here through binding its receptor, Ednra. Controlled positioning of Edn1 ligand is required for proper patterning as its receptor, Ednra, is ubiquitously expressed in the fungible population of the CNC. In the mdBA1 CNC, Edna-mediated signaling normally stimulates *Dlx5* expression which in turn induces *Dlx3* and establishes a molecular cascade yielding lower jaw morphogenesis. In reciprocal studies of the loss of *Dlx5/6*, wherein maxillary identity is molecularly and structurally acquired in the mdBA1 CNC (6, 33), forced expression of Edn1-Ednra signaling in the mxBA1 CNC leads to molecularly and structurally acquiring partial mandibular identity in the mxBA1 field. Edn1 signaling is apparently normally prevented from providing cues to the mxBA1 CNC in part by the specific ectodermal expression of *Six1* at the proximal extremity of the peri-cleftal portion of the hinge signaling center; in the absence *Six1*, *Edn1* is ectopically expressed proximally in the hinge-associated mxBA1, wherein it interacts with Edna, and hinge-centered mandibular structures subsequently begin to develop (354). Thus, the proximal hinge-centered ectoderm is a critical factor in the repression of mdBA1 identity in the mxBA1 CNC field. We have shown that loss of *Foxg1* leads to both significant, aberrant apoptosis in this proximal hinge region ectoderm at E9.25, and the subsequent ectopic induction of *Dlx5, Dlx3* and *Pitx1* in mxBA1 CNC. We suggest that loss of *Foxg1*-mediated cell vitality abrogates cues necessary for hinge-mediated suppression of Edn1 signaling in the mxBA1 CNC (a notion currently being further investigated).

A second notable early embryonic mis-regulation of apoptosis due to the loss of *Foxg1* occurs at E9.25 with the loss of the patch of apoptotic cells normally found clustered in the SCE between the olfactory placode and the cluster of *Fgf8^+^* cells normally found ventro-lateral to the ANR/CP. The specific functional consequence of programmed cell death here remains unclear though there are a number of plausible hypotheses. For instance, the focalization of apoptosis may be programmed so as to ensure a range limitation to *Fgf8*-mediated patterning cues within the SCE; or it may assist in the removal of cells in order to help establish a necessary and specific cell orientation and polarity within this region of the SCE. While specific function(s) remain to be determined, the absence of this focalization of apoptosis in the SCE of *Foxg1* null embryos coupled with the aberrant patterning of the regionally associated skeleton suggests the hypothesis that these bilateral apoptotic patches are required for the normal compartmentalization of the SCE, the subsequent elaboration of the λ-junction, and overall patterning of the upper jaws.

### Foxg1, Dlx5 and the facial midline

*Foxg1* is one of a number of transcription factors extensively expressed in the early embryonic SCE. To examine the broader context in which *Foxg1* regulates SCE development and craniofacial patterning, we examined the potential genetic interactions of *Foxg1* and *Dlx5*, which is also highly expressed throughout the early SCE (79). Demonstrative of a clear genetic interaction, we found that among the numerous perturbations of craniofacial development evinced by the *Foxg1^−/−^*;*Dlx5^−/−^* mutants there occurred a fundamental change in the character of the rostral midline. Compound *Foxg1^−/−^*;*Dlx5^−/−^* perinates altogether lacked frontonasal vibrissae, rhinaria, and external nares; rather, their frontonasal region was represented by three nearly identical raised tissue masses each remarkably containing within its own cartilaginous rod bound on both sides by dermatocranial bone that was, in turn, associated with a dysmorphic, diminished incisor. This suggests that in the compound mutants bilateral duplications of the midline had occurred. Examination of E10- E11 compound null mutants showed that together *Foxg1* and *Dlx5* are essential for the development in the SCE of olfactory pits and the λ-junction. These structural changes were accompanied by the complete absence at E10.25 of λ-junction associated *Bmp4* expression accompanied by an extension of expression at the midline associated with the CP. Notably, by E10.75 the CNC topographically subjacent to where each olfactory pit would normally have invaginated had proliferated enough to form a small mound of tissue the center of which contained focal clusters of *Fgf8* expressing cells. It is tempting to speculate each *Fgf8* expressing cluster is associated with one of the lateral cartilaginous rods that we take as duplicated nasal midline (TBP) structures. How then to explain the tripartite nature of the perinatal skull? These bilateral clusters of expression were accompanied by decreased *Fgf8* at the CP but a substantial bilateral increase in SCE expression running from the mxBA1 oral ectoderm to the dorsomedial midline: thus, one possible explanation is that this abnormal, strong band of *Fgf8* expressing cells that crosses the midline acts as an inductive signal to the subjacent CNC in a manner mirrored by the two lateral, focal clusters of cells. It is clear from our initial examination of compound *Foxg1^−/−^*;*Dlx5^−/−^* mutants that the ontogeny of the SCE in these and other mutants bears further investigation.

### *Foxg1* and the human condition

*FOXG1* involvement has been implicated in both oncogenesis and congenital human deficits (237). With regard to the latter, sufficient evidence of neurodevelopmental deficits stemming from *FOXG1* haploinsufficiency or duplication in humans has led to the recognition of ‘FOXG1-related disorders’ as a distinct clinical classification variously involving microcephaly, mental impairment, autism spectrum disorders, and a congenital variant of Rett (RTT) syndrome (FOXG1 syndrome (OMIM 613454); 355–369). RTT, the typical form of which has been liked to mutations involving MECP2, is one of the most common causes of intellectual disability in girls and is characterized by neurodevelopmental regression after the first year, including of acquired spoken language and hand skills, together with the onset of stereotypic hand movements and gait abnormalities. The FOXG1-related congenital variant of RTT appears earlier in development, usually by six months, and clinically presents a core set of neurodevelopment features, including microcephaly, developmental delay, severe cognitive disability, absence or minimal language development, early-onset dyskinesia and hyperkinetic movement, epilepsy, and cerebral malformation. Additionally, FOXG1 syndrome is typified by characteristic craniofacial features, including: a prominent metopic suture, large ears, bilateral epicanthic folds, a bulbous nasal tip, depressed nasal bridge, tented or thickened upper lip, prognathism, hypermetropia, and synophyrys (360). A significant linkage between the *FOXG1* locus at 14q12 and nonsyndromic orofacial clefting has also recently been reported (370). Notably, we have shown here that *Foxg1^−/−^* mutant cranial dysmorphologies align remarkably with this gestalt of defects, making *Foxg1* mutants ideal models to further investigate both the neurological and non-neurological aspects of the congenital variant of RTT, specifically, and craniogenetic development and evolution, generally.

## Materials and Methods

### Mice

Generation and genotyping of both the *Foxg1* (formerly *Brain Factor-1*) and *Dlx5* mice has previously been reported (130). Unless otherwise noted, multiple embryos or perinates were used for each experimental parameter.

### Detection of β-galactosidase activity

The *Foxg1* mutant alleles (or *Foxg1^LacZ/+^*) possess an in-frame β- galactosidase cassette that permits the detection of transcription of the endogenous *Foxg1* locus (130). *Foxg1^-/^and Foxg1^+/−^*littermate embryos were harvested from timed pregnancies, rinsed in phosphate-buffered saline (PBS), fixed (while on a rocker) in 4% paraformaldehyde (PFA) in phosphate-buffered saline (PBS) for 5-30 minutes (depending on stage), and then rinsed again in PBS. β-galactosidase activity was detected in prepared embryos through use of either the X-gal or Salmon-gal (Roche) substrates prepared as follows: 100 µl 1M TrisHCl pH=7.3; 100 µl of 0.005% Na-deoxycholate; 10 µl of 0.01 % Nonidet P40; 200 µl of 5mM K_4_Fe(CN)_6_; 200 µl of K_3_Fe(CN)_6_; 20 µl of 2mM MgCl_2_; 200 µl of 0.8 mg/ml X-Gal (or Salmon-gal) and 9.7 ml of distilled H_2_O. The reactions took place on a rocker in an incubator held at 37° C for 2 to 6 hours. The specimens were subsequently fixed overnight in 4% PFA at 4° C, photographed, and then prepared for histological examination.

### Murine anatomical analyses

*Foxg1* and compound *Foxg1;Dlx5* embryos and perinates were collected, rinsed in PBS, and the processed for either histological or whole anatomy hard tissue analysis. Differential staining of bone and cartilage through use of Alizarin red and Alcian blue was achieved following established protocols (133, 134). Skeletal preparations were photographed using a Leica MZFLIII microscope with a Leica DFC300FX camera and then stored in 100% glycerol at room temperature. Animals utilized for histological analysis were fixed in 4% PFA in PBS, embedded in paraffin or OCT (when previously stained with X-gal), and then cut in 8–20 µm serial sections. Sections were then stained with a modified Gimori trichrome (as per 133, 134).

### Whole mount *in situ* hybridization and immunohistochemical assays

Embryos were fixed overnight in 4% PFA in PBS at 4**°**C, rinsed, and then passed through a grades series of MeOH. Whole mount *in situ* hybridization and preparation of ribroprobes followed standard protocols (as described in 79). Unless otherwise noted, multiple embryos were used for each experimental parameter. Embryonic blood vessels were detected using rat α-endomucin antibodies (gift of C. Ruhrberg, University College London, UK).

### Whole mount TUNEL assay for Apoptosis

Whole mount apoptosis analysis was carried out using a modified protocol for the ApopTag Peroxidase *In Situ* Apoptosis Detection Kit (Chemicon international, S7100). Dehydrated embryos stored in MeOH were re-hydrated through a graded methanol series (100%, 75%, 50%, 30%) and PBS. They were then incubated in 20 µg/ml Proteinase K 20 in PBT (PBS and 0.1% Tween 20) for 15 minutes at room temperature without agitation and washed twice for 20 minutes in PBS followed by a 10 minutes incubation in 3% H_2_O_2_ in PBS and again washed in PBS (2 x 5 minutes). They were then placed in equilibration buffer for 5 minutes before being incubated in the digoxigenin labelled working strength TdT enzyme (1 ml of reaction buffer and 10 µl of TdT enzyme) for 1 hour at 37° C. Stop/Wash buffer was then applied, the samples were agitated for 10 seconds and then allowed to rest for 15 min at RT before being washed 3 times in MABT for 5 minutes each. The embryos were then incubated for 1 hour in blocking solution (MABT + 2% BBR + 20% heat-treated serum + 0.5 mg/ml levamisole) before being placed in blocking solution + anti-digoxigenin antibody (1:2000 dilution) and incubated overnight at 4° C on a rocker. The following day the samples were rinsed 3 times and then washed (6 x 1 hour) in MABT + 0.5 mg/ml levamisole at room temperature and then washed overnight in the same solution at 4° C. The following morning, the samples were again washed in fresh MABT + 0.5 mg/ml levamisole for 1 hour, then twice in NTMT + 0.5 mg/ml levamisole for 10 minutes before incubating in NTMT + 4.5 µl/ml NBT + 3.5 µl/ml BCIP + 0.5 mg/ml levamisole (protected from light) in order to allow the color reaction to proceed. Once suitably developed, the reaction was halted by washing 3 x 5 minutes in PBT. Samples were then photographed and stored in 4% PFA at 4**°** C.

### Scanning electron microscopy

Preparation of embryonic material for electron microscopy followed protocols (79). Briefly, embryos from timed pregnancies were harvested and fixed at 4**°**C overnight in 4% PFA and 0.2% glutaraldehyde, then washed in PBS, dehydrated in a graded ethanol series, critical point dried, mounted onto alluminium stubs, sputter coated with gold-palladium and subsequently examined and photographed with a FEI Quanta FEG scanning electron microscope operating at 10kV.

## Acknowledgements

We thank JLR Rubinstein and E Lai for the *Foxg1* (*BF1*) mutant line and P Crossley, G Martin, J Rubenstein, L Selleri, and P Sharpe for various riboprobe plasmids, as well as Q Schwarz and C Ruhrberg for the anti-endomucin antibody. MJD was funded by the Royal Society (UK), the Dental Institute of King’s College London, Friends of Guy’s Hospital and the Charite Universitätsmedizin Berlin. CC was funded by a Marie Curie Early Training Fellowships (MEST-CT-2004-504025).

## Competing Interests

We have no competing interests to report.

## Figure Legends

**Supplementary Figure 1.**
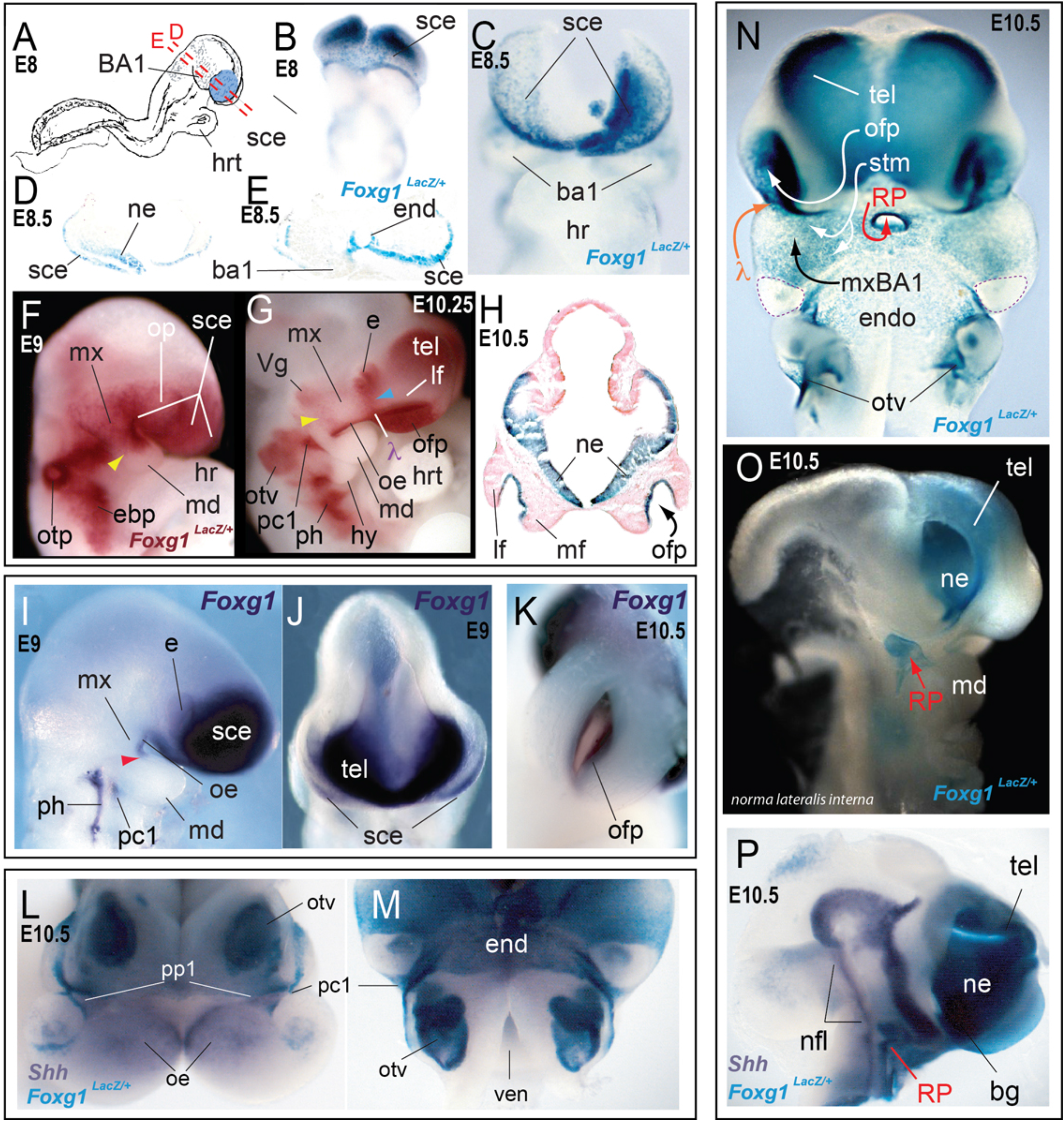
Early ontogeny of *Foxg1* expression in the craniogenic primordia demonstrating the extensive expression in the surface cephalic ectoderm. <u>(**A**)</u> Diagram of E8 murine embryo indicating the position of the sce and planes of section for D and E. (B-H) β-galactosidase enzymatic activity in the SCE of *Foxg1^LacZ/+^* embryos at E8 (B), E8.5 (C-E), E9 (F), E10.25 (G) and E10.5 (sectioned, H). (I-K) *In situ* hybridization of Foxg1 at E9 (I, J) and E10.5 (K). (L-P) β-galactosidase enzymatic activity in the cranial primordia of *Foxg1^LacZ/+^* embryos at E10.5. **<u>Abbreviations</u>**: ***ba1***, first branchial arch; ***bg***, basal ganglia primordia; ***e***, eye primordia; ***ebp***, epibranchial cells; ***end***, endoderm; ***hrt***, heart; ***hy***, hyoid arch; ***lf***, lateral frontonasal process; ***md***, mandibular branch of the first branchial arch; ***mf***, medial frontonasal process; ***mx***, maxillary branch of the first branchial arch; ***ne***, neuroectoderm; ***nfl***, neural tube floor; ***oe***, oral epithelium; ***ofp***, olfactory pit; ***op***, optic primordia; ***otp***, otic pit; ***otv***, otic vesicle; ***pc1***; first pharyngeal cleft; ***pp1***, first pharyngeal plate; ***RP***, Rathke’s pouch; ***sce***, surface cephalic ectoderm; ***stm***, stomodeum; ***tel***, telencephalon; ***ven***, ventricle **λ**, center of the lambdoidal junction.

**Supplementary Figure 2.**
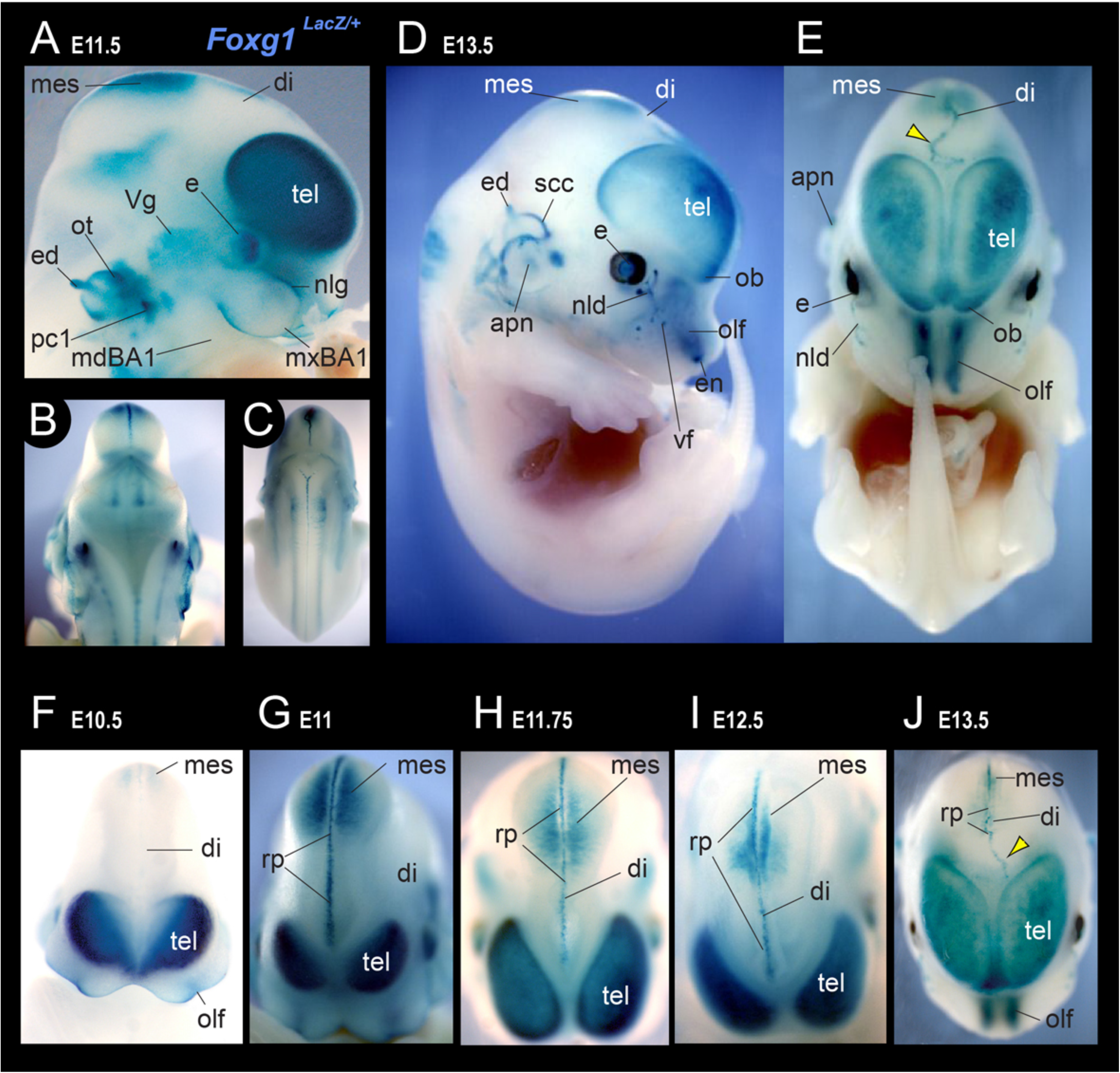
Mid-fetal ontogeny of Foxg1 expression in the craniogenic primordia demonstrating the extent of *Foxg1* expression previously uncharacterized. <u>(**A-J**)</u> β-galactosidase enzymatic activity in the embryonic cranial primordia of *Foxg1^LacZ/+^* embryos at E11.5 (A-C), E13.5 (D,E), E10.5 (F), E11 (G), E11.75 (H), E12.5 (I) and E13.5 (J). **<u>Abbreviations</u>**: ***apn***, auricular pinnae; **di**, diencephalon; ***e***, eye primordia; ***ebp***, epibranchial cells; ***ed***, endolymphatic duct; ***en***, external nares; ***mdBA1***, mandibular branch of the first branchial arch; ***mes***, mesencephalon; ***mxBA1***, maxillary branch of the first branchial arch; ***nld***, nasolacrimal duct; ***nlg***, nasolacrimal groove; ***olf***, olfactory primordia; ***ob***, olfactory bulb primordia; ***ot***, otic vesicle; ***pc1***; first pharyngeal cleft; ***rp***, roof plate; ***scc***, semicircular canal; ***tel***, telencephalon; ***Vg***, trigeminal ganglion; ***vf***, vibrissae follicle.

**Supplementary Figure 3.**
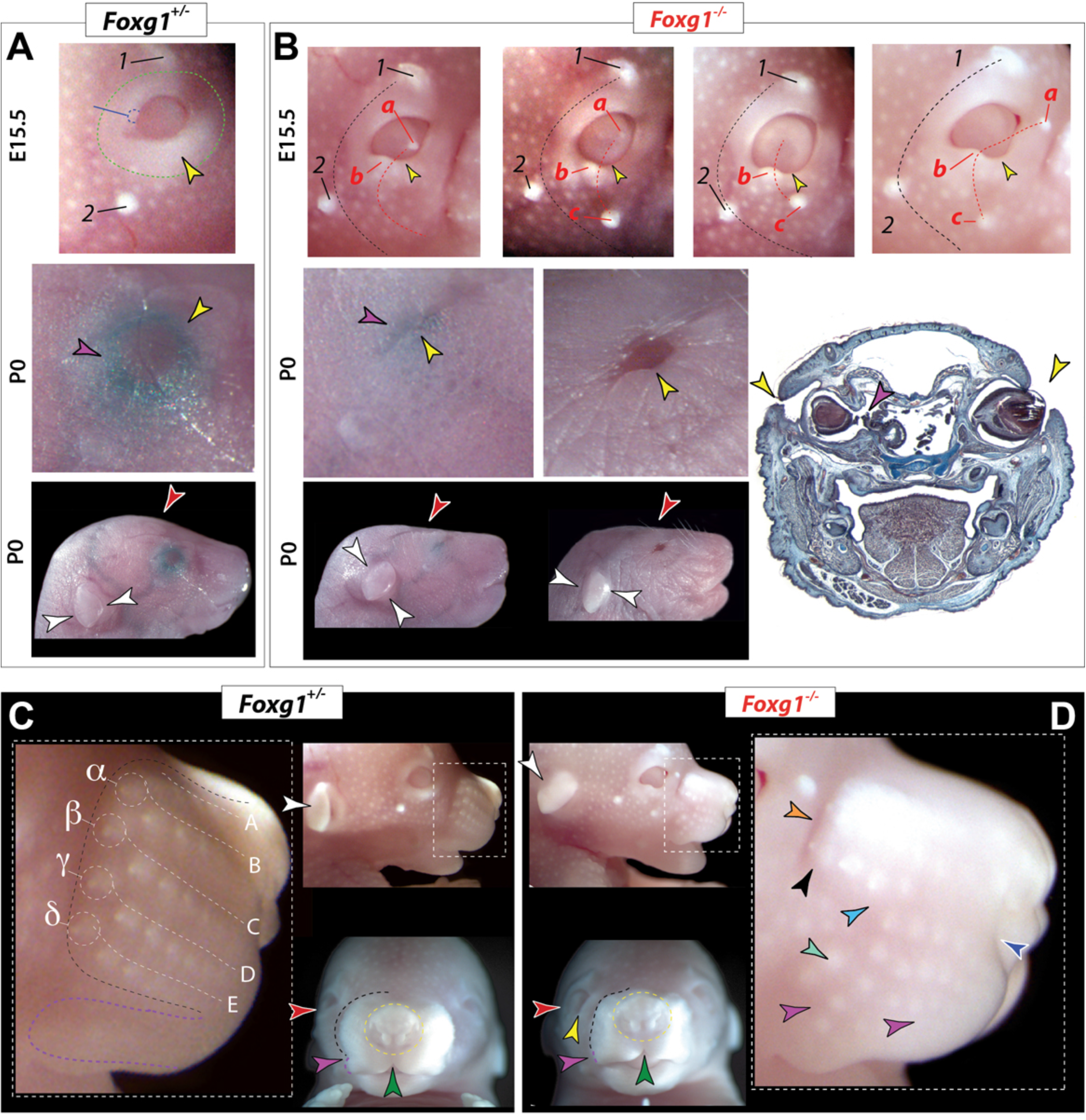
Deficits of the ectodermal derivatives, including of the auxiliary eye structures, vibrissae, and upper lip in Foxg1^−/−^ mutants. Development of the eyes and eyelids of E15.5 and P0 *Foxg1^+/−^* (A, C) and *Foxg1^−/−^* littermates (B, D). (**A, B**) Ocular and eyelid defects in mutants. ***1***, supra-orbital tubercle; ***2***, post-orbital tubercle; ***a, b, c*** ectopic sinus tubercles. Yellow arrowhead, lower eyelid; blue arrowheads, eyes open phenotype; red and white arrowheads, highlight the flattened crania of the mutants; black and white arrowheads, the positioning of the pinnae. (**C, D**) Defects of vibissae rows, upper lip and nasal development at E15.5. A-E, α-δ, vibrissae tubercles. black and white arrowheads, the positioning of the pinnae. Black dashed lines, the outline of the vibrissae pad and upper lip; yellow dashed lines, outline the rhinarium.

**Supplementary Figure 4.**
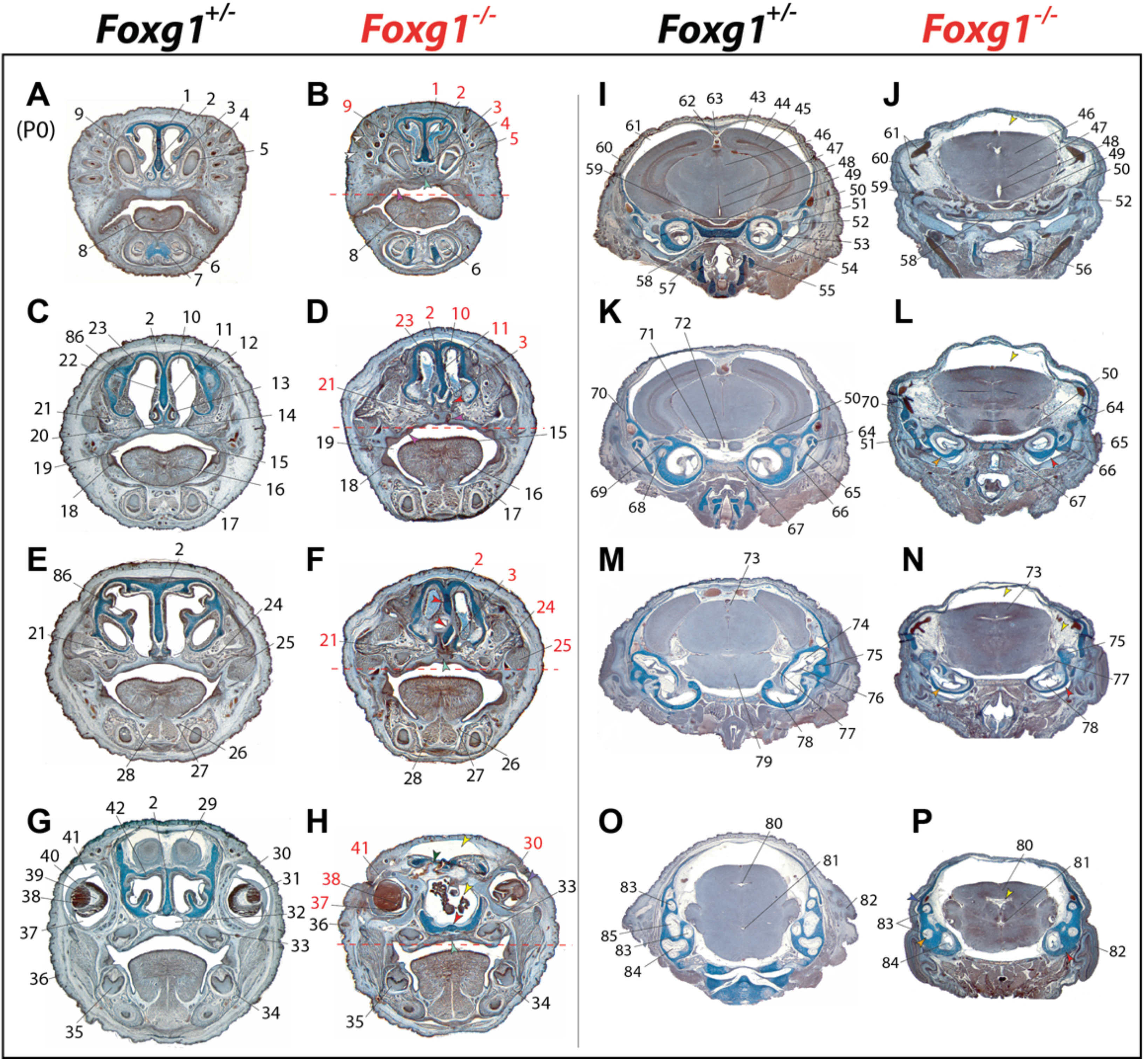
Skeletal and soft tissue defects of the crania of Foxg1^+/−^ and Foxg1^−/−^ *P0 mutants.* Gimori trichrome stained histological sections of *Foxg1^+/−^* (**A, C, E, G, I, K, M, O**) and *Foxg1^−/−^* (**B, D, F, H, J, L, N, P**) mutants. **1**: nasal capsule; **2**: nasal septum; **3**: paraseptal cartilages; **4**: longitudinal section of follicle of vibrissa; **5**: primordium of left upper incisor tooth; **6**: primordium of left lower incisor tooth; **7**: site of fusion of the right and left Meckel’s cartilage; **8**: tongue; **9**: ossification in membranous primordium of premaxillary; **10**: nasal cavity; **11**: tubules of serous glands associated with nasal septum; **12**: cartilage of pars intermedia; **13**: vomeronasal organ (Jacobson’s organ); **14**: anterior extremity of secondary palate (palatal shelf of maxilla) ; **15**: dorsal surface of tongue; **16**: intrinsic muscle of the tongue; **17**: ossification in body of the dentary; **18**: mucous membrane of oral cavity; **19**: oral cavity; **20**: ossification in inferomedial part of nasal septum; **21**: palatal process of maxilla; **22**: cartilage primordium of nasal septum; **23**: olfactory epithelium; **24**: ossification in maxilla; **25**: anterior fibres of masseter muscle; **26**: sublingual duct; **27**: submandibular duct; **28**: genioglossus muscle; **29**: olfactory bulb; **30**: line of fusion between the upper and lower eyelids; **31**: lens; **32**: naso-paharynx; **33**: primordium of upper molar tooth; **34**: ossification of dentary; **35**: primordium of lower molar tooth; **36**: hair follicle in skin; **37**: intra-retinal space; **38**: neural layer of retina; **39**: hyaloid cavity; **40**: cornea of eye ; **41**: conjuctival sac; **42**: cartilage primordium of cribriform plate of ethmoid bone; **43**: cortical plate of neopallial cortex; **44**: intermediate zone; **45**: ependymal zone; **46**: diencephalon (thalamus); **47**: diencephalon (hypothalamus); **48**: third ventricle; **49**: infundibulum of pituitary; **50**: trigeminal (V) ganglion; **51**: facial (VII) ganglion; **52**: Meckel’s cartilage; **53**: cartilage primordium of petrous part of temporal bone; **54**: tubo-tympanic recess; **55**: posterior part of pharynx; **56**: entrance from pharynx into the esophagus; **57**: fibres of superior and middle constrictor muscle; **58**: parachordal basal plate; **59**: residual lumen of Rathke’s pouch; **60**: subarachnoid space; **61**: vascular elements of pia-arachnoid; **62**: falx cerebri; **63**: superior sagittal dural venous; **64**: cartilage primordium of head of the malleus; **65**: cartilage primordium of body of the incus; **66**: cartilage primordium of crus longum of the incus; **67**: ossification in central part of cartilage primordium of basioccipital bone ; **68**: lateral semicircular canal; **69**: vestibular (VIII) ganglion; **70**: cartilage primordium of squamous part of temporal bone; **71**: floor of pons; **71**: basilar artery; **72**: roof (tectum) of midbrain; **73**: cochlear duct; **74**: superior semicircular canal; **75**: spiral organ of Corti; **76**: vestibular (VII) ganglion; **77**: cochlea; **78**: pons; **79**: mesencephalic vesicle; **80**: fourth ventricle; **81**: cortical plate of neopallial cortex; **82**: ear pinna; **83**: anterior semicircular canal; **84**: posterior semicircular canal; **85**: lateral semicircular canal.

**Supplementary Figure 5.**
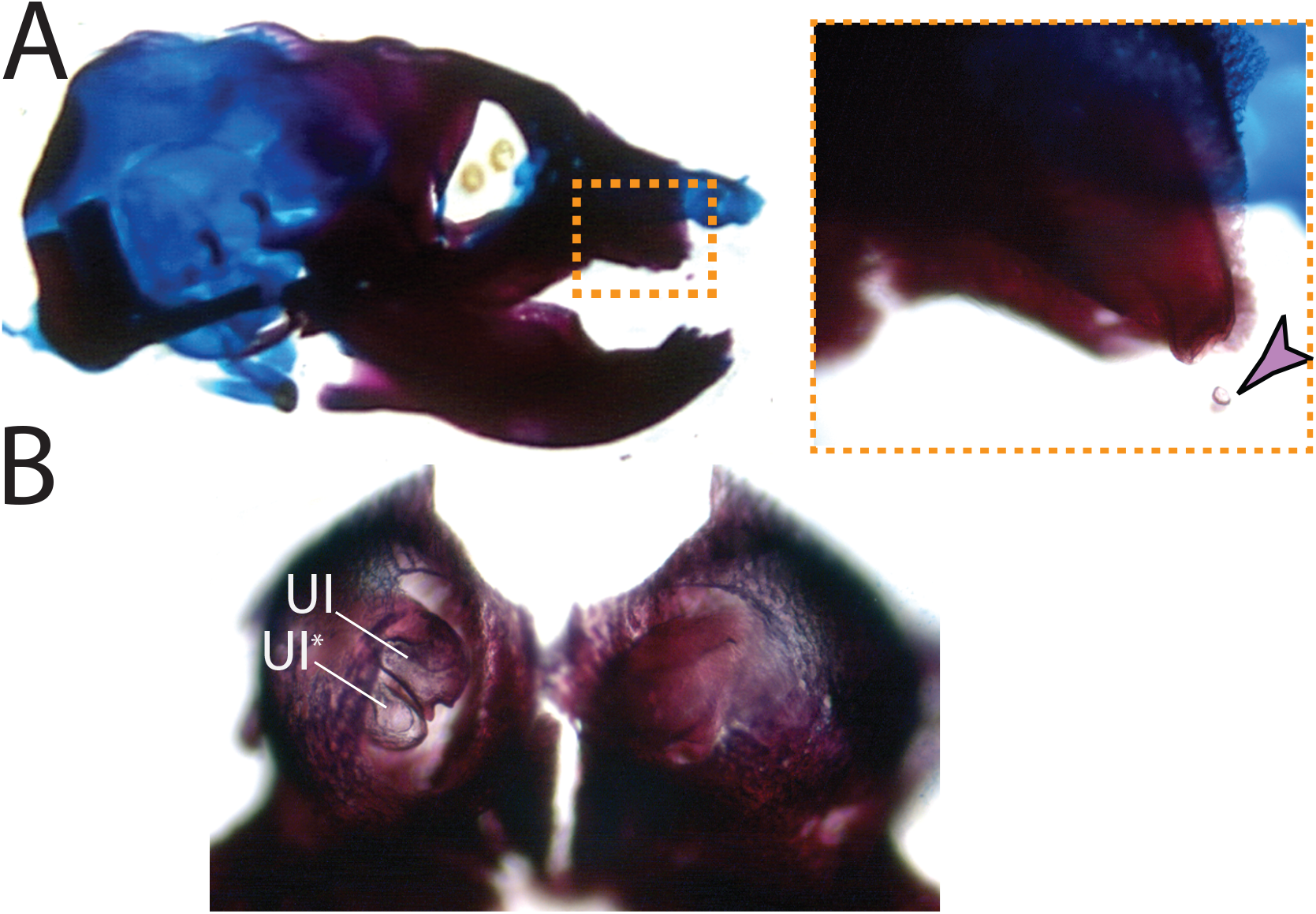
Upper incisor development in Foxg1^−/−^ *P0 mutants*. (**A**) Small mineralized nodules associated with mutant upper incisors. (**B**) Duplicated incisor. UI, upper incisor.

